# Novel Methylation Markers in a Prostate Cancer Cohort are Associated with Disease Development and Relapse

**DOI:** 10.64898/2026.08.26.747370

**Authors:** Radoslaw P. Lach, Sara Pita, Wing-Kit Leung, Anne Babbage, Susan Merson, Steve Hawkins, Hayley Luxton, Jonathan Kay, Hayley C. Whitaker, Dan J. Woodcock, Valeriia Haberland, Zsofia Kote-Jarai, Toby Milne-Clark, Katherine O’Neill, Timothy Brendler-Spaeth, Melissa Cheung, Matthew Ko, CRUK ICGC Prostate Cancer Group, Harveer Dev, Adam Butler, Adam Lambert, Freddie C. Hamdy, Clare Verrill, Sally Field, G. Steve Bova, Christopher Foster, David E. Neal, David C. Wedge, Vincent J. Gnanapragasam, Anne Y. Warren, Ros A. Eeles, Colin S. Cooper, Daniel S. Brewer, Charlie E. Massie, Andy G. Lynch

## Abstract

Prostate cancer remains one of the most common cancers among men globally. While significant strides have been made in diagnosis and treatment, understanding the complex genetic and epigenetic underpinnings of the disease remains crucial for guiding intervention and developing more personalized and effective therapies. The importance of DNA methylation in prostate cancer has been known for some time, but important facets of the modulation of the epigenome during carcinogenesis remain obscure, partly because the bulk of cancer methylation data have been produced using microarray technologies.

Here we utilise the TruSeq methyl capture method (EPICseq) to profile the, previously defined, UK Prostate ICGC cohort of well-annotated primary prostate cancers. To this we add methylation sequencing of benign tissue from the same men. These data allow us to identify differentially methylated regions distinguishing cancerous and non-cancerous prostate tissue, while identifying numerous genes whose methylation profiles can perform that task as well as distinguishing between classes of prostate cancer. We describe a describe a methylation-based control mechanism for prostate-cancer-associated SNPs, and show that this seems a likely mechanism of action for a SNP near the MMP7 gene.

We describe three novel molecular signatures that arise from different aspects of the biology of prostate cancer revealed by sequencing. Each is shown to be an independent classifier of cancers into groups with different expected times to relapse. These consist of patterns in driver gene methylation, strand-specific methylation, and signal arising in mitochondrial reads.

We show that these signatures, combined with existing molecular tools, provide a powerful predictor of time to recurrence. By substantially enhancing understanding of prostate cancer risk, detection, and prognosis, we pave the way for the development of clinical practices that will benefit patients and improve outcomes.

## Introduction

Genomic analyses of prostate cancer have revealed important insights into mechanisms of cancer evolution [1–3], exemplified by the long tail of infrequently mutated driver genes in this disease[4–6]. Our previous papers as part of the International Cancer Genome Consortium (ICGC) UK Prostate group on prostate cancers from 205 patients also identified new driver genes and druggable targets, novel trajectories of metastatic spread, as well as two distinct evolutionary pathways in the development of primary disease termed the Alternative and Canonical evotypes. [1, 6, 7] While somatic alterations including point mutations, gene fusions and large deletions have been extensively studied, by far the most consistent molecular alterations observed in primary prostate cancer are DNA methylation changes[8–10].

Methylation array studies have provided evidence for distinct molecular subtypes of prostate cancer[4], and in a recent case showing that molecular subtypes can be linked to differences in survival[11]. Studies have identified potential prognostic markers (notably methylation of GSTP1)[12, 13], and have generated a first map of epigenetic phylogenies in prostate tumours[9, 14]. Multi-region analysis of prostate tumours has revealed common epigenetic changes that may precede the first genetic alterations[14], with hundreds of DNA methylation alterations commonly shared between genetically distinct tumour foci[9]. The results are consistent with an epigenetic ‘field effect’ that may contribute to the high rates of tumour formation in the prostate, multi-focal disease, and extensive genomic heterogeneity in localized prostate tumours[1–3]. It is also suspected that there is a close link between the metabolic condition in the prostate, and patterns of DNA methylation [10]. Our prior review of 18 studies found that more than 5,000 genes have altered methylation in cancerous prostate tissue, but fewer than 1,000 of these were identified in more than one study, suggestive of a large pool of changes across the genome, from which each study identified and reported a subset. [10]. Taken together, there is strong evidence supporting multiple DNA methylation changes as early, recurrent events in prostate tumorigenesis[12, 13, 15, 16].

Multiple expression-based markers have been reported for prostate cancer [17] including DECIPHER [18], Prolaris [19], and DESNT categories [20]. Genome-based biomarkers have also been reported ranging from individual genes and loci [21, 22] to distinct evolutionary pathways [7]. While a few DNA methylation markers have been reported as indicating outcome, including time to recurrence (as in those described more than a decade ago [23, 24]), these have been rarer than those based on gene expression.

Methylation studies have suffered from being predominantly microarray-based while genomic and expression-based studies have now largely switched to the use of sequencing technologies. Whole-genome sequencing, while the gold standard, is an expensive approach for methylation analysis, with excessive irrelevant data being generated, for example, for CpG-sparse regions. Sequence capture approaches offer a compromise between those two extremes[25].

We employed the Illumina TruSeq methyl capture method (EPICseq)[26], which covers over 3.3 million CpGs (10.7% of CpGs) [25], including regulatory elements and Infinium array targets, from 107Mbp of the human genome. EPICseq performed well in comparison with other targeted and reduced representation bisulfite sequencing approaches in a recent review, with more CpGs being investigated than all but the RRBS methods, high hybridization efficiency and strong reproducibility of coverage between runs [25]. Here, by applying EPICseq to our ICGC cohort of cancers and adjacent morphologically normal prostate tissues, we identify new signatures with diagnostic or prognostic value and provide new insights into prostate cancer biology not visible with microarray platforms.

## Results

### The Cohort

We present EPICseq data for 188 men from the Prostate ICGC UK cohort of patients with linked clinical data. We report also two previous molecular classifications that we have published. The DESNT classification is based on expression data [20] and includes a poor-prognosis class that we refer to hereafter as “DESNT tumours”. Evotypes, derived from genome sequencing data, classify tumours into two distinct evolutionary paths [7], of which the “alternative evotype” class was seen to have poorer survival.

For 172 men we have EPICseq data from both tumour and benign tissue obtained from prostatectomy specimens (“benign” here we use to mean tissue assessed by pathologists as non-cancerous). In addition, data from ten tumour samples with no paired benign sample, and from six benign prostate samples with no paired tumour sample, complete the cohort. [Table 1, Supplementary Figure S1, Supplementary Table S1]. We obtained an average of 55 million reads per sample, with two million CpGs reaching at least 10x sequencing coverage. [Supplementary Table S2]

**Table 1.** Basic clinical and demographic descriptors of the cohort.

|  |  | number (unless specified) | percentage |
| --- | --- | --- | --- |
| **Samples after QC** | Tumour Benign Pairs | 172 |  |
|  | Tumour Only | 10 |  |
|  | Benign Only | 6 |  |
| **Follow up** | Median | 2057 days |  |
| **Age at prostatectomy** | 40-49 years | 8 | 4% |
|  | 50-59 years | 61 | 32% |
|  | 60-69 years | 100 | 53% |
|  | 70-79 years | 19 | 10% |
| **Grade Group** | 1 | 11 | 6% |
|  | 2 | 117 | 62% |
|  | 3 | 44 | 23% |
|  | 4 | 7 | 4% |
|  | 5 | 9 | 5% |
| **DESNT** | Low risk | 145 | 77% |
|  | High risk | 37 | 20% |
|  | Unclassified | 6 | 3% |
| **Evotype** | Alternative | 28 | 15% |
|  | Canonical | 113 | 60% |
|  | Unclassified | 47 | 25% |

### Methylation profiles of driver genes predict progression

Across the captured regions, CpG methylation levels were higher in tumour samples (mean 45%) than in benign prostate samples (mean 43%) (Wilcoxon signed rank test, *p <* 0.0001) (Supplementary Table S2). The differences (tumour subtract benign) of general methylation levels were higher in lower grade group tumours than higher (Kruskal-Wallis, *p* = 0.0139), higher in Canonical-evotype than Alternative (Wilcoxon rank sum test, *p <* 0.0001), and higher in men with a *TMPRSS2-ERG* fusion than men without (Wilcoxon rank sum test, *p* = 0.0019) (Supplementary Figure S2A).

We can consider patterns of cytosine methylation in contexts other than CpG (e.g. a C followed by a non-G base followed by a G “CHG”, or a C followed by two non-G bases “CHH”). In these contexts, we see similar but muted patterns (Supplementary Figure S2B, S2C). EPICseq-wide, methylation differences (tumour subtract benign) were greater in men with a TMPRSS2-ERG fusion (Wilcoxon rank sum test, *p <* 0.0001 CHG and CHH), in high-risk DESNT groups (Wilcoxon rank sum test, CHG *p* = 0.0083, CHH *p* = 0.0021), and in the Canonical-evotype (Wilcoxon rank sum test, CHG *p* = 0.0014, CHH *p* = 0.0007). Overall, this manifests as a statistically significant, but small, average increase in the median methylation in CHG (median difference 0.01%, *p* = 0.0032) and CHH (median difference 0.01%, *p* = 0.0004) contexts for the regions captured (Supplementary Figure S2).

We compiled a list of prostate cancer driver genes through literature review. Variation of methylation in these genes [Supplementary Tables S3, S4] is greater in tumour samples than benign samples (115 genes out of 150, sign test, *p <* 0.0001), where there is notable homogeneity (Figure 1). Clustering of tumour profiles across the thirty-three most variable driver genes reveals two broad groups of tumours. These show a difference in relapse-free survival following surgery (Supplementary Figure S3; *p* = 0.018; Log-rank test) with the ‘right-hand’ high-risk cluster being enriched for alternative Evotype tumours, DESNT tumours and high-grade tumours (Supplementary Figure S3).

**Figure 1.**
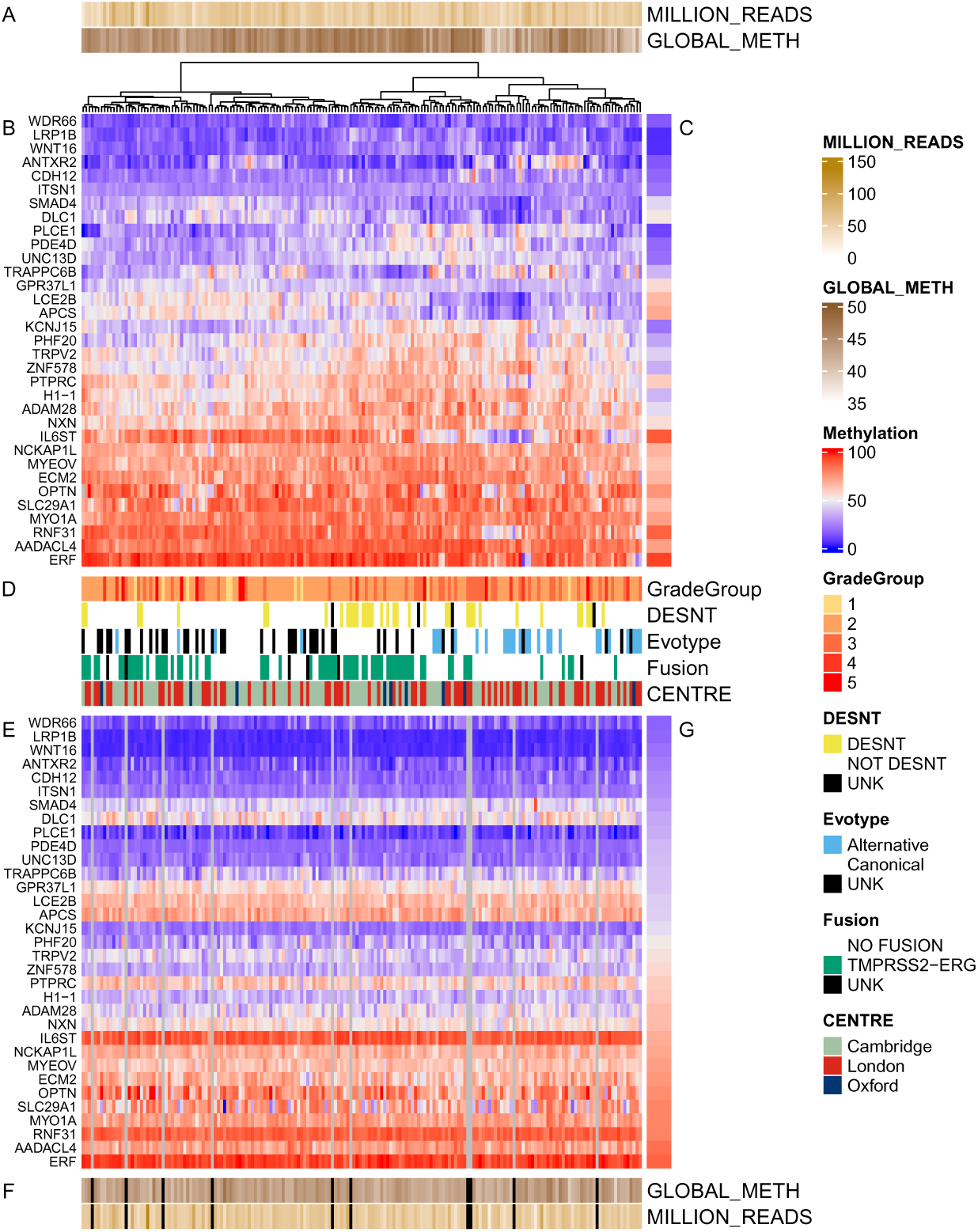
Methylation of prostate cancer driver genes in the cohort. From 158 previously identified prostate cancer driver genes, the 33 most variable (mad*>*5) across the combined tumour and benign samples are shown. A) For 182 primary tumour samples illustrated, the number of sequencing reads and the mean methylation levels of observed CpGs are shown. B) A heatmap of the 33 driver genes for those 182 tumour samples. Samples are clustered by Euclidean distance and Ward D2 agglomeration. Genes are ordered by mean methylation value in these samples. C) The mean values for the 33 driver genes from 172 matched benign samples are shown in the margin for easy comparison. D) Important demographic and clinically relevant characteristics of the cohort are given. These consist of the Grade Group, whether men are in the high risk DESNT category, which Evotype category they are in, whether they have a TMPRSS2-ERG fusion [defined by RNA-seq], and from which centre the patient was recruited. E) A heatmap of methylation levels for the driver genes in matched benign samples. Columns and rows are ordered to match the upper heatmap for easy comparison. Note that only 172 of the cases had benign samples that passed quality assurance. F) For the 172 matched benign samples are illustrated the number of sequencing reads and the mean methylation levels of observed CpGs. G) The mean values for the 33 driver genes from matched tumour samples are shown in the margin for easy comparison.

This high-risk cluster divides into two sub-clusters, one enriched for Alternative-evotype tumours (Fisher’s exact test; *p* = 0.0002) and one enriched for DESNT tumours (*p <* 0.0001) (Figure 1, Supplementary Figure S4). The average driver-gene profiles for these two enriched sub-clusters highlight four discordant genes (*ANTXR2, TRAPPC6B, IL6ST* and *LCE2B*) (Supplementary Figure S4). Just two of those genes (*TRAPPC6B* and *LCE2B*) show some ability to distinguish DESNT and Alternative-evotype individuals, correctly classifying 55/64 men (number expected by chance = 33, Cohen’s Kappa = 0.7).

### Methylation changes from benign tissue to tumour are widespread

Consistent with anticipated wide-scale alterations in the epigenome, we identified 202,775 differentially methylated cytosines (DMCs) between 182 tumour and 178 benign samples (Supplementary Table S5). These were recurrent events, with 50% of DMCs occurring in at least 74% of patients (Supplementary Figure S5). Of these, 148,475 were hypermethylated in the tumour (*q <* 0.01, methylation difference *>* 22%; Figure 2). Through merging DMCs, we identified 32,873 differentially methylated regions (DMRs) recurrent in at least 50% of patients (see Supplementary Methods and online code for details, Supplementary Table S6).

**Figure 2.**
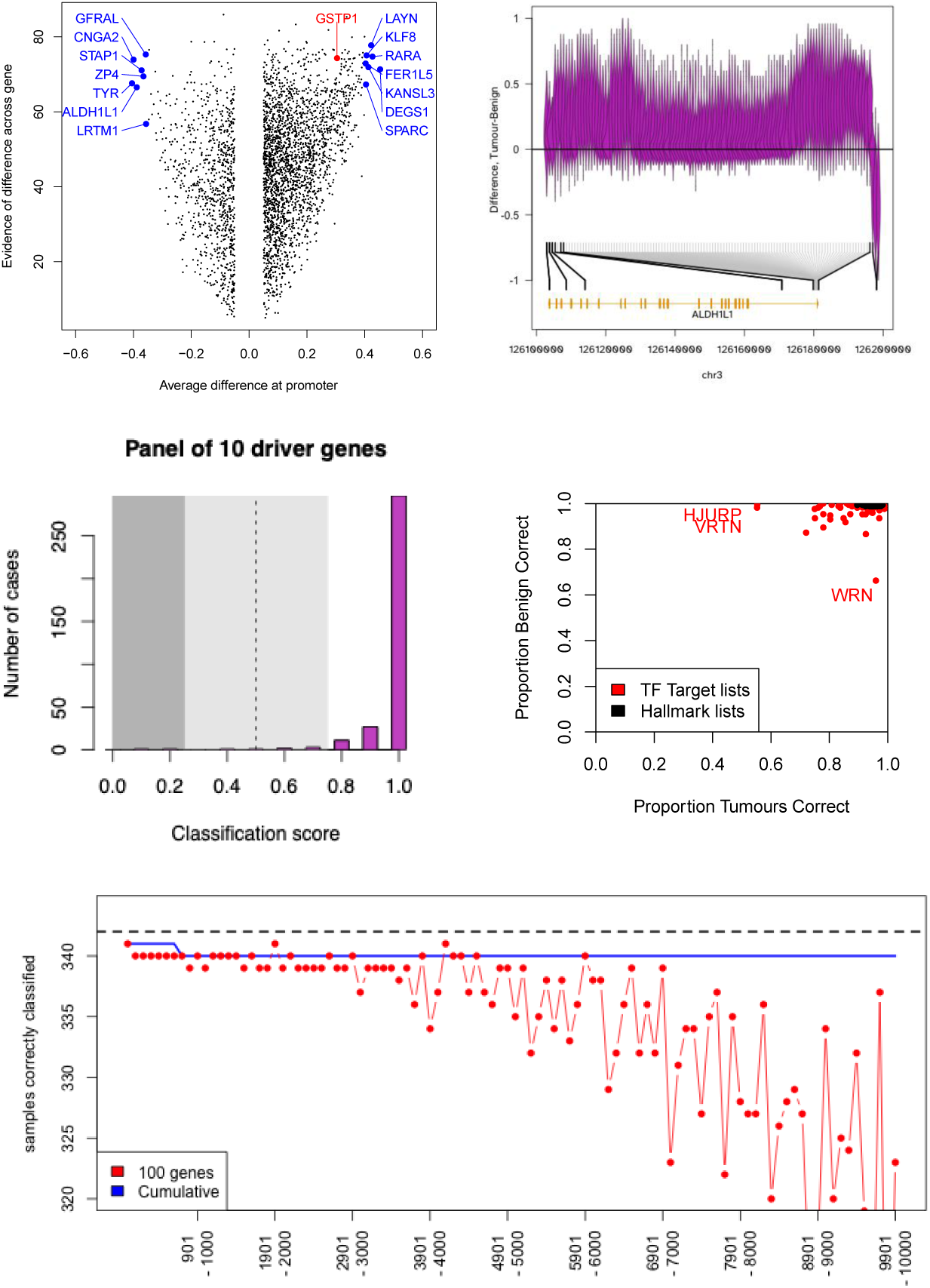
Genes that distinguish Tumour and Benign tissue. A) Plot of evidence of ability of genes to distinguish tumour samples from benign (*−log*_10_(*p*)) against difference in proportion of methylation across the promoter region. Demonstrating that there are informative genes with little change at the promoter, and genes where promoter methylation is higher in benign samples. B) An example of a gene (ALDH1L1, also highlighted in panel A) where a region in the promoter region highly methylated in benign tissue, but the gene body shows higher methylation in tumour samples. C) Giving the distribution of classification scores based on the consensus of an 10 gene panel. A value of 1 meaning that the sample is correctly classified by all 10 genes. D) Highlighting the ability of gene sets to classify Benign and Tumour samples. Transcription factor target lists and Cancer Hallmark lists were obtained from the Human MSigDB Collections (www.gsea-msigdb.org/). Samples were classified based on majority voting amongst all genes in the gene list. E) For genes ordered by their individual predictive ability, the performance in classifying 344 samples for the complete list of genes down to the gene indicated on the x axis, and for 100 genes around the gene indicated on the x axis. Given the nature of three samples [see supplementary methods] the maximum performance expected is 342/344 and this is indicated by a horizontal line.

From the observed differentially methylated regions (DMRs) (Supplementary Figure S5A), the average patient sees differential methylation at 84.7% of DMRs (Supplementary Figure S5B), and 24, 613 DMRs are present in more than 75% of patients (Supplementary Figure S5C). Methylation patterns in the DMRs define a subset of patients demonstrating increased risk of recurrence (HR = 3.36, p-value = 7.73e-05, Supplementary Figure S5D).

The genome-wide changes in the methylome, associated with prostate cancer, mean that the majority of genes (13832 genes when controlling FDR at 1%) have profiles that show some association with tumour and benign status. Focussing on 172 tumour/benign pairs of samples, we identify 679 genes that individually through unsupervised clustering of methylation profiles can categorize tumour and benign tissue with at most 10 misclassifications across the set; performance comparable to GSTP1 (Supplementary Table S7). This despite some samples defying classification – see methods. We see a trend that the fewer misclassifications there are in our data, the more likely it is that the gene will have been identified in previous surveys [10] [Supplementary Figure S6A], but 401 of those 679 genes were not previously identified in those studies. Other predictors of whether the gene has been previously identified are the size of the difference between tumour and benign methylation levels, and the number of discriminating CpGs immediately upstream of the gene (Supplementary Figure S6C).

Naturally, taking the consensus of a panel of genes can provide stronger performance in classification. For example, a panel of 10 prostate cancer driver genes, not identified in our previous survey of 18 studies investigating differentially methylated genes (*CASZ1*, *FOXP1*, *ROBO2*, *ZBTB20*, *CDH12*, *DSE*, *CHD7*, *CFAP251*, *ZFHX3*, *SMAD7*), correctly classifies 340/344 samples, with one ambiguous classification [Figure 2c]. Hallmark gene sets and transcription factor targets obtained from the Molecular Signatures Database [27] also provide excellent classification [Figure 2d], but in fact any large enough set of genes will match this performance: A set of 100 genes can provide excellent classification even if individually none are in the top 5000 discriminatory genes [Figure 2e].

### Gene-level summaries obscure tumour sub-group-specific details

While summaries of gene-body methylation or gene-promoter methylation are convenient to work with, the resolution of our data allows us to see localized patterns that distinguish not only tumour from benign tissue, but also sub-categories of tumour. We anticipate tens of thousands of loci differing with *TMPRSS2-ERG* fusion status [28, 29], but defining five categories based on *TMPRSS2-ERG* fusion, DESNT status and evotype (Supplementary Table S9), we see that the changes are more subtle.

For example, in SPOP (Figure 3) – a key Prostate cancer gene that is frequently mutated or downregulated [30] – we see a region with twelve CpGs that can distinguish the five tumour categories. One might conclude that ERG+ men had low methylation and ERG-men had high methylation akin to benign tissue in that region, but men in the DESNT risk group have lower methylation also, an effect that may be additive to that of the fusion. Elsewhere in the gene, there is a region where all of the tumour categories exhibit high methylation, distinct from the low methylation seen in benign tissue.

**Figure 3.**
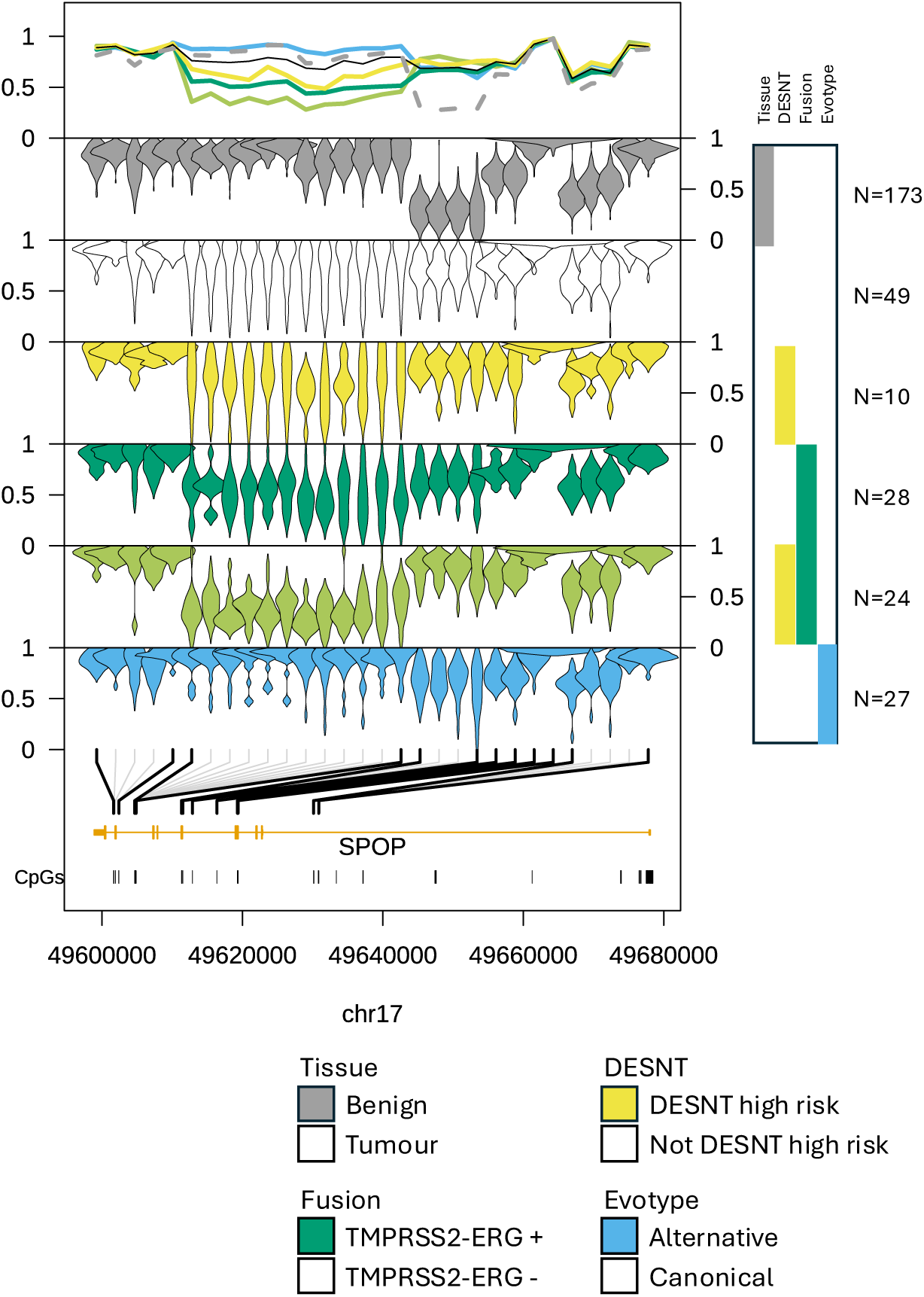
Illustrating, for selected CpGs in *SPOP*, the observed methylation levels in each of the five groups of tumours. Bottom we see the gene, coordinates, and locations of CpGs. Then, moving up, we see violin plots for selected CpGs for the alternative Evotype samples, for DESNT high-risk samples with a detected *TMPRSS2-ERG* fusion, for other samples with a detected *TMPRSS2-ERG* fusion, for other high-risk DESNT samples, and for tumour samples with none of the three characteristics (high-risk DESNT, alternative Evotype, or fusion). Profiles for Benign samples are shown at the top for comparison, and then we see mean values for the five tumour categories and Benign tissue in the top section of the plot. For the plots of the mean profiles, the tumour samples with none of the three characteristics are shown with a narrow black like, while the benign samples have a dashed line. For the violin plots, and mean methylation plot, the CpGs have been equally spaced. Below the first of the violin plots lines indicate the loci in the gene at which the CpGs are situated.

Other genes show different patterns [Supplementary Table S9] and a selection are shown in Supplementary Figures S7 to S21. These genes were chosen as they highlight differences between the five categories, and demonstrate the risks of misinterpreting a gene-average methylation summary score, or misinterpreting the results of a tumour-benign differential analyses when the cohort make-up is not considered. These effects, coupled with low numbers in some grade groups in our cohort, make it unlikely that we would see trends with Grade groups (Supplementary Figures S22 to S36, but a trend is apparent for *CDH4* (Supplementary Figure S37).

### DNA Methylation reveals a mechanism of action for risk-associated germline variants

One of the first genomic variants to be associated with prostate cancer risk was rs11568818 in the region of *MMP7* [31]. It is a known eQTL in prostate tissue [32], with unclear mechanism of action. We note that the risk allele (A/T) ‘removes’ a CpG from the promoter region of *MMP7* [Figure 4A], and we hypothesize that this may impact on methylation patterns in the region.

**Figure 4.**
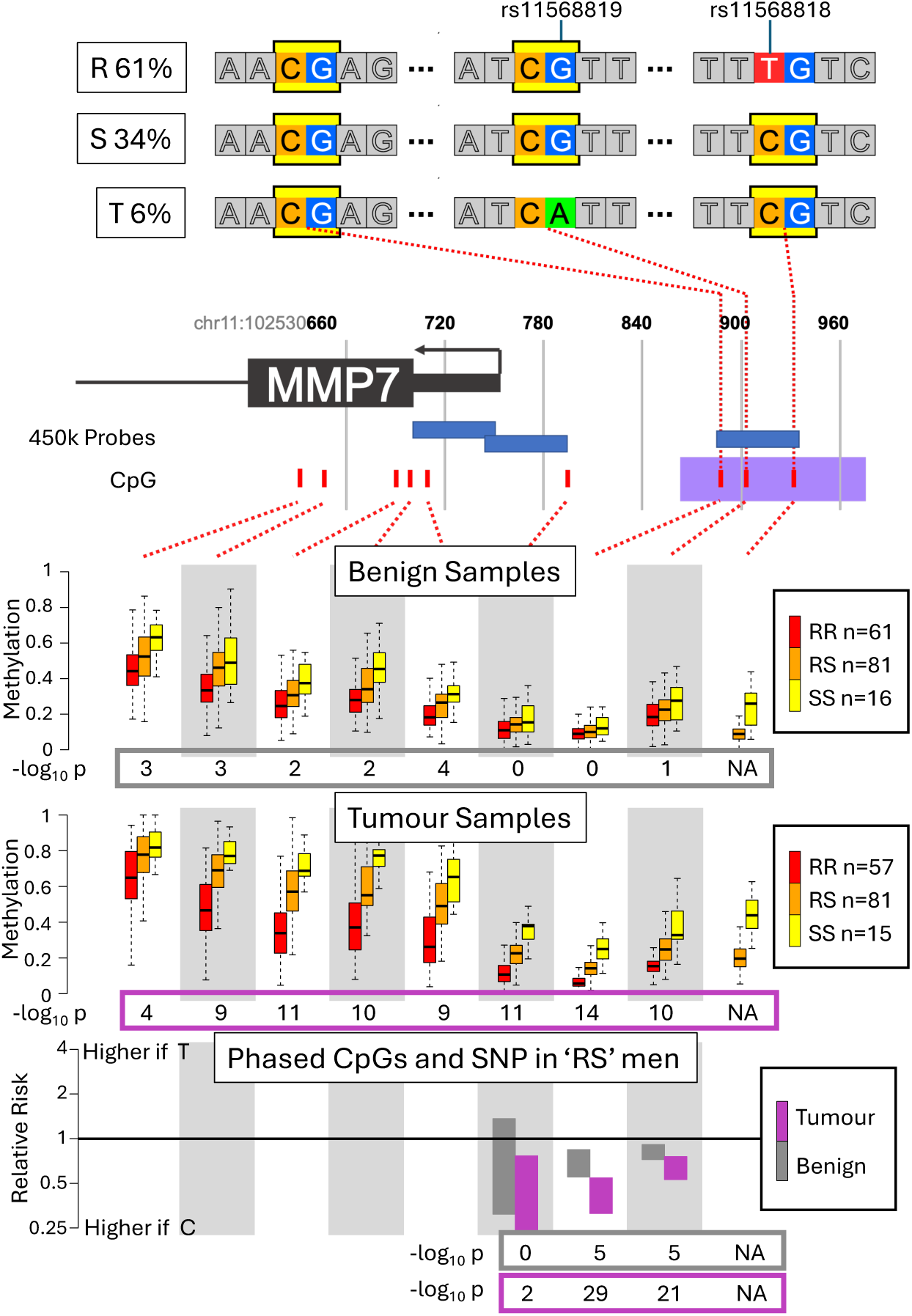
Associations between genomic variants and methylation patterns in the *MMP7* gene. A) Illustrating and naming the three haplotypes, defined by rs11568818 and rs11568819, seen in this cohort. B) Illustrating the promoter and first exon of the *MMP7* gene, with the locations of CpGs, common SNPs, and Illumina 450k array probes indicated. C) For benign samples, for each of the three haplotype combinations possible when rs11568819 is a “G”, illustrating the distribution of methylation values seen at each of the nine CpG sites in the region. D) For tumour samples, for each of the three haplotype combinations possible when rs11568819 is a “G”, illustrating the distribution of methylation values seen at each of the nine CpG sites in the region. E) Within men who are heterozygous at rs11568818, for three CpG sites close enough to phase methylation levels with the SNP, the relative risk of the CpG being methylated if rs11568818 is a “T”. The confidence interval arising from a meta-analysis treating man as a random effect.

A second common SNP in the region (rs11568819) determines the presence of a second CpG, leading to three observed haplotypes in our cohort [Figure 4A]. The promoter and first exon of *MMP7* are shown in Figure 4b. We see a trend with rs11568818 genotype in the methylation at all nine CpGs in the region. The association is clear in the benign tissue [Figure 4c] and more obvious still in the tumour tissue [Figure 4d]. Phasing methylation and genotype in men who are heterozygous at rs11568818, we see that the effects are specific to the molecules carrying the risk allele [Figure 4e]. Methylation is highest across the region when rs11568818 takes the C/G allele.

This association is visible in different subsets of samples (Supplementary Figures S41 to S45). There is negative correlation between *MMP7* expression and methylation, seen in data from TCGA (Supplementary Figure S39) and this cohort. Despite this, the effect of the SNP on methylation appears to be direct, i.e. SNP *→* Methylation *→* Expression, and not SNP *→* Expression *→* Methylation. This is evidenced by the perseverance of the SNP/Methylation association when controlling for expression, and by the effect of silencing methylation on *MMP7* expression in prostate cell lines (Supplementary Figure S40).

We used the GWAS catalogue [33] to obtain details of 2217 prostate cancer risk variants. We can productively investigate 54 through these data (Supplementary Figure S38), and of these 28 show an association between local methylation patterns and the status of the variant. Several of these are driven by trivial SNP/CpG interactions, full details are given in Supplementary Table S10 (More notable examples are shown in Supplementary Figures S46 to S50).

### Hemimethylation levels predict time to recurrence after prostatectomy

EPICseq data on both DNA strands at CpG loci were available for 0.4 million CpGs (1.4% of the total CpG number) enabling investigation of strand-asymmetric CpG methylation, hereafter referred to as hemimethylation. Figure 5A shows sequence coverage for five cancers and matching benign tissues of a run of 10bp across the hemimethylated locus at chromosome 1:1,668,501-1,668,502. Data from the Watson and Crick strands are shown, respectively, above and below the x-axis. The summed reads for all of the cancers (Figure 5B) and benign (Figure 5C) are also shown.

**Figure 5.**
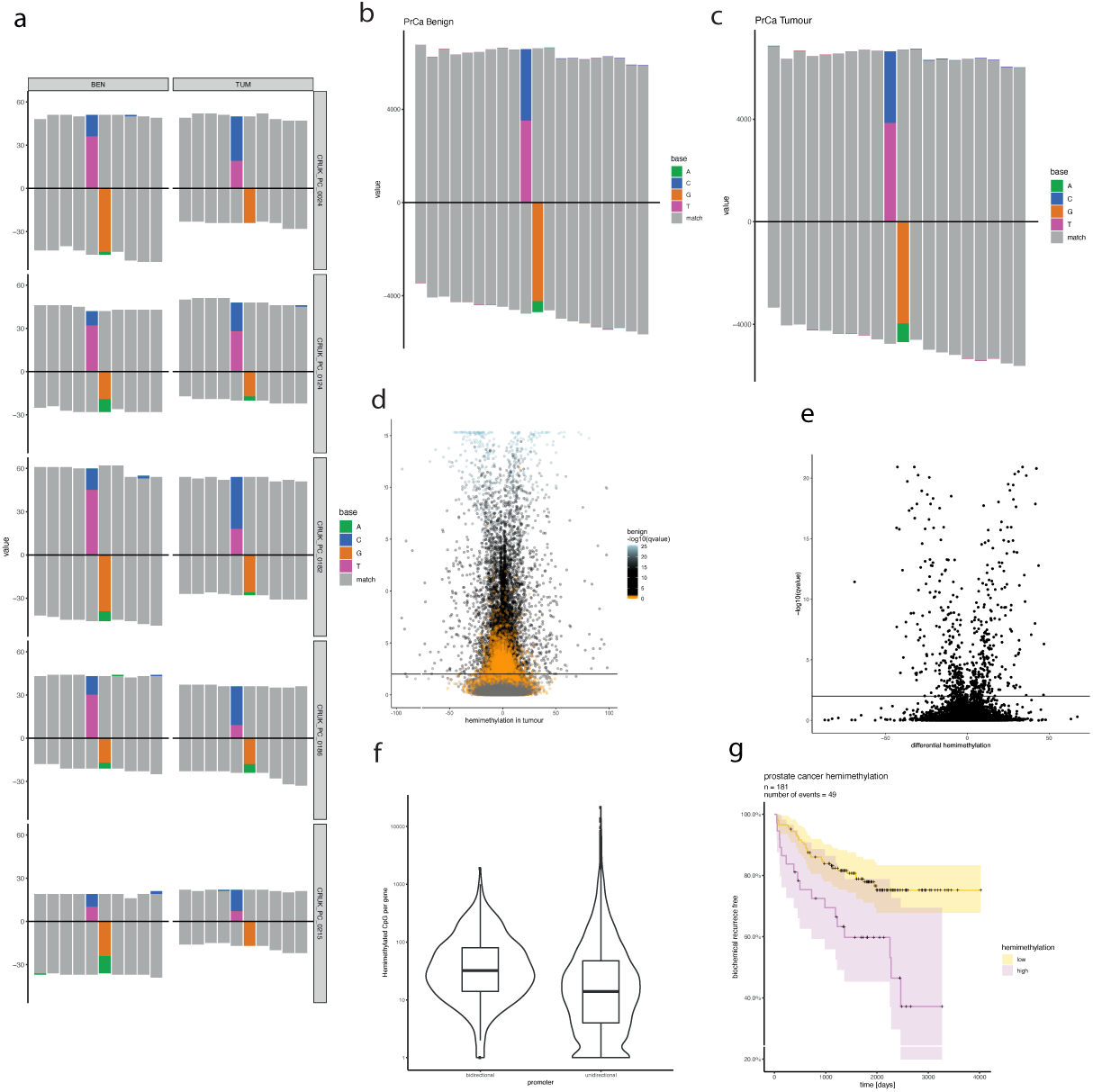
Hemimethylation. (A) For five men, showing a hemimethylated locus at chr1:1,668,501 where Watson strand reads are pictured above the x-axis, bottom strand bases are pictured below the x-axis. Bases matching to the reference genome are displayed in grey. For the hemimethylated CpG, methylated basepairs are cytosine (blue) and guanine (orange) and unmethylated bases are pink (tymine) and green (adenine). (B) A plot aggregted across all benign samples showing the same locus. (C) A plot aggregted across all tumour samples showing the same locus. (D) A volcano plot of hemimethylated sites observed in benign and tumour samples. Hemimethylation in tumour (x-axis) and qvalue (y-axis). The qvalue obtained from benign samples is indicated through colour. (E) Differential hemimethylation between tumour and benign samples. X-axis is difference between tumour hemimethylation and benign hemimethylation. (F) Bidirectional promoters show higher number of hemimethylated CpG when compared unidirectional promoters. (G) Biochemical recurrence in patients showing high hemimethylation (violet) and low hemimethylation (yellow).

For individual CpGs, we define hemimethylation as the distribution of imbalances (across men who could provide an estimate) being centred significantly away from zero (Wilcoxon signed-rank test). We detected 4055 hemimethylated CpGs in tumour tissue, and 3775 hemimethylated CpGs in benign tissue (2893 CpGs common to both tissues) (Figure 5D). Formal testing across our cohort of paired tumour and benign samples detected 1490 significantly differentially hemimethylated sites (Figure 5E).

Functional enrichment analysis of hemimethylated sites in prostate tumours revealed enrichment in transcription factor binding sites including at the *AR*, *FOXA1*, *JUN* and *MYC* binding loci as well as for the *CTCF* transcriptional repressor gene (Supplementary Table S11). Single stranded DNA binding has been observed for mammalian transcription factors and could be mediating this type of modification [34]. Where genes were identified as containing hemimethylated CpGs, the transcribed strand showed lower methylation (–21%, p-value = 0.02566, Wilcoxon Rank sum) consistent with a regulatory role in gene transcription. Notably, genes sharing promoter regions were enriched for hemimethylated CpGs, compared to genes with promoter regions facilitating a gene only on one strand (Figure 5F, p-value *<* 0.0001, Wilcoxon rank sum). For hemimethylated promoters shared by two genes, an imbalance in expression levels increases with the degree of hemimethylation (Supplementary Figure S51).

As well as identifying recurrently hemimethylated CpG sites, it is possible to assign a ‘global’ hemimethylation score to individual men. Men with high hemimethylation levels demonstrated increased risk of recurrence (HR = 2.48, p-value = 0.0025, [Figure 5G, Supplementary Figure S52A) and metastasis (HR = 6.72, p-value = 0.0061, Supplementary Figure S52B).

### Mitochondrial aligning methylation signal predicts time to recurrence after prostatectomy

The consensus that mitochondrial DNA does not undergo methylation has been challenged recently with several papers suggesting a presence [35–37]. While some reports focus on the D-loop and others suggest that methylation is higher in a non-CpG context, there are also concerns about technical biases such as whether the reads are truly mitochondrial, whether there is a technical artefact, whether an alternative epigenetic marker is being detected, and whether changes in DNA morphology are being picked up [38, 39].

In our data we see reads demonstrably originating from the mitochondrial DNA (see Methods), although the methylation signals are low. We quantified those signals occurring at: (a) CpG sequences; (b) CHH sequences, where H is A, C or T; and (c) CHG sequences. Overall, non-CpG levels are correlated with, but lower than those in the nuclear genome. The apparent CHG levels of methylation in the mitochondria are in good agreement with CHH levels, and are correlated with CpG levels (Figure 6A), however the CpG levels are notably higher for a subset of men, (blue dots in Figure 6A). These men with higher CpG apparent methylation signal in the mitochondria see a decreased time to PSA recurrence (Figure 6B) and more rapid metastasis (Figure 6C), indicating a potential link between mitochondrial CpG methylation and clinical outcome.

**Figure 6.**
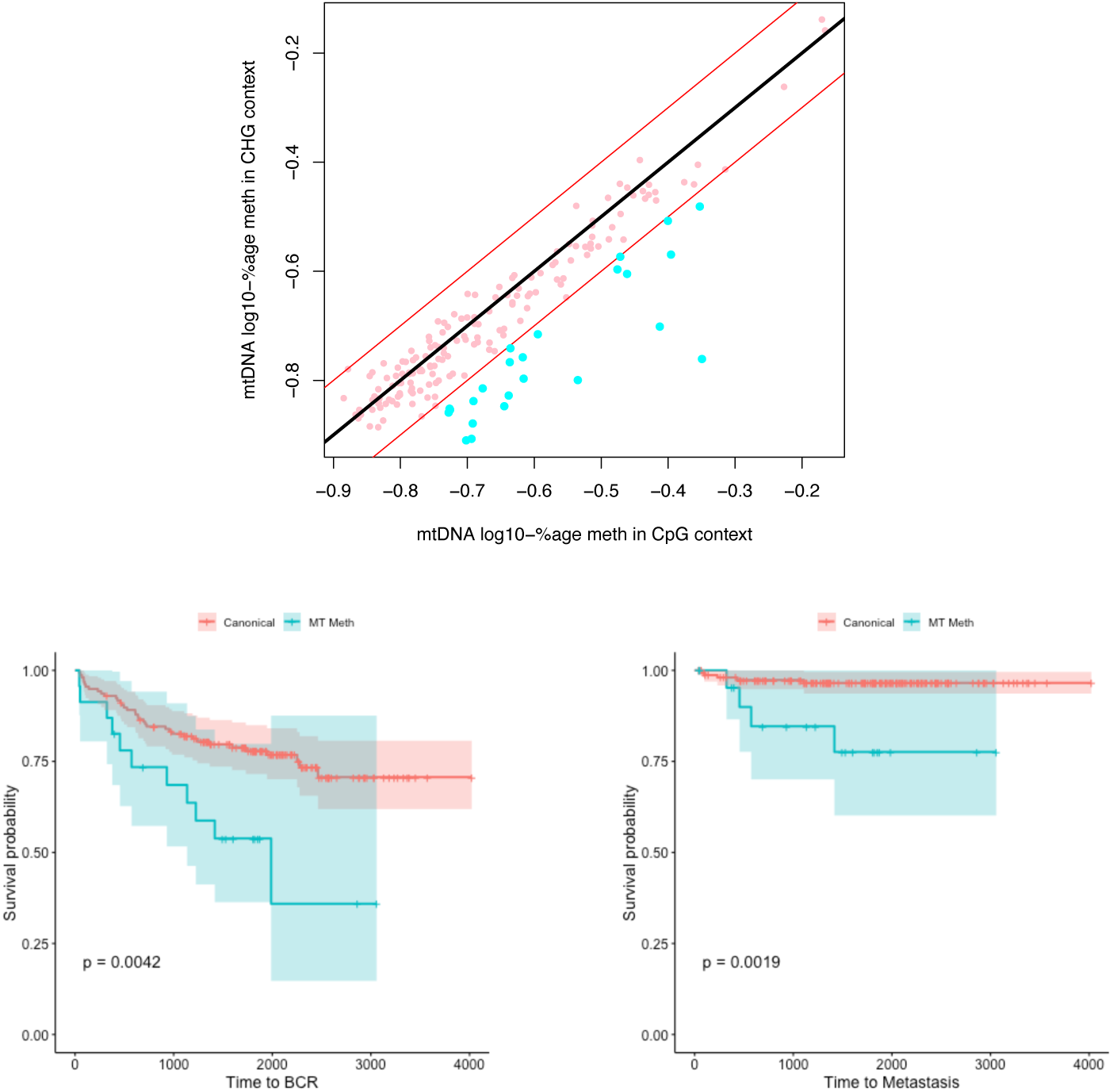
Illustrating weak signal arising from mitochondrial DNA. A) A scatterplot on the log-scale showing signal in CpG contexts within the mitochondrial DNA against signal in CHH contexts. From this we can define a group of men for whom the ratio of signals shows an excess for the CpG context. B) A Kaplan–Meier plot showing the association of the excess signal in mitochondrial CpG contexts and relapse-free survival. C) As for B, but showing time to metastasis.

Examining the methylation signal from mitochondria CpG sites in these higher-hazard men does not identify hot-spots, but rather a fairly uniform spread.

### Combinations of predictive indicators outperform any individual

We have presented above three novel indicators of accelerated time to recurrence (biochemical or metastatic) derived from targeted methylation sequencing data (Supplementary Table S13).

Groups with shorter expected time to recurrence (hereafter referred to as high-hazard) were identified: (a) by clustering of methylation of driver genes (Supplementary Figure S3); with high overall levels of hemimethylation (Figure 5G); with mitochondrial DNA exhibiting higher levels of CpG methylation (Figure 6A). Additionally, we have previously shown that DESNT and Alternative Evotypes also define high-hazard groups.

The driver gene methylation cluster, the high hemimethylation group, and the high mitochondrial-aligning signal group are all defined based on characteristics of the methylation without reference to clinical data. They offer comparable predictive value, but are not identifying the same subsets of men, nor are they picking out the same men as classified by the alternative Evotype or the high-hazard DESNT groups [Figure 7A]. Each of these five stratifications identified a distinct group of men with overlaps of membership of the high-hazard group shown in Figure 7A.

**Figure 7.**
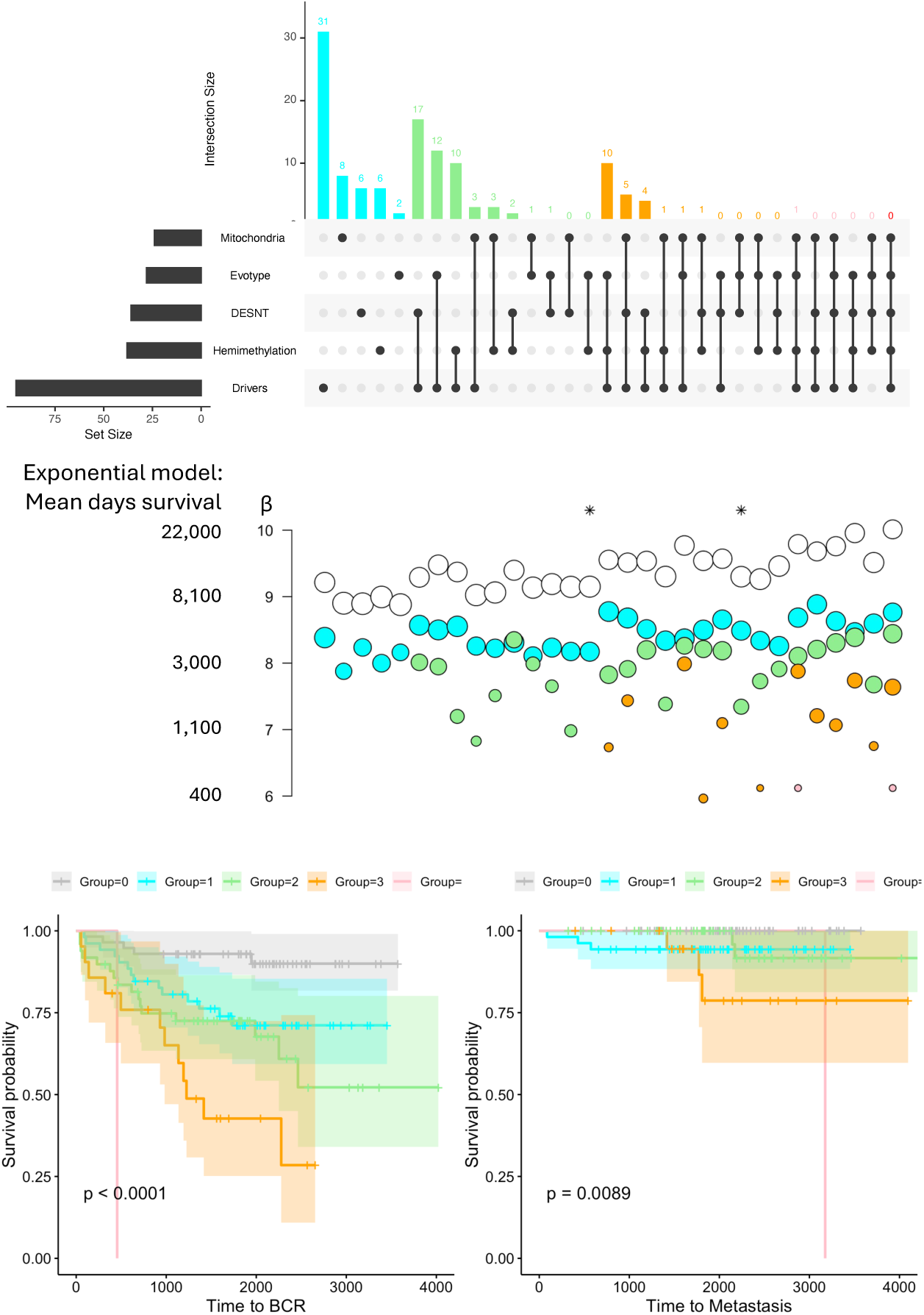
Combinations of risk factors. A) An upset plot indicating the incidence of 31 combinations of high risk groups. Single risk factors are indicated in cyan, combinations of two risk factors are indicated in green, combinations of three risk factors are indicated in peach, and combinations of four risk factors are indicated in pink. The combination of all five risk factors is indicated in red, but is an empty set for this cohort. B) For the sets of variables indicated in the Upset diagram above, a variable is created as the sum of the number of those variables in which a man is to be found. An exponential regression survival model is then fitted to time to biochemical recurrence using this sum as the explanatory grouping factor. Estimated survival for each level of the factor is shown (grey in none of the risk groups, cyan in any one of the risk groups, green for being in any two of the risk groups, peach for being in any three of the risk groups, pink for being in any four of the risk groups. The size of the circle indicates the number of men in that number of risk groups. C) When all five risk groups are considered, Kaplan–Meier estimates of time to biochemical recurrence for men in zero, one, two, three or four of the risk groups. D) As for part C, Kaplan–Meier estimates of time to metastasis.

Since the Alternative Evotype and high hazard DESNT groups are almost mutually exclusive, any man may be within zero to four of these high-hazard groups, and that number of memberships is strongly predictive of time to recurrence [Figure 7C, *p <* 0.0001 log rank test] and time to metastasis (Figure 7D p=0.0089). Moreover, as this figure increases from zero to four, the effect on survival time to biochemical recurrence is monotonic. All five high-hazard groups improve the model. No combination of a subset of the risk groups achieves the same separation of fitted mean survival times [Figure 7B].

## Discussion

The targeted sequencing of DNA methylation in a prostate cancer cohort revealed novel biology and markers pertaining to the risk of prostate cancer, the detection of prostate cancer, and the progression of prostate cancer post-surgery; none of which have been revealed using conventional microarray approaches. We have also provided a unique data resource for the research community. Not only by providing methylation sequencing data for an already well-profiled cohort, but also by providing a substantial data set of methylation profiles for matched benign tissue, including cases of BPH.

Although our estimates of ‘global’ methylation must be considered in light of the functional focus of the CpGs targeted by the EPIC-Seq platform, we have seen increases of methylation from benign tissue to tumour tissue in all contexts, but naturally greatest in CpG contexts. We have seen also that the difference from benign to tumour differs between classes of tumour, being higher in low grade groups, Canonical evotypes, DESNT high-risk groups and in tumours with a TMPRSS2-ERG fusion. These trends, excepting that for grade group, are visible in non-CpG contexts, but to a much lesser degree. These changes in the low levels of methylation in non-CpG contexts are distinguishable from technical noise due to the inclusion of matched tumour and benign samples in the same experimental batches.

While multiple papers have identified genes whose methylation can distinguish prostate tumour tissue, the proposed lists have been inconsistent, and the field has been reliant on a very few genes for the purpose, headed by GSTP1. Our results suggest that one reason for the variation in the genes reported previously is that each underpowered study has returned a random subset from a very large pool of genes that can perform the function. Another contributing factor is the degree to which many previous studies have had to summarize the methylation profiles of a gene or gene promoter, either for practicability or due to the technology being used. We have seen that the methylation profiles of some genes are quite complex, and can not only distinguish tumour and benign tissue, but can reveal characteristics of the tumour, in a manner that may be lost with an automated summarization.

DNA methylation changes around *GSTP1* have long been studied as a possible biomarker for prostate cancer with potential for non-invasive detection [8, 40]. While *GSTP1* methylation performs admirably as a biomarker [40], by identifying more differentially methylated genes, and confirming the results of earlier studies that did likewise, we increase the scope to develop powerful non-invasive clinical tools for prostate cancer, while providing greater flexibility and power for studies requiring automated assessment of pathology.

We have seen also that SNPs associated with an increased risk of prostate cancer can directly influence local methylation patterns. In the case of rs11568818 near MMP7, we have shown evidence that the effect is direct, and not mediated by expression of the gene. rs11568818 is an example of a SNP that creates or removes a CpG sequence. We have shown that this is not a necessary characteristic for a SNP to affect local methylation levels, but it provides an obviously direct mechanism for so doing, and more than a quarter of the SNPs associated with prostate cancer risk either create or remove a CpG. The majority of such SNPs lie without the regions targeted by the EPIC-Seq panel, and so are not examinable within these methylation data. These findings invite the reappraisal of approaches for the prioritization of germline mutations for investigation after genome-wise association studies, but also, presumably, the characterization and prioritization of somatic mutations at or near CpG locations.

Understanding the effect of SNPs on methylation patterns also invites the reinterpretation of large prostate methylation microarray studies. Being able to explain variation in methylation data not associated with the effect of interest can only increase the power of a study, while appreciating that polymorphisms do not have to be within a probe sequence in order to affect the methylation seen by that probe could cast new light on microarray probe quality control processes.

Aside from tools for classification, and elucidation of risk mechanisms, the main contributions of this work lie in the identification of characteristics that stratify men into groups with different hazard distributions. The cohort are post-surgery cases with an over-representation of intermediate risk cancers and represent men for whom additional tools for and insights into risk stratification could be most valuable. The three predictive characteristics we identify from the EPICseq data are the stronger for having been derived to capture an aspect of the cancer biology that would not obviously be associated with survival, and nor did they require knowledge of survival for their discovery and definition. That the classifiers are identifying different subsets of men is also a strength.

The strand-specific ‘hemimethylation’ patterns we have shown may be essential if we wish to allow for epigenetic control of the expression of genes that share promoter regions, and we have seen enrichment in these areas. The characterisation of hemimethylation is not complete with this work, and the potential for re-evaluating previous microarray studies is substantial in light of these results.

The reporting of methylation in mitochondrial DNA is controversial despite a number of recent studies suggesting it may occur [35–37]. We are not reporting the high levels or hot-spots of mitochondrial signal that would require a mechanism for copying patterns from one molecule to descendent molecules to explain them, but rather a low-level, apparently random, signal that would be consistent with infrequent independent *de novo* events.

Nevertheless, with these data we cannot show that the cause is DNA methylation, and not some other physical or epigenetic aspect that would inhibit bisulfite conversion. What we can be sure of, is that there is a signal arising from the mitochondria following bisulfite conversion that for a group of men is higher in CpG contexts than other contexts (and never the other way around as might be expected if this were a technical artefact), and that the men with this characteristic have poorer times to recurrence. Given the potential utility of the signal, it demands further investigation.

Our key finding is that the three novel classifications presented here, each developed in an unsupervised manner, can be combined with existing high-hazard groups defined by DESNT [20] and Evotype [7] to create a classifier with much greater power and discrimination. This, despite cohort characteristics which should hamper such performance. At present we note that to generate all five classifications would require, at least, targetted methylation sequencing, mRNA expression, and further genomic sequencing. We have shown though that it may be possible to call all five classes from targetted methylation sequencing. Coupled with the rise of assays that can report on DNA sequence and methylation patterns simultaneously, the potential for a single assay classifier for our five classes is strong.

### Limitations of the study

The genomic phase of the UK prostate ICGC cohort had a deliberate bias towards intermediate-grade-group tumours (conditional on these being prostatectomy cases) as a cohort for which new tools might provide the greatest benefit, and we are constrained by that cohort definition. A consequence is that we may lack sufficient variance for established clinical tools to demonstrate their true performance. It is not surprising then that adding while the Grade Group is a known predictor of time to biochemical relapse, adding it into the final model for our restricted cohort has only a moderate effect (Figure S54). This neither diminishes existing predictors, nor invalidates the results we present, but rather constrains the patient population for which we can claim evidence that the results we present will provide benefits.

A limitation of the study is that the different prognostic tools have been developed in the same cohort, and the unique characteristics of the data set that are being exploited mean that data for external validation cannot be identified. However, while the prognostic tools have been developed here in the same cohort, as has been seen, different high-risk groups are identified by each tool.

## Methods

### The cohort, clinical data and sample selection

DNA methylation data at base-pair resolution were generated from 190 patients within the CRUK ICGC cohort [Supplementary Table S1]. Supplementary Figure S1] for which tissue was available. Matched pairs of tumour and pathologist-assessed-benign tissue samples (elsewhere in the paper referred to as ‘benign’) from the same prostate were collected.

Extracted DNA samples were bisulfite-converted, amplified and sequenced in pools of 12 samples (150 single end reads over two Illumina HiSeq4000 lanes; Supplementary Table S2).

### Library preparation and sequencing

Genomic DNA samples from fresh-frozen prostate tissue samples were extracted using the Qiagen AllPrep kit and fragmented using the Covaris system. Targeted methylation library preparations were generated using the TruSeq Methyl Capture EPIC platform (Illumina). Briefly, fragmented DNA was used to generate sample barcoded PCR-free library preparations, followed by equimolar 4-plex pooling and hybridisation with sequence-capture baits covering 107Mbp of the human genome (EPIC targets). The resulting target-enriched DNA library pools were subjected to sodium bisulfite conversion, followed by library amplification using universal adapter primers. Bisulfite converted, target-enriched library pools were combined in sets of 3 (i.e. 3 x 4-plex pools) and sequencing was performed on the Illumina HiSeq4000 platform (two lanes of SE150 per 2-plex pool).

Benign tissue was not available for two men, meaning that 378 EPIC-seq libraries were generated in batches of 12. Benign and tumour samples from the same man were processed in the same batch.

### EPICseq data processing

Bisulfite sequencing data were pre-processed, aligned to the human genome (GRCh38) and analysed using a combined workflow with Bismark [41] and Methylkit [42].

### Copy number profiles

The identities of 785,583 A*>*T SNPs were downloaded from the UCSC Table browser (https://genome.ucsc.edu/cgi-bin/hgTables) and filtered to 716,986 on autosomal chromosomes. A reported minor allele frequency of at least 0.1 was required, and a high-coverage sample was investigated to determine which SNPs had at least 20 reads of coverage in that sample. Filtering on these two properties reduced the list to approximately 12 thousand SNPs that were investigated in each sample. A and T allele counts for each SNP in each sample, as well as the total coverage for a 2001bp window centred on the SNP, were recorded. Only SNPs that had a median coverage in their 2001bp window of more than 40 reads in benign samples were retained, leaving 11,628 [Supplementary Table S12]. For some analyses, 173 SNPs in the Major Histocompatibility Complex region were additionally removed.

An average of approximately 3,300 SNPs were heterozygous per sample. Diagnostic plots were generated for each sample, normalized against a panel of benign tissue samples. Autocorrelations along the genome were calculated, and segmentation performed using the DNAcopy [43] package from BioConductor.

### EPICseq quality control

Sample identities were confirmed at the man level using germline variants identified in previously published whole-genome sequencing data [7], and a minimum of 15 million reads were required. Tissue type (Tumour or Benign) was verified via DNA copy number profiles for the autosomal chromosomes, *GSTP1* methylation profiles, and the presence of somatic mutations identified in whole-genome sequencing data.

Nominally Benign samples that had high *GSTP1* methylation or showed evidence of copy number changes were investigated, as were nominally Tumour samples that had low *GSTP1* methylation and no evidence of copy number changes. Comparison of the matched samples from that man, expected copy number changes and variants from matched whole-genome sequencing data, and pathologist’s assessment were used to determine final status.

17 samples were removed from the potential 378 cases following these processes. Even after this, there are some samples that do not behave as expected when looking across the 100, 1, 000 or 10, 000 most discriminating genes [Supplementary Figure 5].

The benign sample from CRUK PC 0045 is classified as tumour by the vast majority of genes (approximately 90%), including known classifiers. Yet in comparison with the tumour sample [Supplementary Figures S55 and S56], and investigation of somatic variants, the classifications are confirmed: The locations of 109 somatic variants identified in the WGS data had sufficient coverage for investigation in the tumour methylation data and 93 (85%) had some support, while for the benign methylation sample this was only 4/113 (3%).

Three other samples (Tumour samples from CRUK PC 0160, CRUK PC 0222 and the Benign sample from CRUK PC 0202) are misclassified by more genes than we would expect, but not so many that we have strong evidence of a switch of sample type. [Supplementary Figures S57 to S62] depict these, and demonstrate that the tumour samples, in particular, are correctly labelled. Additionally, the tumour sample from CRUK PC 0160 has some support for 100/143 somatic variants, the tumour sample from CRUK PC 0222 has some support for 75 out of 100 somatic variants, and the benign sample for CRUK PC 0202 has support for no variants (out of 93 with sufficient coverage) in contrast to the matched tumour sample which has support for 100/117 somatic variants. With evidence to support the tissue classifications, these samples are retained in analyses, but place a limit on the performance of classifiers etc.

### Methylation analysis methods

#### DMR/DMC methods

Differentially methylated cytosines (DMCs) were detected using Methylkit [42]. All benign and tumour samples were analysed in unpaired comparison with patient identifiers as covariates. Differentially methylated regions (DMRs) were created by merging DMCs within 50bp radius of each other. Regions with at least two DMCs per region were selected as DMRs. For each DMR the DMC with the highest absolute methylation difference was selected as DMR methylation difference.

#### Driver gene and clustering methods

A list of prostate cancer driver genes was defined by somatic mutation enrichment above background models from two papers [5, 6], and promoter region methylation levels could be identified from these data for 158 such genes. Promoter regions of the genes were annotated using GENCODE [44], and average across all differentially methylate cytosines were calculated for each promoter. Imposing a requirement that the majority of samples provide a value for each gene used, we reduce to a panel of 150 genes.

Clustering of the Tumour sample values for the 33 most variable genes was performed using the “ward.D2” method in the hclust package in R.

#### Tumour/Benign association methods

The performance of genes in distinguishing tumour and benign tissue was assessed as following. For 344 libraries representing 172 tumour and 172 matched benign samples, we identify the methylation levels for all CpGs within 1000 bases of the gene coordinates. Beta values from all CpGs with a median absolute deviation (MAD) across the 344 libraries of at least 0.14826 (the theoretical MAD associated with a uniform 20% shift in methylation from tumour to benign in the absence of other variation) were used for the unsupervised hierarchical clustering of the 344 samples. The dendrogram was cut to provide two clusters, and the two clusters investigated for association with the tissue type.

A similar process was used to investigate the association of genes with TMPRSS2-ERG fusion status, DESNT risk group status and Evotype status.

Where we consider a panel of 10 driver genes, the classifications from each gene in the panel are tabulated and the ‘winner’ recorded or the proportion of genes correctly voting for the winner is recorded. Although this approach allows for some information leakage, simulations show that in the absence of signal, this raises the mean (over samples) classification score from 0.5 to 0.52, while the actual value seen in Figure 2C is 0.97, so we do not consider this a concern.

For panels of 100 genes we report on the ability to classify each sample based on knowing the tissue status of the other 343 in the set, but otherwise follow the same procedure.

### Hemimethylation methods

All CpGs with coverage greater or equal to 10x in at least 10 samples were analysed. Top strand methylation was compared to bottom strand methylation using the Wilcoxon signed-rank test. Sites showing tumour-specific hemimethylation were annotated using ChEA [45]. For strand specific methylation, promoter annotation was obtained from Ensembl (release 100) and hemimethylation of sites overlapping the promoter compared with the strand of protein coding sequence [46]. CpGs that overlap common SNP that mimic methylation changes were removed (dbSNP v155, [47]).

Hemimethylation (HM) for an individual sample was defined as a difference between Watson and Crick strand methylation for the given CpG (Equation HM).

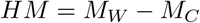

Where *HM* is the hemimethylation score, *M_W_* is the methylation level of the Watson strand, and *M_C_* is the methylation of Crick strand.

For CpGs with strand-balance information represented in at least ten samples, Watson strand methylation was compared to matched Crick strand methylation using the Wilcoxon signed-rank test. Benign and tumour samples were analysed separately. CpGs were denoted as Benign-specific hemimethylation sites and tumour-specific hemimethylation sites if they showed significant imbalance in only one tissue type. The remaining hemimethylation sites were defined as shared.

The difference for a man between paired benign and tumour hemimethylation levels, where both were calculable, was defined as:

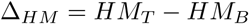

where *HM_T_*and *HM_B_*are the hemimethylation scores calculated for the matched tumour and benign tissues.

For CpGs where tumour and benign methylation scores were both available for at least ten men, the changes between benign and paired tumour tissue hemimethylation were compared with the Wilcoxon signed-rank test.

#### Functional enrichment

Sites showing tumour-specific hemimethylation were annotated for transcription factor binding potential using ChEA [45]. For strand-specific methylation, and hemimethylation of sites overlapping the promoters, promotor annotations were obtained from GENCODE (release 100, [44]).

#### Strand specific promoter analysis

For strand-specific promoter analysis we identified the transcribed strand with GENCODE ([44]).

A list of bidirectional gene promoters were obtained from the Comprehensive Human Expressed SequenceS [48]. Enrichment of bidirectional gene promoters was assessed by comparison with all genes for which hemimethylation was assessed.

#### EPICseq-wide hemimethylation analysis

EPICseq-wide hemimethylation was estimated by averaging absolute hemimethylation for all observed CpGs. Patients were split into high-hemimethylation (80th percentile) and low-hemimethylation groups. Progression free-survival and metastatic events in these groups were compared using Cox proportional hazards model.

### GWAS-identified risk allele associations

The GWAS catalogue was accessed on the 23rd July 2024 to identify 2855 prostate-associated variants. These were filtered to include only variants on chromosomes 1 to 22 and chromosome X, variants associated with the ‘Prostate Cancer’ trait, and which have been identified in a European cohort. Duplicate locations were removed, and only variants for which we have coverage of at least 10 reads in one strand for at least 50 benign sequencing libraries.

To avoid alignment biases, we restricted to cases where the risk allele is a single base. These were annotated to check whether a CpG is created or removed, and if a particular strand was required to see the variant outwith the effects of bisulfite conversion (G*>*A variants need the positive strand, C*>*T need the negative strand).

For basic results, values were calculated for each man using the Rsamtools pileup function. Phased analyses were conducted as random-effects meta-analysis treating men as individual experiments. The ‘metabin’ function in the ‘meta’ R package was used.

### Mitochondrial methods

The above-described processing of the data results in reads being aligned to the mitochondrial genome. Despite having no reason to expect these regions to be targeted, we believe these to be genuinely originating from the mitochondrial genome for the following reasons:

- The mapping qualities do not indicate any problem.
- The sequencing depth of the mitochondrial DNA is high relative to the nuclear genome. To the extent that it would seem to be unlikely that it is coming from NUMT regions.
- The sequencing depth of the mitochondrial DNA differs between tissue types (e.g. being much lower in a blood sample for man 0001 (not shown)).
- Mitochondrial mutations unambigously identified in WGS data where they can be supported by tens of thousands of reads are visible in the methylation sequencing. For some examples see Table 2.

**Table 2.** Examples of high allele fraction mitochondrial mutations identified from whole-genome sequencing data in the cohort, and the observed allele fraction in the methylation sequencing data, demonstrating that the reads are correctly mapping to the mitochondrial genome. Coordinates are given relative to the revised Cambridge Reference Sequence.

| Man | Locus | Mutation | WGS frequency | Methylation freq. |
| --- | --- | --- | --- | --- |
| Man 0025 | ChrM:152 | T>C | 16,123 / 21,001 (77%) | 76 / 92 (83%) |
| Man 0025 | ChrM:2604 | T>C | 8,620 / 16,359 (53%) | 20 / 30 (67%) |
| Man 0072 | ChrM:1177 | G>A | 1,072 / 4,930 (22%) | 299 / 912 (33%) |
| Man 0027 | ChrM:8293 | G>A | 6,008 / 16,960 (35%) | 13 out of 62 (21%) |

Table 2 also highlights the inconsistent coverage of the mitochondrial DNA, and the potential ability of the platform to call mutations in the mitochondrial genome where coverage allows. Note that the majority of the mutations seen in the mitochondria are ones that will be confounded with bisulfite conversion if the wrong strand is used.

For the men with no excess CpG methylation signal (relative to non-CpG contexts), there is structure to the apparent noise, but for the men with signal in the mitochondrial-mapping reads, the signal is consistent with there being a low-level uniform and independent chance of signal at each CpG.

### Survival analyses

Unless otherwise specified, survival analyses were non-parametric, using the Kaplan–Meier estimator and the log-rank test.

Since Figure 7C is suggestive that an exponential survival curve would represent the data well, an accelerated failure time survival regression was fitted to the data to generate the median survival estimates for each group in Figure 7B.

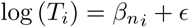

where *T_i_*is the time to relapse for man *i*, *n_i_* is the number of risk groups in which man *i* is found, *β_m_*is the value assigned to the men in *m* high-risk groups, and *ɛ* is an error term taking a Gumbel distribution.

Noting that when all five risk categories are used, the survival estimates are monotonic and roughly evenly spaced on a log scale, we also fit a two parameter model

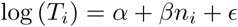

where a *β* value less than zero indicates that the effects of each classification are cumulative in the manner expected. The robustness of the model is assessed via bootstrapping for 5000 iterations using the R boot library [49].

### RNA-seq and TMPRSS2-ERG fusions

Matched RNA libraries were identified from the equivalent RNA project of the CRUK ICGC Prostate Cancer Group described elsewhere [Manuscript in preparation]. Gene expression was evaluated using STAR and TMPRSS2-ERG fusions were identified using STAR-FUSION [50].

### Data Accession

Sequencing files are stored at the European Genome-Phenome Archive, accession number: EGAC00001003628.

## CRUK ICGC Prostate Cancer Group Membership

## Acknowledgments

We thank the patients and families that have made this research possible.

The experimental work and sequencing were funded by Cancer Research UK via the Prostate International Cancer Genome Consortium (ICGC) grant ICGC/A21373, which also supported W-KL. Collaboration across the analysis network was supported by the Isaac Newton Institute and EPSRC grant (Ref: EP/V521929/1).

RPL, SP and CEM were supported by a CRUK Career Development Fellowship (C50604/A26718).

VH, TB-S, DSB and CSC are grateful for and acknowledge support from The Masonic Charitable Foundation Successor to The Grand Charity, Movember, The Prostate Cancer Research, The Big C Cancer Charity, The King Family and the Stephen Hargrave Trust. They acknowledge support from Cancer Research UK C5047/A14835/A22530/ A17528, C309/A11566, C368/A6743, A368/A7990, C14303/A17197, the Dallaglio Foundation (CRUK Prostate Cancer ICGC Project and Pan Prostate Cancer Group). Their work was funded by Prostate Cancer UK, research grant ref: MA-ETNA19-003, RIA15-ST2-029 and TLD-CAF22-011)

Vincent J Gnanapragasam acknowledges infrastructure support from the National Institute for Health and Care Research (NIHR) Cambridge Biomedical Research Centre(NIHR203312). The views expressed are those of the authors and not necessarily those of the NIHR or the Department of Health and Social Care.

DCW is supported by the NIHR Manchester Biomedical Research Centre (NIHR203308).

We thank Simon Tavaré, James Hadfield, Clarissa Gerhauser, Greg Leeman and Felix Kreuger for helpful input and discussions. We thank Tom Drever for help in making the data accessible.

## Author contributions

AB, SM, SS, SH, HL, JK. HCW, JR, DJW, VH, KO’N, T M-C, MC, HD, AL, FCH, CV, CF, DEN, VJG, AYW, RAE, DSB, ZK-J, CEM and AGL contributed to the identification of samples, preparation of nucleic acids, data generation and clinical annotation. WKL and SF performed and oversaw the sequencing experiments. RPL, SP, CEM and AGL performed the analyses with inputs from MK, DJW, DW, DSB and CSC.

CEM cowrote an early version of this manuscript and CSC, DCW, RAE, and DSB made substantial contributions to the organisation and conduct of the study and critiqued the output for important intellectual content

The CRUK ICGC Prostate Group created and maintains overall project direction; the principal investigators being GSB, CEM, AGL, DSB, RAE, CSC, and DCW.

## 1 Supplements

### 1.1 Supplementary Tables

**Table S1**

**Table S2.** Sequencing metrics for the project. The table consists of, for each man in the study, The Study ID assigned to the individual; The BAM file associated with the Tumour sample for this individual; The batch in which the sample was run; Bismark metrics for the Tumour BAM file, The BAM file associated with the Benign sample for this individual; The batch in which the sample was run; and Bismark metrics for the Benign BAM file.

**Table S3.** Methylation levels (as percentages) in Tumour samples associated with 150 Prostate Cancer driver genes. Columns are headed by the UK ICGC ID as defined in Supplementary Table S1. Rows are annotated with gene symbols.

**Table S4.** Methylation levels (as percentages) in Benign samples associated with 150 Prostate Cancer driver genes. Columns are headed by the UK ICGC ID as defined in Supplementary Table S1. Rows are annotated with gene symbols.

**Table S5.** Differentially methylated cytosines (DMCs) identified in a comparison of tumour and benign tissue. Columns indicate location of cytosine in CpG context and results of logistic regression model comparing tumour and benign tissue. Average difference between the tumor and benign tissue is also indicated.

**Table S6.** Differentially methylated regions (DMRs) identified in a comparison of tumour and benign tissue. Columns indicate start and end of the DMRs, number of cytosines showing significant methylation difference within the DMR, as well as the results of logistic regression model comparing tumour and benign tissue for the most extreme cytosine in the region.

**Table S7.** Gene-wise associations within 172 tumour-benign pairs. For 20049 genes, we report the gene symbol and coordinates used (chromosome, start and end in hg38). Then for seven regions [5000-1001 based upstream, 1000-501 bases upstream, 1-500 bases upstream, the gene region, 1-500 bases downstream, 501-1000 bases downstream, 1001-5000 bases downstream] we report (E-K) The number of CpGs examined, (L) the minimum p-value associated with any CpG [tumour vs benign, paired t-test] within 1000 bases of the gene, (M-S) The number of CpGs showing significant differences between tumour and benign [paired t-test, holm adjusted p-value¡0.05], (T) the minimum p-value associated with any CpG showing a difference of tumour – benign *>* 0.2, (U-AA) The number of CpGs showing significant differences between tumour and benign and a mean difference of tumour – benign *>* 0.2, (AB) the minimum p-value associated with any CpG showing a difference of benign – tumour *>* 0.2, (AC-AI) The number of CpGs showing significant differences between tumour and benign and a mean difference of benign – tumour *>* 0.2, (AJ) For an unsupervised clustering on methylation (of all CpGs with a median absolute deviation across the 344 paired samples of at least 0.14826), if we define two clusters, the size of the second largest cluster, (AK-AN) For the same clustering, as for AJ if we define 3,4,5 or 6 clusters, (AO) The number of CpGs used in the clustering, (AP) The p-value for the association between the clusters defined in AJ and the tumour/benign status of samples [Fisher’s exact test] (AQ-AT) The p-value for the association between the clusters and the tumour/benign status of samples when 3-6 clusters are used [Fisher’s exact test, p-value estimated from 10,000 simulations], (AU-AY) For different numbers of clusters, the number of samples miss-assigned if each cluster is set uniformly to be the most prevalent sample type in that cluster, (AZ) The number of studies in which [10] saw the gene had been identified as differing between tumour and benign tissue, (BA) Whether the gene is in our driver gene list. Note that the results for GSTP1 are not valid, as this gene was used for quality control purposes, but they are included here for completeness.

**Table S8.** Contingency tables for Evotype, TMPRSS2-ERG Fusion and DESNT status for 136 cases with all three data known.

**Table S9.** Gene-wise associations within 182 tumours. For 20049 genes, we report the gene symbol and coordinates used (chromosome, start and end in hg38). Then for seven regions [5000-1001 based upstream, 1000-501 bases upstream, 1-500 bases upstream, the gene region, 1-500 bases downstream, 501-1000 bases downstream, 1001-5000 bases downstream] we report (E-K) The number of CpGs examined, (L) the log of the minimum p-value associated with any CpG [five groups (alternative evotype + DESNT *×* ERG fusion), Kruskal-Wallis] within 1000 bases of the gene, (M-S) The number of CpGs showing significant associations with the five groups for each region [Kruskal Wallis, number with holm adjusted p-value¡0.05], (T) the log of the minimum p-value associated with any CpG [Association with ERG fusion, t-test] within 1000 bases of the gene, (U-AA) The number of CpGs showing significant associations with fusion status for each region [t-test, number with holm adjusted p-value¡0.05], (AB) the log of the minimum p-value associated with any CpG [High risk DESNT group vs the rest, t-test] within 1000 bases of the gene, (AC-AI) The number of CpGs showing significant associations with the DESNT status for each region [t-test, number with holm adjusted p-value¡0.05], (AJ) the log of the minimum p-value associated with any CpG [Alternative vs Canonical evotype, t-test] within 1000 bases of the gene, (AK-AQ) The number of CpGs showing significant associations with the Evotype status for each region [t-test, number with holm adjusted p-value¡0.05].

**Table S10.** Summary of results for the 74 GWAS-identified SNPS associated with cancer risk that we investigate. (A-C) The date, first author, and journal of the GWAS that identified the risk variant, (D-E) The location of the SNP, (F) The genes that the GWAS catalogue reports as being associated, (G-H) The ID of the SNP and the risk allele, (I-J) The reference base for the SNP and the sequence around the SNP with the reference base, (K-L) The alternative base for the SNP and the sequence around the SNP with the alternative base, (M) The minor allele percentage observed in our benign cohort [Too low a percentage and we have no ability to detect an association], (N) The number of CpGs examined in the region of the SNP, (O) The distance from the SNP to the nearest CpG [Too far and we cannot directly phase the allele with methylation status], (P) mean methylation in benign tissue at the nearest CpG, (Q) Coverage at the SNP in benign tissue, (R) mean methylation in tumour tissue at the nearest CpG, (S) Coverage at the SNP in tumour tissue, (T) Whether the SNP changes the number of CpG sites in the region, moves a CpG, or doesn’t affect the CpG numbers, (U) Whether there is an apparent association with methylation patterns in benign tissue, (V) Whether there is an apparent association with methylation patterns in tumour tissue, (W) Comments on any associations.

**Table S11.** ChIP Enrichment Analysis (ChEA). Columns represent results of gene set enchriment analysis as defined by ChiP-X Enrichment Analysis.

**Table S12.** The 11628 A/T SNPs used for copy number profiles, giving the positions and IDs of the SNPs as well as the reported minor allele frequency.

**Table S13.** For each man, indicating the DESNT and Evotype categories into which the man falls, as well as the three novel methylation categories defined in this manuscript.

### 1.2 Supplementary Figures

**Figure S1.** Flowchart of Cohort Samples. A flowchart depicting the numbers of samples available for analyses.

**Figure S2.** A) Showing the difference, between tumour and matched benign tissue, of EPICseq-wide methylation levels of DNA cytosines in CpG contexts. Data are shown for 173 matched pairs. B) Showing the difference, between between tumour and matched benign tissue, of EPICseq-wide methylation levels of DNA cytosines in CHG contexts (i.e. CAG, CCG or CTG). Data are shown for 173 matched pairs. C) Showing the difference, between tumour and matched benign tissue, in EPICseq-wide methylation levels of DNA cytosines at CHH contexts (i.e. where neither of the two bases following the cytosine is a guanine). Data are shown for 173 matched pairs.

**Figure S3.** The difference in survival between the left-hand and right-hand clusters identified in Figure 1 is illustrated (A Kaplan–Meier curve with log-rank test). The associations between the two clusters identified in Figure 1 and various characteristics of interest.

**Figure S4.** Illustrating the Alternative-Evotype-rich and poor-prognosis DESNT-rich sub-clusters. A) Illustrating the heatmap of methylation levels from 182 primary tumour samples with three clusters indicated. Mean values for the three clusters are shown to the right of the heatmap, while the proportions of Alternative Evotype tumours, DESNT tumours, and tumours with TMPRSS2-ERG fusions in each cluster are indicated below the heatmap. The higher-risk cluster seen in Figure 1 now splits into two, one of which is enriched for DESNT tumours and we refer to as DESNT rich, and one which is enriched for Evotype-1 tumours which we refer to as Evotype 1 rich. B) For the 33 driver genes used for clustering, a scatterplot of the mean levels of expression between the DESNT rich cluster and Evotype 1 rich cluster. The four most discrepant genes are indicated. C-F) Boxplots of the methylation levels of the four genes indicated in B) for DESNT tumours, Evotype 1 tumours, and tumours with the TMPRSS2-ERG fusion.

**Figure S5.** Differentially methylated regions in prostate cancer. (a) Hierarchical clustering of differentially methylated regions in prostate cancer. (b) Number of patients (y) in which the proportion of DMRs (x) are seen to be differentially methylated. (c). Number of DMRs (y) in which the proportion of patients with that DMR is as indicated (x) (d). K-means clustering demonstrates epigenetic markers defining increased risk of recurrence

**Figure S6.** Differentially methylated genes that have been seen previously. A) Genes are ranked by the performance of their methylation patterns in distinguishing tumour and benign tissue, and for every group of 100, the proportion of genes that had been reported previously was calculated. B) Genes are classified by the average methylation change in their promoter region, and for each class the proportion of genes that had been reported previously was calculated. C) The influence of the number of CpGs in the gene promoter and gene body, and the number of individually significant CpGs in the gene promoter and gene body, are plotted against the probability of the gene having been reported previously.

**Figure S7.** Illustrating, for selected CpGs in *AJAP1*, the observed methylation levels in each of the five groups of tumours. Bottom we see the gene, coordinates, and locations of CpGs. Then, moving up, we see violin plots for selected CpGs for the alternative Evotype samples, for DESNT high-risk samples with a detected *TMPRSS2-ERG* fusion, for other samples with a detected *TMPRSS2-ERG* fusion, for other high-risk DESNT samples, and for tumour samples with none of the three characteristics (high-risk DESNT, alternative Evotype, or fusion). Profiles for Benign samples are shown at the top for comparison, and then we see mean values for the five tumour categories and Benign tissue in the top section of the plot. For the plots of the mean profiles, the tumour samples with none of the three characteristics are shown with a narrow black like, while the benign samples have a dashed line. For the violin plots, and mean methylation plot, the CpGs have been equally spaced. Below the first of the violin plots lines indicate the loci in the gene at which the CpGs are situated.

For much of the region, all tumour categories track the benign profile, except for the Alternative evotype group who show lower methylation. However, for one small region we see the pattern switching with benign tissue having low methylation, alternative evotype tumours showing high methylation, and other ERG-tumours lying in between.

**Figure S8.** Illustrating, for selected CpGs in *JAK1*, the observed methylation levels in each of the five groups of tumours. Bottom we see the gene, coordinates, and locations of CpGs. Then, moving up, we see violin plots for selected CpGs for the alternative Evotype samples, for DESNT high-risk samples with a detected *TMPRSS2-ERG* fusion, for other samples with a detected *TMPRSS2-ERG* fusion, for other high-risk DESNT samples, and for tumour samples with none of the three characteristics (high-risk DESNT, alternative Evotype, or fusion). Profiles for Benign samples are shown at the top for comparison, and then we see mean values for the five tumour categories and Benign tissue in the top section of the plot. For the plots of the mean profiles, the tumour samples with none of the three characteristics are shown with a narrow black like, while the benign samples have a dashed line. For the violin plots, and mean methylation plot, the CpGs have been equally spaced. Below the first of the violin plots lines indicate the loci in the gene at which the CpGs are situated.

We have a region where tumours with the *TMPRSS2-ERG* fusion track the benign profile and the remaining three groups show higher methylation, while in another region the three DESNT and/or fusion groups showing higher methylation, while the other two now track the benign profile.

**Figure S9.** Illustrating, for selected CpGs in *ZNF536*, the observed methylation levels in each of the five groups of tumours. Bottom we see the gene, coordinates, and locations of CpGs. Then, moving up, we see violin plots for selected CpGs for the alternative Evotype samples, for DESNT high-risk samples with a detected *TMPRSS2-ERG* fusion, for other samples with a detected *TMPRSS2-ERG* fusion, for other high-risk DESNT samples, and for tumour samples with none of the three characteristics (high-risk DESNT, alternative Evotype, or fusion). Profiles for Benign samples are shown at the top for comparison, and then we see mean values for the five tumour categories and Benign tissue in the top section of the plot. For the plots of the mean profiles, the tumour samples with none of the three characteristics are shown with a narrow black like, while the benign samples have a dashed line. For the violin plots, and mean methylation plot, the CpGs have been equally spaced. Below the first of the violin plots lines indicate the loci in the gene at which the CpGs are situated.

In three regions along this gene we see tumours with the fusion tracking the same profile as the benign tissue, while Alternative evotype tumours are the most distinct. In two regions between those, we see a contrast where the fusion positive tumours are most distinct and all other tumour categories track the benign profile.

**Figure S10.** Illustrating, for selected CpGs in *CPNE8*, the observed methylation levels in each of the five groups of tumours. Bottom we see the gene, coordinates, and locations of CpGs. Then, moving up, we see violin plots for selected CpGs for the alternative Evotype samples, for DESNT high-risk samples with a detected *TMPRSS2-ERG* fusion, for other samples with a detected *TMPRSS2-ERG* fusion, for other high-risk DESNT samples, and for tumour samples with none of the three characteristics (high-risk DESNT, alternative Evotype, or fusion). Profiles for Benign samples are shown at the top for comparison, and then we see mean values for the five tumour categories and Benign tissue in the top section of the plot. For the plots of the mean profiles, the tumour samples with none of the three characteristics are shown with a narrow black like, while the benign samples have a dashed line. For the violin plots, and mean methylation plot, the CpGs have been equally spaced. Below the first of the violin plots lines indicate the loci in the gene at which the CpGs are situated.

We see that the ordering of profiles from the prostate categories reverses between the regions depicted.

**Figure S11.** Illustrating, for selected CpGs in *SEPTIN9*, the observed methylation levels in each of the five groups of tumours. Bottom we see the gene, coordinates, and locations of CpGs. Then, moving up, we see violin plots for selected CpGs for the alternative Evotype samples, for DESNT high-risk samples with a detected *TMPRSS2-ERG* fusion, for other samples with a detected *TMPRSS2-ERG* fusion, for other high-risk DESNT samples, and for tumour samples with none of the three characteristics (high-risk DESNT, alternative Evotype, or fusion). Profiles for Benign samples are shown at the top for comparison, and then we see mean values for the five tumour categories and Benign tissue in the top section of the plot. For the plots of the mean profiles, the tumour samples with none of the three characteristics are shown with a narrow black like, while the benign samples have a dashed line. For the violin plots, and mean methylation plot, the CpGs have been equally spaced. Below the first of the violin plots lines indicate the loci in the gene at which the CpGs are situated.

We see that the ordering of profiles from the prostate categories reverses between the regions depicted.

**Figure S12**. Illustrating, for selected CpGs in *FZD10*, the observed methylation levels in each of the five groups of tumours. Bottom we see the gene, coordinates, and locations of CpGs. Then, moving up, we see violin plots for selected CpGs for the alternative Evotype samples, for DESNT high-risk samples with a detected *TMPRSS2-ERG* fusion, for other samples with a detected *TMPRSS2-ERG* fusion, for other high-risk DESNT samples, and for tumour samples with none of the three characteristics (high-risk DESNT, alternative Evotype, or fusion). Profiles for Benign samples are shown at the top for comparison, and then we see mean values for the five tumour categories and Benign tissue in the top section of the plot. For the plots of the mean profiles, the tumour samples with none of the three characteristics are shown with a narrow black like, while the benign samples have a dashed line. For the violin plots, and mean methylation plot, the CpGs have been equally spaced. Below the first of the violin plots lines indicate the loci in the gene at which the CpGs are situated.

We see that the ordering of profiles from the prostate categories changes between the regions depicted. To the left of the figure, only tumours lacking a *TMPRSS2-ERG* fusion show signal, while to the right, the fusion positive and DESNT positive tumours have methylation profiles that exceed those of benign tissue.

**Figure S13.** Illustrating, for selected CpGs in *SMOC2*, the observed methylation levels in each of the five groups of tumours. Bottom we see the gene, coordinates, and locations of CpGs. Then, moving up, we see violin plots for selected CpGs for the alternative Evotype samples, for DESNT high-risk samples with a detected *TMPRSS2-ERG* fusion, for other samples with a detected *TMPRSS2-ERG* fusion, for other high-risk DESNT samples, and for tumour samples with none of the three characteristics (high-risk DESNT, alternative Evotype, or fusion). Profiles for Benign samples are shown at the top for comparison, and then we see mean values for the five tumour categories and Benign tissue in the top section of the plot. For the plots of the mean profiles, the tumour samples with none of the three characteristics are shown with a narrow black like, while the benign samples have a dashed line. For the violin plots, and mean methylation plot, the CpGs have been equally spaced. Below the first of the violin plots lines indicate the loci in the gene at which the CpGs are situated.

The ordering of the profiles from the five tumour categories remains fairly constant for the regions shown. By contrast, the benign tissue profile switches from having the lowest methylation levels to the highest at a point within the gene.

**Figure S14.** Illustrating, for selected CpGs in *PRPSAP1*, the observed methylation levels in each of the five groups of tumours. Bottom we see the gene, coordinates, and locations of CpGs. Then, moving up, we see violin plots for selected CpGs for the alternative Evotype samples, for DESNT high-risk samples with a detected *TMPRSS2-ERG* fusion, for other samples with a detected *TMPRSS2-ERG* fusion, for other high-risk DESNT samples, and for tumour samples with none of the three characteristics (high-risk DESNT, alternative Evotype, or fusion). Profiles for Benign samples are shown at the top for comparison, and then we see mean values for the five tumour categories and Benign tissue in the top section of the plot. For the plots of the mean profiles, the tumour samples with none of the three characteristics are shown with a narrow black like, while the benign samples have a dashed line. For the violin plots, and mean methylation plot, the CpGs have been equally spaced. Below the first of the violin plots lines indicate the loci in the gene at which the CpGs are situated.

The behaviour of the tumour category methylation profiles in this gene are complicated, cautioning against a simple gene-level summary.

**Figure S15**. Illustrating, for selected CpGs in NME5, the observed methylation levels in each of the five groups of tumours. Bottom we see the gene, coordinates, and locations of CpGs. Then, moving up, we see violin plots for selected CpGs for the alternative Evotype samples, for DESNT high-risk samples with a detected *TMPRSS2-ERG* fusion, for other samples with a detected *TMPRSS2-ERG* fusion, for other high-risk DESNT samples, and for tumour samples with none of the three characteristics (high-risk DESNT, alternative Evotype, or fusion). Profiles for Benign samples are shown at the top for comparison, and then we see mean values for the five tumour categories and Benign tissue in the top section of the plot. For the plots of the mean profiles, the tumour samples with none of the three characteristics are shown with a narrow black like, while the benign samples have a dashed line. For the violin plots, and mean methylation plot, the CpGs have been equally spaced. Below the first of the violin plots lines indicate the loci in the gene at which the CpGs are situated.

We see a region of the gene where the Alternative evotype profile shows higher methylation than that of benign tissue, while the *TMPRSS2-ERG* fusion and DESNT cases show a markedly lower level of methylation than is seen in the benign. Summarizing tumour vs benign or ERG+ vs ERG-would risk missing these changes.

**Figure S16.** Illustrating, for selected CpGs in *WNK2*, the observed methylation levels in each of the five groups of tumours. Bottom we see the gene, coordinates, and locations of CpGs. Then, moving up, we see violin plots for selected CpGs for the alternative Evotype samples, for DESNT high-risk samples with a detected *TMPRSS2-ERG* fusion, for other samples with a detected *TMPRSS2-ERG* fusion, for other high-risk DESNT samples, and for tumour samples with none of the three characteristics (high-risk DESNT, alternative Evotype, or fusion). Profiles for Benign samples are shown at the top for comparison, and then we see mean values for the five tumour categories and Benign tissue in the top section of the plot. For the plots of the mean profiles, the tumour samples with none of the three characteristics are shown with a narrow black like, while the benign samples have a dashed line. For the violin plots, and mean methylation plot, the CpGs have been equally spaced. Below the first of the violin plots lines indicate the loci in the gene at which the CpGs are situated.

We see a region of the gene where the Alternative evotype profile shows higher methylation than that of benign tissue, while the *TMPRSS2-ERG* fusion and DESNT cases show a markedly lower level of methylation than is seen in the benign. Summarizing tumour vs benign or ERG+ vs ERG-would risk missing these changes.

**Figure S17.** Illustrating, for selected CpGs in *VPS26C*, the observed methylation levels in each of the five groups of tumours. Bottom we see the gene, coordinates, and locations of CpGs. Then, moving up, we see violin plots for selected CpGs for the alternative Evotype samples, for DESNT high-risk samples with a detected *TMPRSS2-ERG* fusion, for other samples with a detected *TMPRSS2-ERG* fusion, for other high-risk DESNT samples, and for tumour samples with none of the three characteristics (high-risk DESNT, alternative Evotype, or fusion). Profiles for Benign samples are shown at the top for comparison, and then we see mean values for the five tumour categories and Benign tissue in the top section of the plot. For the plots of the mean profiles, the tumour samples with none of the three characteristics are shown with a narrow black like, while the benign samples have a dashed line. For the violin plots, and mean methylation plot, the CpGs have been equally spaced. Below the first of the violin plots lines indicate the loci in the gene at which the CpGs are situated.

We see a short region of the gene where the Alternative evotype profile shows higher methylation than that of benign tissue, while the *TMPRSS2-ERG* fusion and DESNT cases show a markedly lower level of methylation than is seen in the benign. Summarizing tumour vs benign or ERG+ vs ERG-would risk missing these changes.

**Figure S18.** Illustrating, for selected CpGs in *CNNM4*, the observed methylation levels in each of the five groups of tumours. Bottom we see the gene, coordinates, and locations of CpGs. Then, moving up, we see violin plots for selected CpGs for the alternative Evotype samples, for DESNT high-risk samples with a detected *TMPRSS2-ERG* fusion, for other samples with a detected *TMPRSS2-ERG* fusion, for other high-risk DESNT samples, and for tumour samples with none of the three characteristics (high-risk DESNT, alternative Evotype, or fusion). Profiles for Benign samples are shown at the top for comparison, and then we see mean values for the five tumour categories and Benign tissue in the top section of the plot. For the plots of the mean profiles, the tumour samples with none of the three characteristics are shown with a narrow black like, while the benign samples have a dashed line. For the violin plots, and mean methylation plot, the CpGs have been equally spaced. Below the first of the violin plots lines indicate the loci in the gene at which the CpGs are situated.

We see two regions of the gene where the Alternative evotype profile largely tracks that of benign tissue, while the *TMPRSS2-ERG* fusion cases show a substantial reduction in methylation. There is a suggestion that canonical evotype cases that are not fusion positive show a profile slightly lower than that of benign tissue.

**Figure S19.** Illustrating, for selected CpGs in *CPNE4*, the observed methylation levels in each of the five groups of tumours. Bottom we see the gene, coordinates, and locations of CpGs. Then, moving up, we see violin plots for selected CpGs for the alternative Evotype samples, for DESNT high-risk samples with a detected *TMPRSS2-ERG* fusion, for other samples with a detected *TMPRSS2-ERG* fusion, for other high-risk DESNT samples, and for tumour samples with none of the three characteristics (high-risk DESNT, alternative Evotype, or fusion). Profiles for Benign samples are shown at the top for comparison, and then we see mean values for the five tumour categories and Benign tissue in the top section of the plot. For the plots of the mean profiles, the tumour samples with none of the three characteristics are shown with a narrow black like, while the benign samples have a dashed line. For the violin plots, and mean methylation plot, the CpGs have been equally spaced. Below the first of the violin plots lines indicate the loci in the gene at which the CpGs are situated.

We see that, for most of the CpGs showm, the tumour profiles separate out into the Alternative evotype cases closest to the benign profile, the two DESNT high-risk categories furthest from the benign profule, and the other two Canonical evotype categories in between. The clustering of profiles by DESNT status rather than by *TMPRSS2-ERG* fusion status is a behavioural change from earlier examples.

**Figure S20.** Illustrating, for selected CpGs in *NKAIN1*, the observed methylation levels in each of the five groups of tumours. Bottom we see the gene, coordinates, and locations of CpGs. Then, moving up, we see violin plots for selected CpGs for the alternative Evotype samples, for DESNT high-risk samples with a detected *TMPRSS2-ERG* fusion, for other samples with a detected *TMPRSS2-ERG* fusion, for other high-risk DESNT samples, and for tumour samples with none of the three characteristics (high-risk DESNT, alternative Evotype, or fusion). Profiles for Benign samples are shown at the top for comparison, and then we see mean values for the five tumour categories and Benign tissue in the top section of the plot. For the plots of the mean profiles, the tumour samples with none of the three characteristics are shown with a narrow black like, while the benign samples have a dashed line. For the violin plots, and mean methylation plot, the CpGs have been equally spaced. Below the first of the violin plots lines indicate the loci in the gene at which the CpGs are situated.

We see that, for most of the CpGs showm, the tumour profiles separate out into the Alternative evotype cases closest to the benign profile, the two DESNT high-risk categories furthest from the benign profule, and the other two Canonical evotype categories in between. The clustering of profiles by DESNT status rather than by *TMPRSS2-ERG* fusion status is a behavioural change from earlier examples.

**Figure S21.** Illustrating, for selected CpGs in *CDH4*, the observed methylation levels in each of the five groups of tumours. Bottom we see the gene, coordinates, and locations of CpGs. Then, moving up, we see violin plots for selected CpGs for the alternative Evotype samples, for DESNT high-risk samples with a detected *TMPRSS2-ERG* fusion, for other samples with a detected *TMPRSS2-ERG* fusion, for other high-risk DESNT samples, and for tumour samples with none of the three characteristics (high-risk DESNT, alternative Evotype, or fusion). Profiles for Benign samples are shown at the top for comparison, and then we see mean values for the five tumour categories and Benign tissue in the top section of the plot. For the plots of the mean profiles, the tumour samples with none of the three characteristics are shown with a narrow black like, while the benign samples have a dashed line. For the violin plots, and mean methylation plot, the CpGs have been equally spaced. Below the first of the violin plots lines indicate the loci in the gene at which the CpGs are situated.

An example where the Alternative evotype shows greatest difference to benign tissue.

**Figure S22.** Illustrating, for selected CpGs in *SPOP*, the observed methylation levels by tumour grade group (GG1 *n* = 9, GG2 *n* = 113, GG3 *n* = 44, GG4 *n* = 7, GG5 *n* = 9). Bottom we see the gene, coordinates, and locations of CpGs. Then, moving up, we see violin plots for selected CpGs for the five grade groups. Profiles for Benign samples are shown at the top for comparison, and then we see mean values for the five tumour categories and Benign tissue in the top section of the plot. For the violin plots, and mean methylation plot, the CpGs have been equally spaced. Below the first of the violin plots lines indicate the loci in the gene at which the CpGs are situated.

It is notable that categorizing by grade group highlights apparent differences between tumour and benign tissue, but hides the inter-tumour heterogeneity we saw in Figure 3.

**Figure S23.** Illustrating, for selected CpGs in *AJAP1*, the observed methylation levels by tumour grade group (GG1 *n* = 9, GG2 *n* = 113, GG3 *n* = 44, GG4 *n* = 7, GG5 *n* = 9). Bottom we see the gene, coordinates, and locations of CpGs. Then, moving up, we see violin plots for selected CpGs for the five grade groups. Profiles for Benign samples are shown at the top for comparison, and then we see mean values for the five tumour categories and Benign tissue in the top section of the plot. For the violin plots, and mean methylation plot, the CpGs have been equally spaced. Below the first of the violin plots lines indicate the loci in the gene at which the CpGs are situated.

Categorizing by grade group hides the inter-tumour heterogeneity we saw in Supplementary Figure S7. While Grade group 4 seems to show some departure from the rest, this is the smallest set of tumours and should be treated with caution.

**Figure S24.** Illustrating, for selected CpGs in *JAK1*, the observed methylation levels by tumour grade group. Bottom we see the gene, coordinates, and locations of CpGs. Then, moving up, we see violin plots for selected CpGs for the five grade groups. Profiles for Benign samples are shown at the top for comparison, and then we see mean values for the five tumour categories and Benign tissue in the top section of the plot. For the violin plots, and mean methylation plot, the CpGs have been equally spaced. Below the first of the violin plots lines indicate the loci in the gene at which the CpGs are situated.

While variation is apparent, and some tumours differ from benign, there is no obvious trend due to Grade group.

**Figure S25.** Illustrating, for selected CpGs in *ZNF536*, the observed methylation levels by tumour grade group. Bottom we see the gene, coordinates, and locations of CpGs. Then, moving up, we see violin plots for selected CpGs for the five grade groups. Profiles for Benign samples are shown at the top for comparison, and then we see mean values for the five tumour categories and Benign tissue in the top section of the plot. For the violin plots, and mean methylation plot, the CpGs have been equally spaced. Below the first of the violin plots lines indicate the loci in the gene at which the CpGs are situated.

There is a suggestion that higher grade groups separate more from the benign methylation profiles, but the trend is not clear.

**Figure S26.** Illustrating, for selected CpGs in *CPNE8*, the observed methylation levels by tumour grade group. Bottom we see the gene, coordinates, and locations of CpGs. Then, moving up, we see violin plots for selected CpGs for the five grade groups. Profiles for Benign samples are shown at the top for comparison, and then we see mean values for the five tumour categories and Benign tissue in the top section of the plot. For the violin plots, and mean methylation plot, the CpGs have been equally spaced. Below the first of the violin plots lines indicate the loci in the gene at which the CpGs are situated.

**Figure S27.** Illustrating, for selected CpGs in *SEPTIN9*, the observed methylation levels by tumour grade group. Bottom we see the gene, coordinates, and locations of CpGs. Then, moving up, we see violin plots for selected CpGs for the five grade groups. Profiles for Benign samples are shown at the top for comparison, and then we see mean values for the five tumour categories and Benign tissue in the top section of the plot. For the violin plots, and mean methylation plot, the CpGs have been equally spaced. Below the first of the violin plots lines indicate the loci in the gene at which the CpGs are situated.

**Figure S28.** Illustrating, for selected CpGs in *FZD10*, the observed methylation levels by tumour grade group. Bottom we see the gene, coordinates, and locations of CpGs. Then, moving up, we see violin plots for selected CpGs for the five grade groups. Profiles for Benign samples are shown at the top for comparison, and then we see mean values for the five tumour categories and Benign tissue in the top section of the plot. For the violin plots, and mean methylation plot, the CpGs have been equally spaced. Below the first of the violin plots lines indicate the loci in the gene at which the CpGs are situated.

**Figure S29.** Illustrating, for selected CpGs in *SMOC2*, the observed methylation levels by tumour grade group. Bottom we see the gene, coordinates, and locations of CpGs. Then, moving up, we see violin plots for selected CpGs for the five grade groups. Profiles for Benign samples are shown at the top for comparison, and then we see mean values for the five tumour categories and Benign tissue in the top section of the plot. For the violin plots, and mean methylation plot, the CpGs have been equally spaced. Below the first of the violin plots lines indicate the loci in the gene at which the CpGs are situated.

**Figure S30.** Illustrating, for selected CpGs in *PRPSAP1*, the observed methylation levels by tumour grade group. Bottom we see the gene, coordinates, and locations of CpGs. Then, moving up, we see violin plots for selected CpGs for the five grade groups. Profiles for Benign samples are shown at the top for comparison, and then we see mean values for the five tumour categories and Benign tissue in the top section of the plot. For the violin plots, and mean methylation plot, the CpGs have been equally spaced. Below the first of the violin plots lines indicate the loci in the gene at which the CpGs are situated.

**Figure S31.** Illustrating, for selected CpGs in *NME5*, the observed methylation levels by tumour grade group. Bottom we see the gene, coordinates, and locations of CpGs. Then, moving up, we see violin plots for selected CpGs for the five grade groups. Profiles for Benign samples are shown at the top for comparison, and then we see mean values for the five tumour categories and Benign tissue in the top section of the plot. For the violin plots, and mean methylation plot, the CpGs have been equally spaced. Below the first of the violin plots lines indicate the loci in the gene at which the CpGs are situated.

**Figure S32.** Illustrating, for selected CpGs in *WNK2*, the observed methylation levels by tumour grade group. Bottom we see the gene, coordinates, and locations of CpGs. Then, moving up, we see violin plots for selected CpGs for the five grade groups. Profiles for Benign samples are shown at the top for comparison, and then we see mean values for the five tumour categories and Benign tissue in the top section of the plot. For the violin plots, and mean methylation plot, the CpGs have been equally spaced. Below the first of the violin plots lines indicate the loci in the gene at which the CpGs are situated.

**Figure S33.** Illustrating, for selected CpGs in *VPS26C*, the observed methylation levels by tumour grade group. Bottom we see the gene, coordinates, and locations of CpGs. Then, moving up, we see violin plots for selected CpGs for the five grade groups. Profiles for Benign samples are shown at the top for comparison, and then we see mean values for the five tumour categories and Benign tissue in the top section of the plot. For the violin plots, and mean methylation plot, the CpGs have been equally spaced. Below the first of the violin plots lines indicate the loci in the gene at which the CpGs are situated.

**Figure S34.** Illustrating, for selected CpGs in *CNNM4*, the observed methylation levels by tumour grade group. Bottom we see the gene, coordinates, and locations of CpGs. Then, moving up, we see violin plots for selected CpGs for the five grade groups. Profiles for Benign samples are shown at the top for comparison, and then we see mean values for the five tumour categories and Benign tissue in the top section of the plot. For the violin plots, and mean methylation plot, the CpGs have been equally spaced. Below the first of the violin plots lines indicate the loci in the gene at which the CpGs are situated.

**Figure S35.** Illustrating, for selected CpGs in *CPNE4*, the observed methylation levels by tumour grade group. Bottom we see the gene, coordinates, and locations of CpGs. Then, moving up, we see violin plots for selected CpGs for the five grade groups. Profiles for Benign samples are shown at the top for comparison, and then we see mean values for the five tumour categories and Benign tissue in the top section of the plot. For the violin plots, and mean methylation plot, the CpGs have been equally spaced. Below the first of the violin plots lines indicate the loci in the gene at which the CpGs are situated.

**Figure S36.** Illustrating, for selected CpGs in *NKAIN1*, the observed methylation levels by tumour grade group. Bottom we see the gene, coordinates, and locations of CpGs. Then, moving up, we see violin plots for selected CpGs for the five grade groups. Profiles for Benign samples are shown at the top for comparison, and then we see mean values for the five tumour categories and Benign tissue in the top section of the plot. For the violin plots, and mean methylation plot, the CpGs have been equally spaced. Below the first of the violin plots lines indicate the loci in the gene at which the CpGs are situated.

**Figure S37.** Illustrating, for selected CpGs in *CDH4*, the observed methylation levels by tumour grade group. Bottom we see the gene, coordinates, and locations of CpGs. Then, moving up, we see violin plots for selected CpGs for the five grade groups. Profiles for Benign samples are shown at the top for comparison, and then we see mean values for the five tumour categories and Benign tissue in the top section of the plot. For the violin plots, and mean methylation plot, the CpGs have been equally spaced. Below the first of the violin plots lines indicate the loci in the gene at which the CpGs are situated.

*CDH4* did not show obvious trends in terms of the five categories used above S21, but does show a clear trend in terms of grade group.

**Figure S38.** Flowchart showing the filtering steps that reduce the GWAS catalog hits to the 74 examined in this study.

**Figure S39.** External evidence of methylation and expression interactions for the gene *MMP7*. A) For various cell line experiments for which the data are publicly accessible, we show the distribution across all genes of standardized changes in expression resulting from methylation silencing via 5-Azacytidine. The genes *MMP7* and *FOXA2* are highlighted. Plots are grouped by the cancer/tissue the cell lines represent. B) From the TCGA data set, for the three Illumina 450k methylation probes in the region (locations indicated in the schematic above), associations of methylation level with expression of *MMP7*.

**Figure S40.** *MMP7* Expression in the UK Prostate ICGC cohort. Expression is given as log 2 RPM. A) Histogram of tumour *MMP7* expression levels in the cohort. Four of the bins are colour coded for later reference. B) Boxplot of tumour *MMP7* expression by haplogroup (defined in Figure 4). C) Correlation of tumour expression and standardlzed methylation [standardized methylation defined as the average over eight CpGs, each of which has been transformed to have mean methylation of 0 and standard deviation of 1]. D) Methylation patterns in RR and RS haplogroups for nine CpGs are still visible when restricting to men with expression levels between 11 and 12. Patterns that are individually statistically significant are indicated with an asterisk. E) – G) As for D, but with different tranches of *MMP7* expression. Methylation levels in the CpG corresponding to the SNP of interest were not subject to a hypothesis test. 32 out of 32 comparisons go in the same direction: The methylation association with genotype is still apparent when restricting expression levels, strongly suggesting that expression does not drive the change. There are insufficient numbers in the SS haplogroup to include that group in this analysis.

**Figure S41.** Associations between genomic variants and methylation patterns in the *MMP7* gene. As Figure 4, but showing patterns in tumour samples split by *TMPRSS2-ERG* fusion.

**Figure S42.** Associations between genomic variants and methylation patterns in the *MMP7* gene. As Figure 4, but showing patterns in tumour samples split by Evotype status.

**Figure S43.** Associations between genomic variants and methylation patterns in the *MMP7* gene. As Figure 4, but showing patterns for all five haplogroups.

**Figure S44.** Associations between genomic variants and methylation patterns in the *MMP7* gene. As Figure 4, but showing patterns in tumour samples split by rs11568818.

**Figure S45.** Associations between genomic variants and methylation patterns in the *MMP7* gene. As Figure 4, but showing patterns in tumour samples split by rs11568819.

**Figure S46.** Associations between the rs1058319 variant and methylation patterns in the *SLC2A4RG* gene. This SNP does not create or remove a CpG. Top: A schematic of the genomic region indicating the region plotted below. 2nd) For 26 CpGs in the region, the distribution of methylation levels in Benign samples is plotted against the distance of the CpG from the SNP in question. 3rd) For 26 CpGs in the region, the distribution of methylation levels in Tumour samples is plotted against the distance of the CpG from the SNP in question. Adjusted *−* log_10_ p-values are indicated when these exceed 2 (e.g. *p <* 0.01). Bottom: For 15 CpGs where it is possible directly to phase methylation and genotype (i.e. those within 150 bases of the SNP) we present a meta-analysis of relative risk of a molecule being methylated given the allele at rs1058319, calculated across heterozygous men. Again, where adjusted *−* log_10_ p-values exceed 2, these are indicated. We note that the effects are stronger in tumour tissue than benign.

**Figure S47.** Associations between the rs5759167 variant and methylation patterns between the *TTLL1* and *BIK* genes. This SNP does not create or remove a CpG. Top: A schematic of the genomic region indicating the region plotted below. 2nd) For 17 CpGs in the region, the distribution of methylation levels in Benign samples is plotted against the distance of the CpG from the SNP in question. Adjusted *−* log_10_ p-values are indicated when these exceed 2 (e.g. *p <* 0.01). 3rd) For 17 CpGs in the region, the distribution of methylation levels in Tumour samples is plotted against the distance of the CpG from the SNP in question. Adjusted *−* log_10_ p-values are indicated when these exceed 2 (e.g. *p <* 0.01). Bottom: For 8 CpGs where it is possible directly to phase methylation and genotype (i.e. those within 150 bases of the SNP) we present a meta-analysis of relative risk of a molecule being methylated given the allele at rs5759167, calculated across heterozygous men. Again, where adjusted *−* log_10_ p-values exceed 2, these are indicated. We note that the effects are stronger in tumour tissue than benign.

**Figure S48.** Associations between the rs10866527 variant at chr5:1891800 and methylation patterns in the local region. This SNP creates a CpG when it takes the C allele, but in our cohort rs10866528 provides a CpG twenty bases away when rs10866527 takes the T allele. Effectively then, this variant moves a CpG 20 bases. Top: For 22 CpGs in the region, the distribution of methylation levels in Benign samples is plotted against the distance of the CpG from the SNP in question. 2nd: For 22 CpGs in the region, the distribution of methylation levels in Tumour samples is plotted against the distance of the CpG from the SNP in question. Adjusted *−* log_10_ p-values are indicated when these exceed 2 (e.g. *p <* 0.01). 3rd: For 13 CpGs where it is possible directly to phase methylation and genotype (i.e. those within 150 bases of the SNP, excluding the CpG created by the SNP) we present a meta-analysis of relative risk of a molecule being methylated given the allele at rs10866527, calculated across heterozygous men. Again, where adjusted *−* log_10_ p-values exceed 2, these are indicated. We note that the effects are present only in tumour tissue. Bottom: Showing that for the cohort, the overall methylation levels don’t change between tumour and benign, but that the effect of the SNP shown above results in contrasting tumour-benign differences in men who are homozygous at this SNP.

**Figure S49.** Associations between the rs5919393 variant and methylation patterns in the *AR* gene. This SNP does not create or remove a CpG. The Androgen Receptor gene is on the X chromosome, meaning that we cannot look within heterozygous men in this case. Also, this SNP has relatively low allele frequency, and there are relatively few CpGs in the region. Despite the effects this has on our power to detect associations, the importance of the *AR* gene to prostate cancer make the results worth considering. Top: A schematic of the genomic region around rs5919393. 2nd) For 3 CpGs in the region, the distribution of methylation levels in Benign samples is plotted against the distance of the CpG from the SNP in question. Bottom) For 3 CpGs in the region, the distribution of methylation levels in Tumour samples is plotted against the distance of the CpG from the SNP in question. Adjusted *−* log_10_ p-values are indicated when these exceed 2 (e.g. *p <* 0.01). We note that while only one CpG in Tumour tissue showed a significant association, the same trend is seen in all three CpGs in both tissues.

**Figure S50.** Associations between the rs5919393 variant and methylation patterns in the *TERT* gene. This SNP does not creates a CpG in a CpGpCpG. Top: A schematic of the genomic region. 2nd) For 13 CpGs in the region, the distribution of methylation levels in Benign samples is plotted against the distance of the CpG from the SNP in question. 3rd) For 13 CpGs in the region, the distribution of methylation levels in Tumour samples is plotted against the distance of the CpG from the SNP in question. Bottom: For 12 CpGs where it is possible directly to phase methylation and genotype (i.e. those within 150 bases of the SNP, excluding the CpG created by the SNP) we present a meta-analysis of relative risk of a molecule being methylated given the allele at rs5919393, calculated across heterozygous men. Again, where adjusted *−* log_10_ p-values exceed 2, these are indicated. The only obvious effect is in the CpG neighbouring that created by the SNP, but effect goes in the opposite direction to that for the CpG created by the SNP.

**Figure S51.** Hemimethylation and transcription. Hemimethylation affects expression of genes sharing promoter. Expression ratio of bidirectional promoter genes (Y-axis) and hemimethylation of gene bodies of the genes grouped by magnitude of absolute hemimethylation (*|HM| <* 10%, 10% *≤ |HM| <* 20%, 20% *≤ |HM| <* 30%, 30% *≤ |HM| <* 40%, 40% *≤ |HM| <* 50%, 50% *≤ |HM| <* 60%, 60% *≤ |HM| <* 70%, 70% *≤ |HM| <* 80%, 80% *≤ |HM| <* 90%, 90% *≤ |HM|*).

**Figure S52.** Hemimethylation and recurrence. (A) Biochemical recurrence in patients showing very-high hemimethylation (95th quantile, violet) versus the rest of the cohort hemimethylation (yellow). (B) Metastasis in patients showing very-high hemimethylation (95th quantile, violet) versus the rest of the cohort hemimethylation (yellow)

**Figure S53.** Checking the robustness of the results relating to the prognostic value of the number of risk groups. A) The Kaplan–Meier curve for time to biochemical recurrence from Figure 7 is reproduced. B) Three exponential models are compared (in order: a model with no group effect, a model with independent survival estimates in each group, and a model where the log-survival-time increases uniformly with the number of risk groups a man is a member of). C) Survival curves obtained from fitting the second model, where there are independent survival estimates in each group. D) Survival curves obtained from fitting the third model, where there the decrease from group to group is constrained. E) Bootstrap estimates of survival time for the different numbers of risk classes a man is a member of using the second model. F) Bootstrap estimates of survival time for the different numbers of risk classes a man is a member of using the third model.

**Figure S54.** A. For the exponential model where the log-survival-time increases uniformly with the number of risk groups a man is a member of (seen in Supplementary Figure S53), we show the effect of adding covariates Grade Group and Age on the parameter of the model. B. We show Kaplan–Meier curves calculated from men in Grade Group 2 only. While the reduction in the number of men reduces power, the basic pattern is apparent.

**Figure S55.** Copy number plots for the tumour sample of Man 0045 generated from 11628 common A*>*T SNPs that will not be affected by bisulfite conversion. Top: Segmented plot of log-ratios of the tumour sample of Man 0160 relative to unrelated benign samples in the same library preparation batch (created using the DNACopy R package). Bottom: Showing the allele fractions observed in the tumour tissue for SNPs that are germline heterozygous (allele fraction between 0.2 and 0.8 in the benign tissue).

**Figure S56.** Copy number plots for the benign sample of Man 0045 generated from 11628 common A*>*T SNPs that will not be affected by bisulfite conversion. Top: Segmented plot of log-ratios of the benign sample of Man 0160 relative to unrelated benign samples in the same library preparation batch (created using the DNACopy R package). Bottom: Showing the allele fractions observed in the benign tissue for SNPs that are germline heterozygous (allele fraction between 0.2 and 0.8 in the benign tissue).

**Figure S57.** Copy number plots for the tumour sample of Man 0160 generated from 11628 common A*>*T SNPs that will not be affected by bisulfite conversion. Top: Segmented plot of log-ratios of the tumour sample of Man 0160 relative to unrelated benign samples in the same library preparation batch (created using the DNACopy R package). Bottom: Showing the allele fractions observed in the tumour tissue for SNPs that are germline heterozygous (allele fraction between 0.2 and 0.8 in the benign tissue).

**Figure S58.** Copy number plots for the benign sample of Man 0160 generated from 11628 common A*>*T SNPs that will not be affected by bisulfite conversion. Top: Segmented plot of log-ratios of the benign sample of Man 0160 relative to unrelated benign samples in the same library preparation batch (created using the DNACopy R package). Bottom: Showing the allele fractions observed in the benign tissue for SNPs that are germline heterozygous (allele fraction between 0.2 and 0.8 in the benign tissue).

**Figure S59.** Copy number plots for the tumour sample of Man 0222 generated from 11628 common A*>*T SNPs that will not be affected by bisulfite conversion. Top: Segmented plot of log-ratios of the tumour sample of Man 0222 relative to unrelated benign samples in the same library preparation batch (created using the DNACopy R package). Bottom: Showing the allele fractions observed in the tumour tissue for SNPs that are germline heterozygous (allele fraction between 0.2 and 0.8 in the benign tissue).

**Figure S60.** Copy number plots for the benign sample of Man 0222 generated from 11628 common A*>*T SNPs that will not be affected by bisulfite conversion. Top: Segmented plot of log-ratios of the benign sample of Man 0222 relative to unrelated benign samples in the same library preparation batch (created using the DNACopy R package). Bottom: Showing the allele fractions observed in the benign tissue for SNPs that are germline heterozygous (allele fraction between 0.2 and 0.8 in the benign tissue).

**Figure S61.** Copy number plots for the tumour sample of Man 0202 generated from 11628 common A*>*T SNPs that will not be affected by bisulfite conversion. Top: Segmented plot of log-ratios of the tumour sample of Man 0202 relative to unrelated benign samples in the same library preparation batch (created using the DNACopy R package). Bottom: Showing the allele fractions observed in the tumour tissue for SNPs that are germline heterozygous (allele fraction between 0.2 and 0.8 in the benign tissue).

**Figure S62.** Copy number plots for the benign sample of Man 0202 generated from 11628 common A*>*T SNPs that will not be affected by bisulfite conversion. Top: Segmented plot of log-ratios of the benign sample of Man 0202 relative to unrelated benign samples in the same library preparation batch (created using the DNACopy R package). Bottom: Showing the allele fractions observed in the benign tissue for SNPs that are germline heterozygous (allele fraction between 0.2 and 0.8 in the benign tissue).

