## SupplementaryFigures for "Novel Methylation Markers in a Prostate Cancer Cohort are Associated with Disease Development and Relapse"

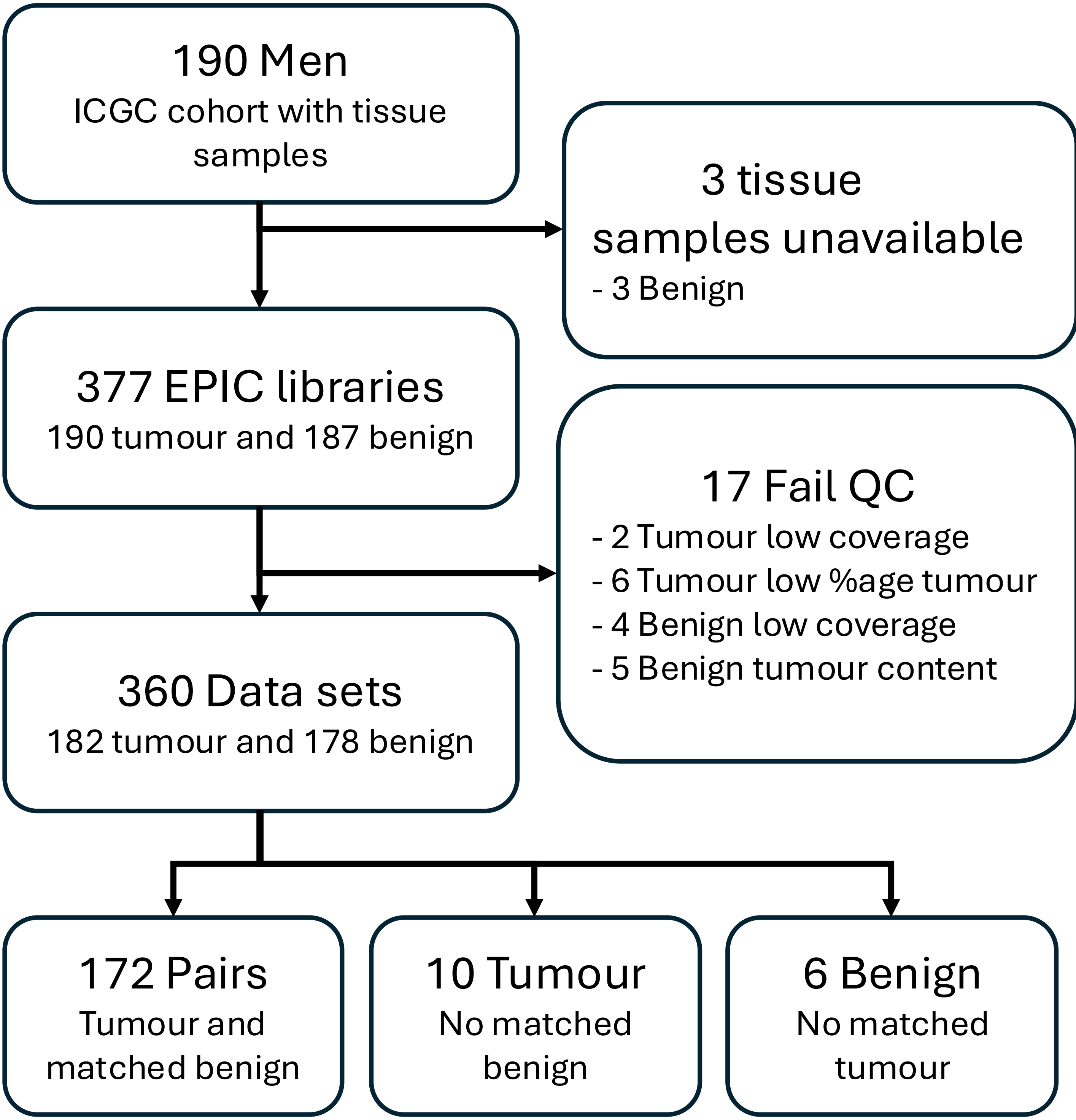


Supp Figure 1

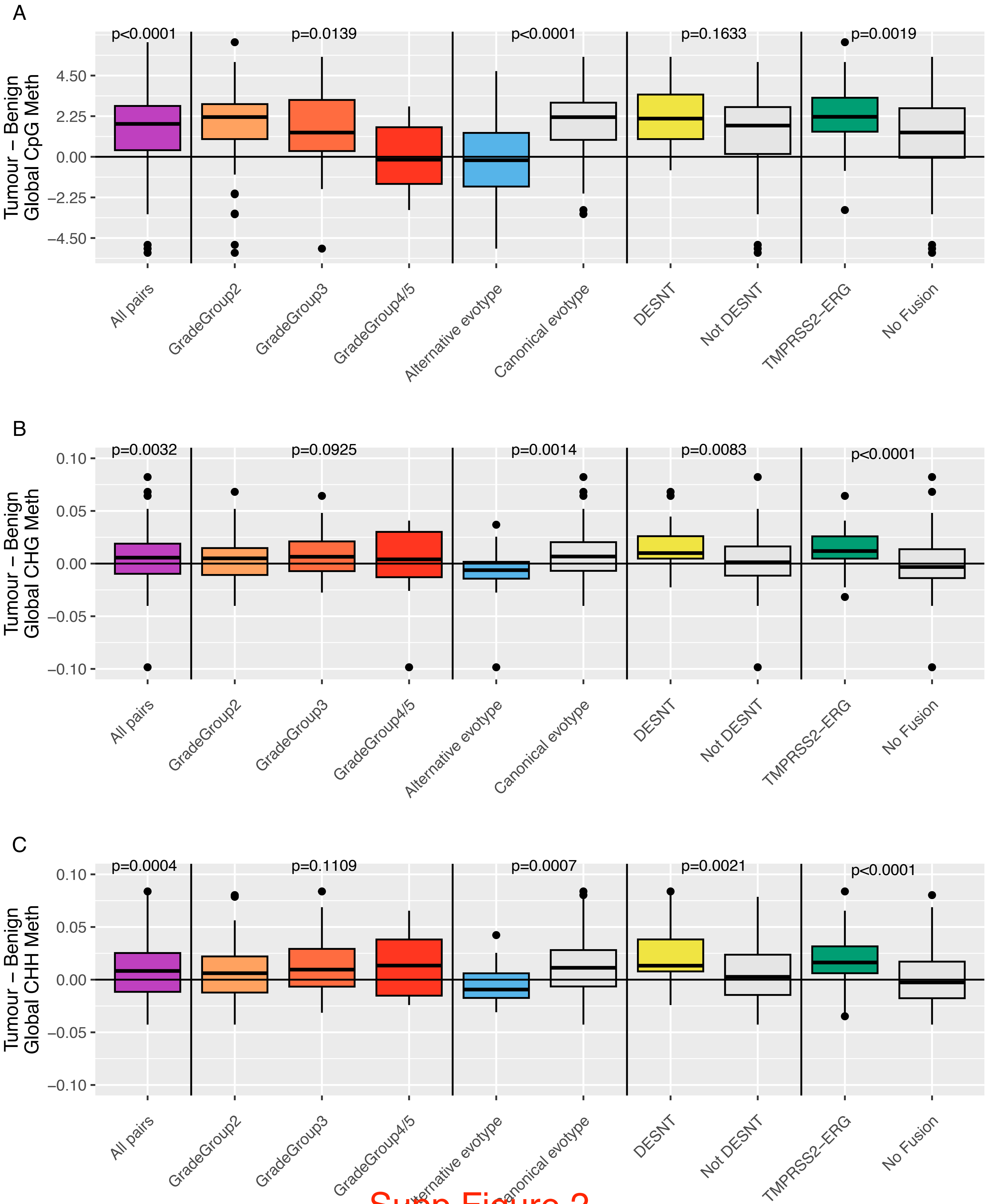


Supp Figure 2

A

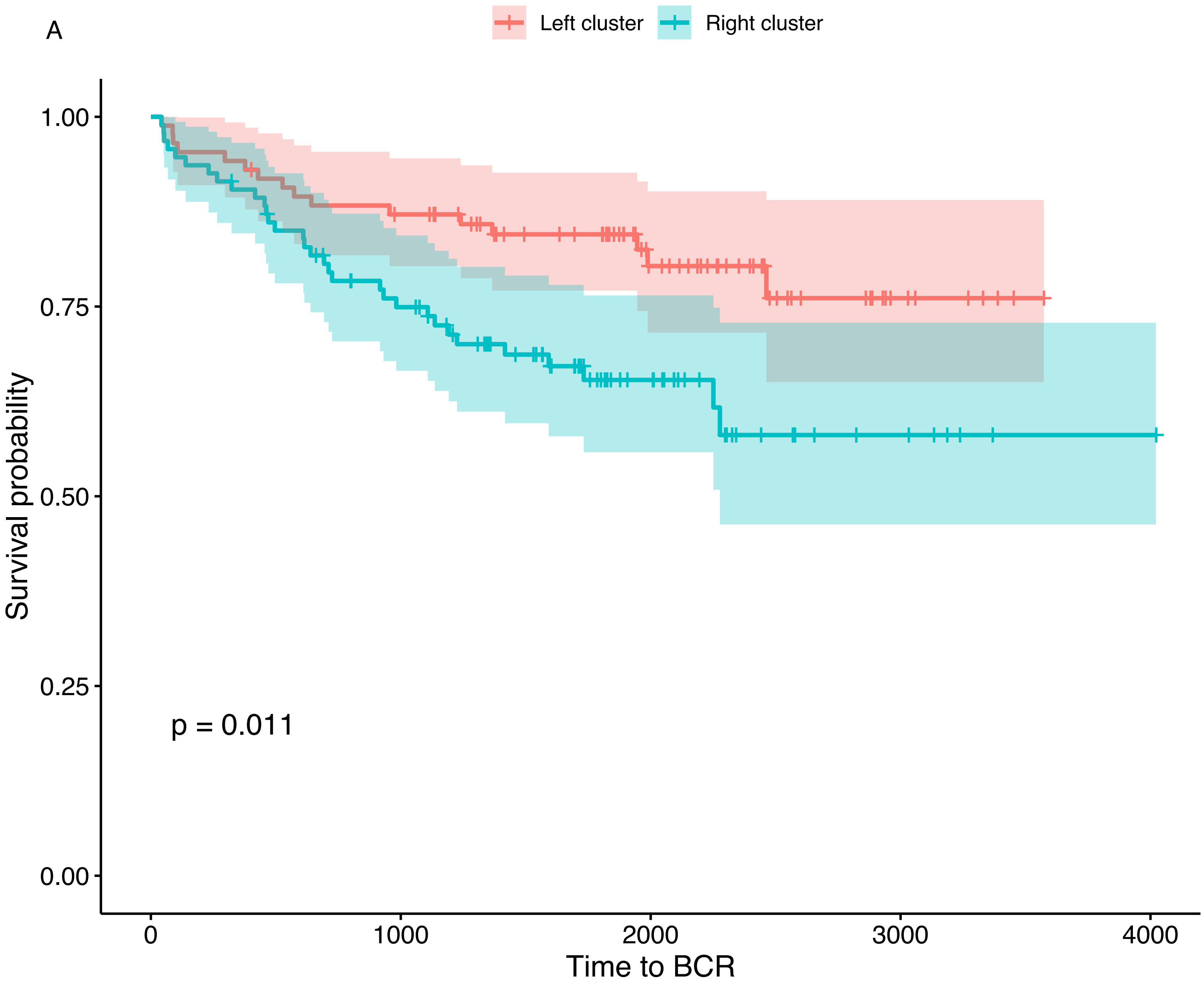


B

| | 1 | 2 | 3 | 4 | 5 |
|---|---|---|---|---|---|
| *Left* | 8 | 70.1 | 12.6 | 2.3 | 6.9 |
| *Right* | 2.1 | 54.7 | 34.7 | 5.3 | 3.2 |

Grade Group % by cluster (p=0.0015)

| | Alternative | Canonical |
|---|---|---|
| *Left* | 7.1 | 92.9 |
| *Right* | 28.2 | 71.8 |

Evotype % by cluster (p=0.0022)

| | DESNT | NOT DESNT |
|---|---|---|
| *Left* | 11.6 | 88.4 |
| *Right* | 28.3 | 71.7 |

DESNT % by cluster (p=0.0084)

| | NO FUSION | TMPRSS2−ERG |
|---|---|---|
| *Left* | 54.9 | 45.1 |
| *Right* | 68.1 | 31.9 |

Fusion % by cluster (p=0.0873)

Supp Figure 3

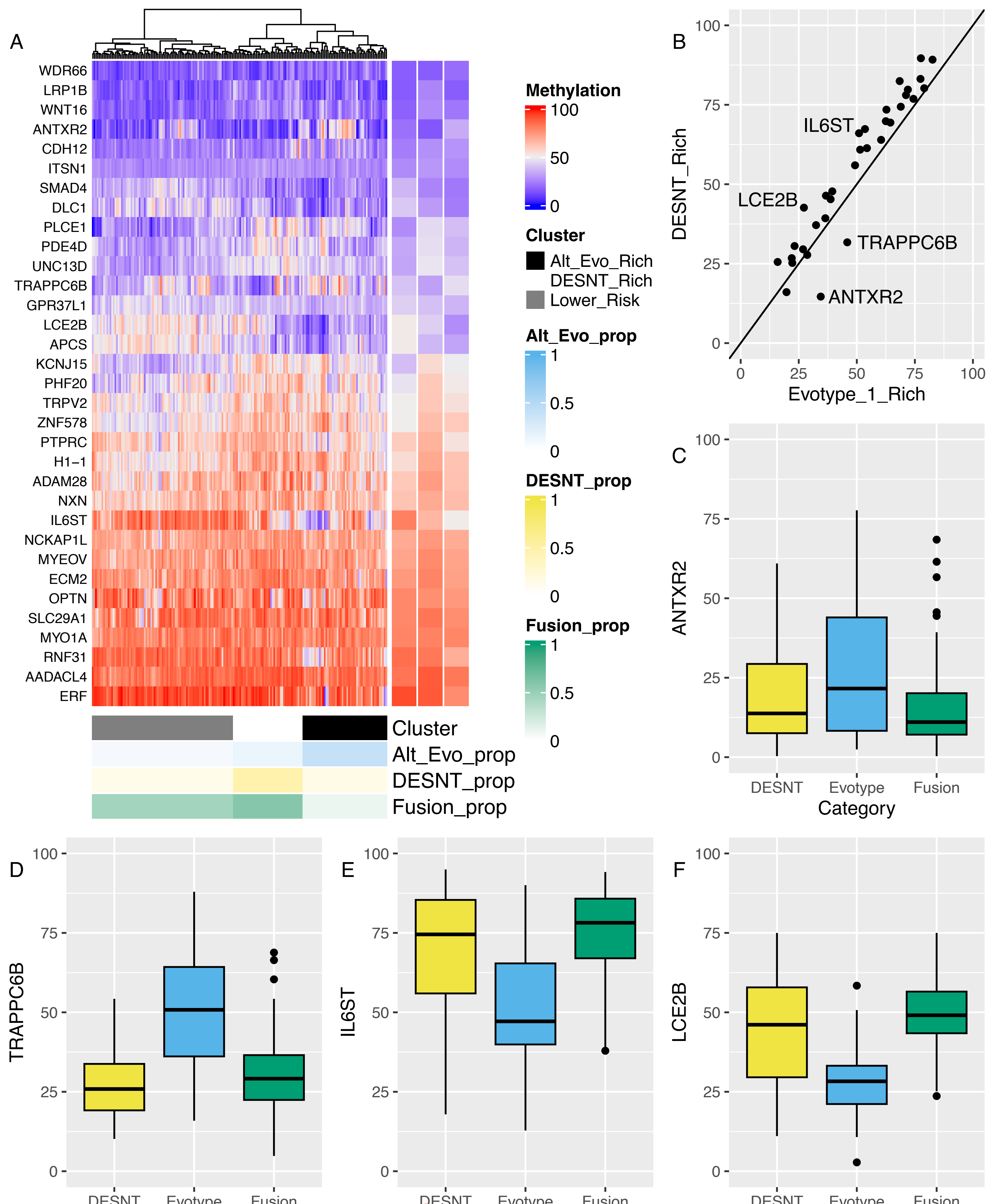


Supp Figure 4

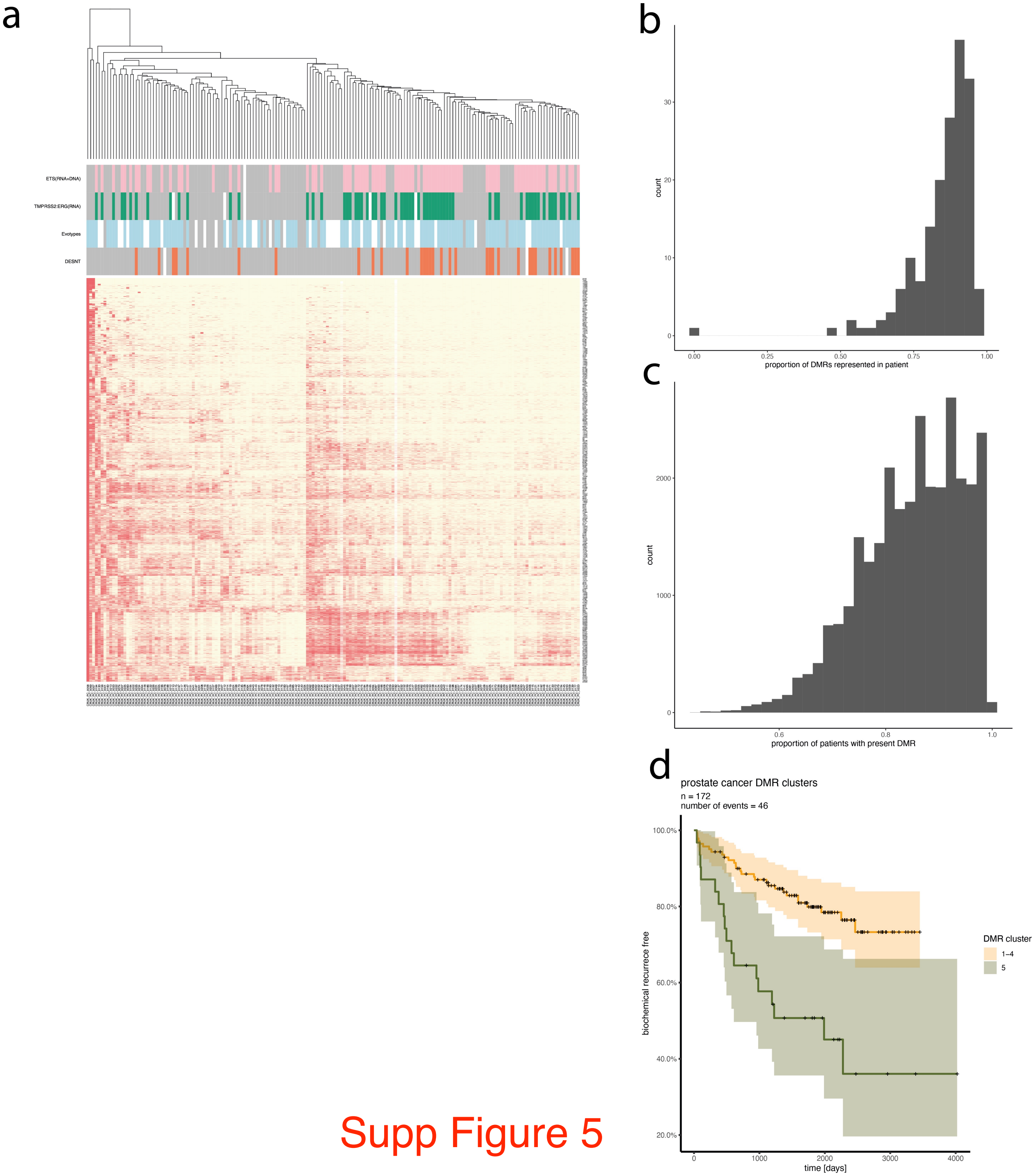


Supp Figure 5

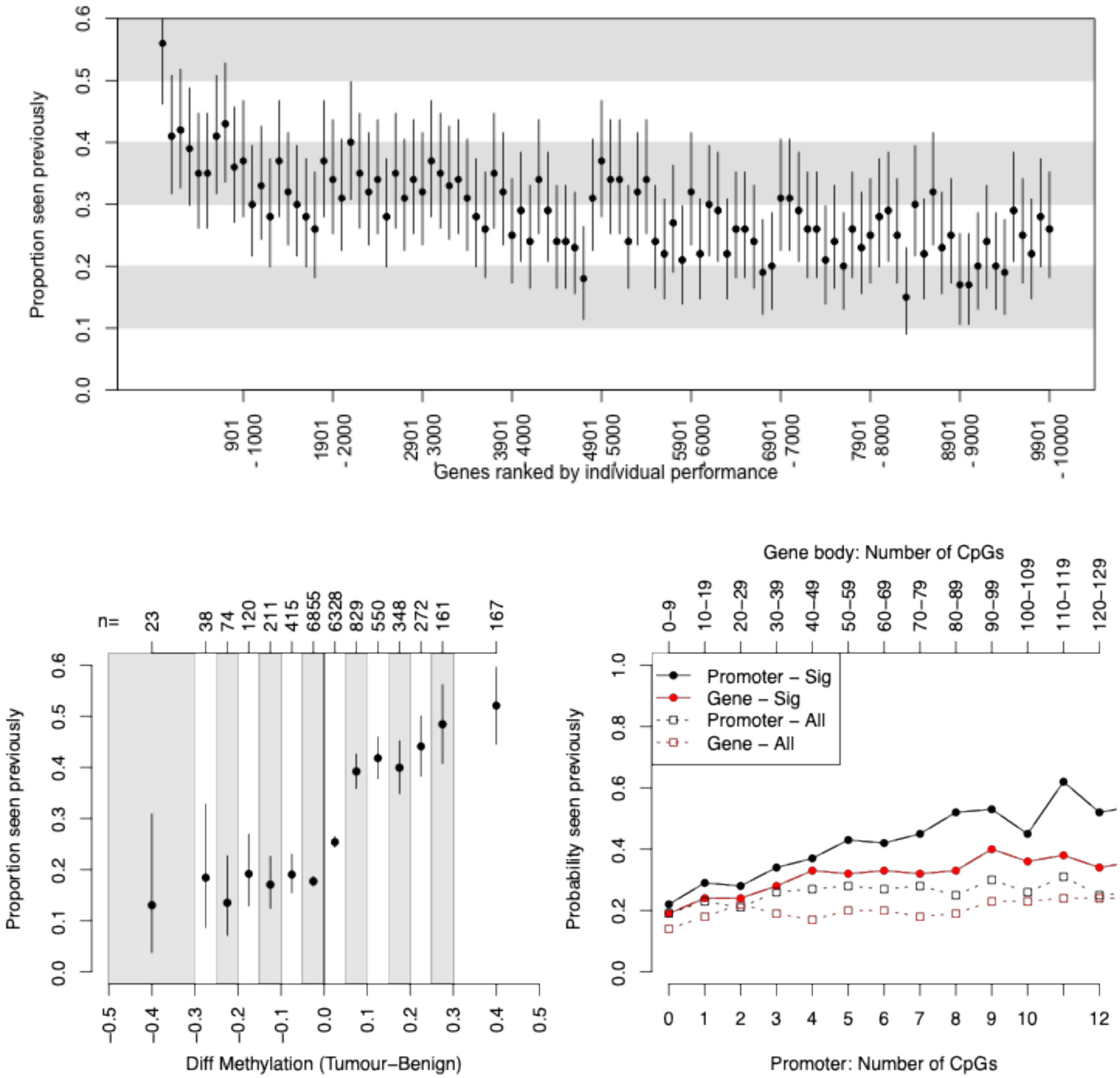


Supp Figure 6

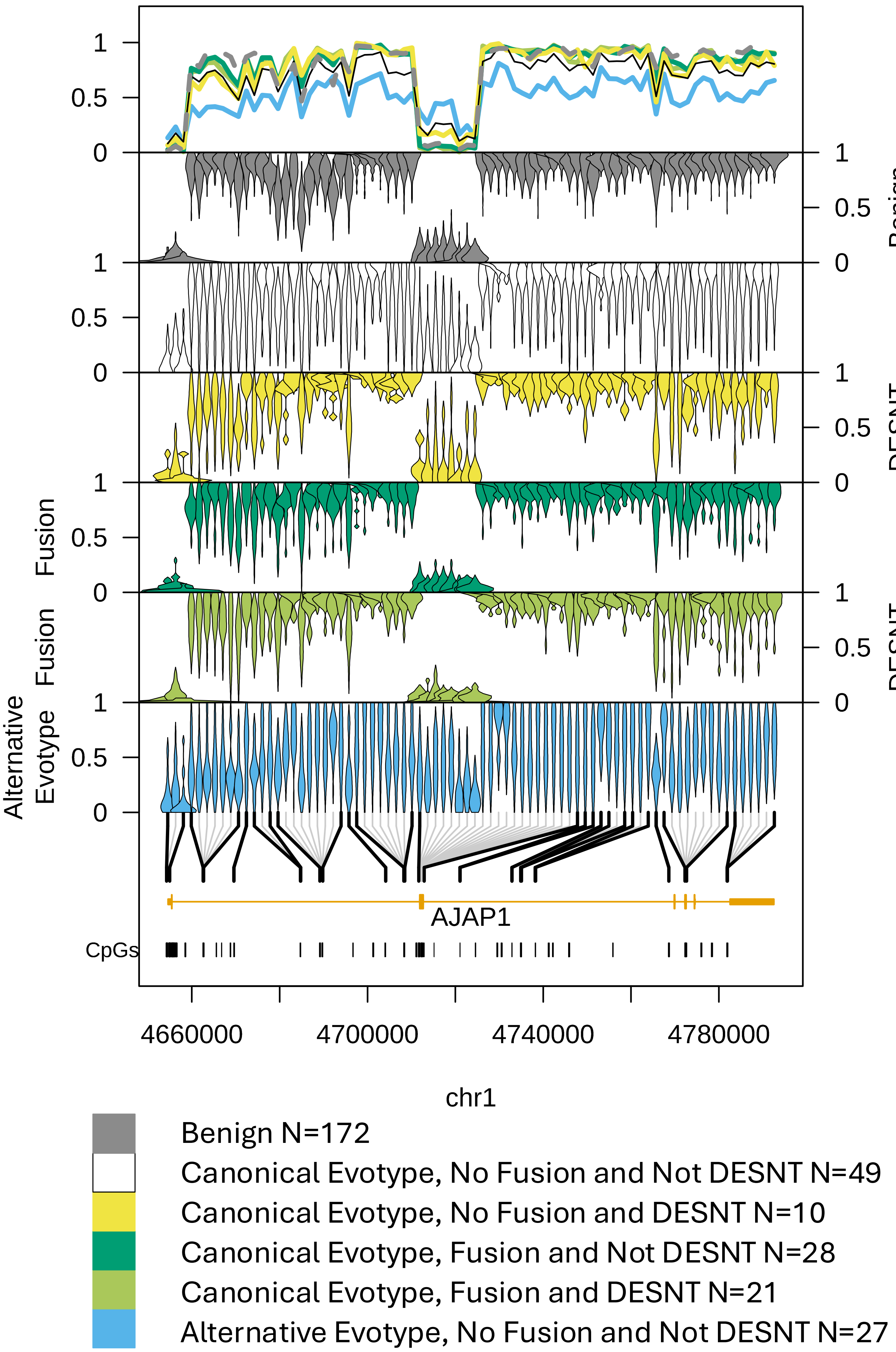

Benign
DESNT
Fusion
Fusion
DESNT
Alternative
Evotype
AJAP1
CpGs
4660000
4700000
4740000
4780000
chr1
Benign N=172
Canonical Evotype, No Fusion and Not DESNT N=49
Canonical Evotype, No Fusion and DESNT N=10
Canonical Evotype, Fusion and Not DESNT N=28
Canonical Evotype, Fusion and DESNT N=21
Alternative Evotype, No Fusion and Not DESNT N=27

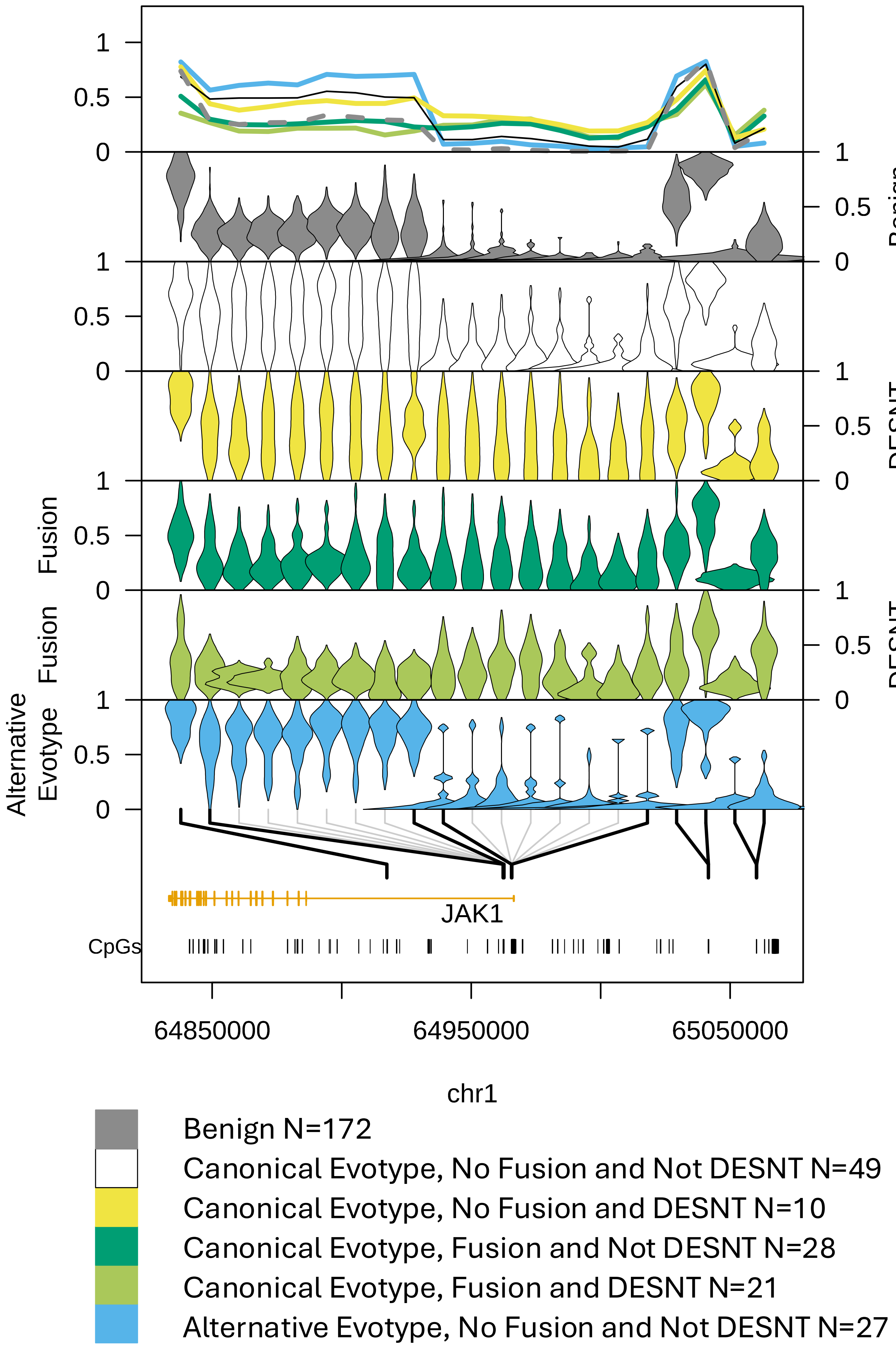

1
0.5
0
Benign
DESNT
Fusion
Fusion DESNT
Alternative Evotype
JAK1
CpGs
64850000
64950000
65050000
chr1
Benign N=172
Canonical Evotype, No Fusion and Not DESNT N=49
Canonical Evotype, No Fusion and DESNT N=10
Canonical Evotype, Fusion and Not DESNT N=28
Canonical Evotype, Fusion and DESNT N=21
Alternative Evotype, No Fusion and Not DESNT N=27

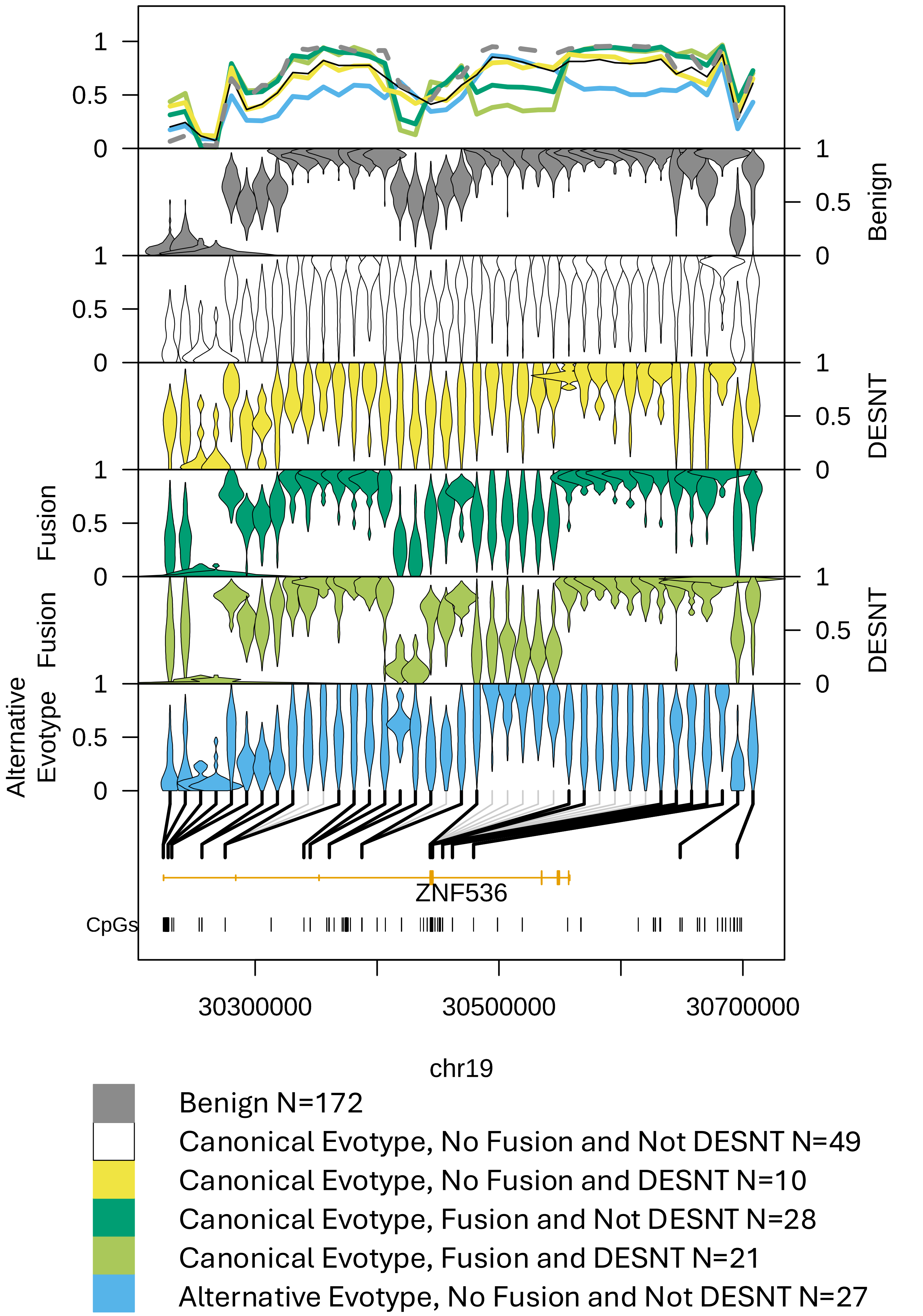

Benign
DESNT
Fusion
Fusion
DESNT
Alternative Evotype
ZNF536
CpGs
30300000
30500000
30700000
chr19
Benign N=172
Canonical Evotype, No Fusion and Not DESNT N=49
Canonical Evotype, No Fusion and DESNT N=10
Canonical Evotype, Fusion and Not DESNT N=28
Canonical Evotype, Fusion and DESNT N=21
Alternative Evotype, No Fusion and Not DESNT N=27

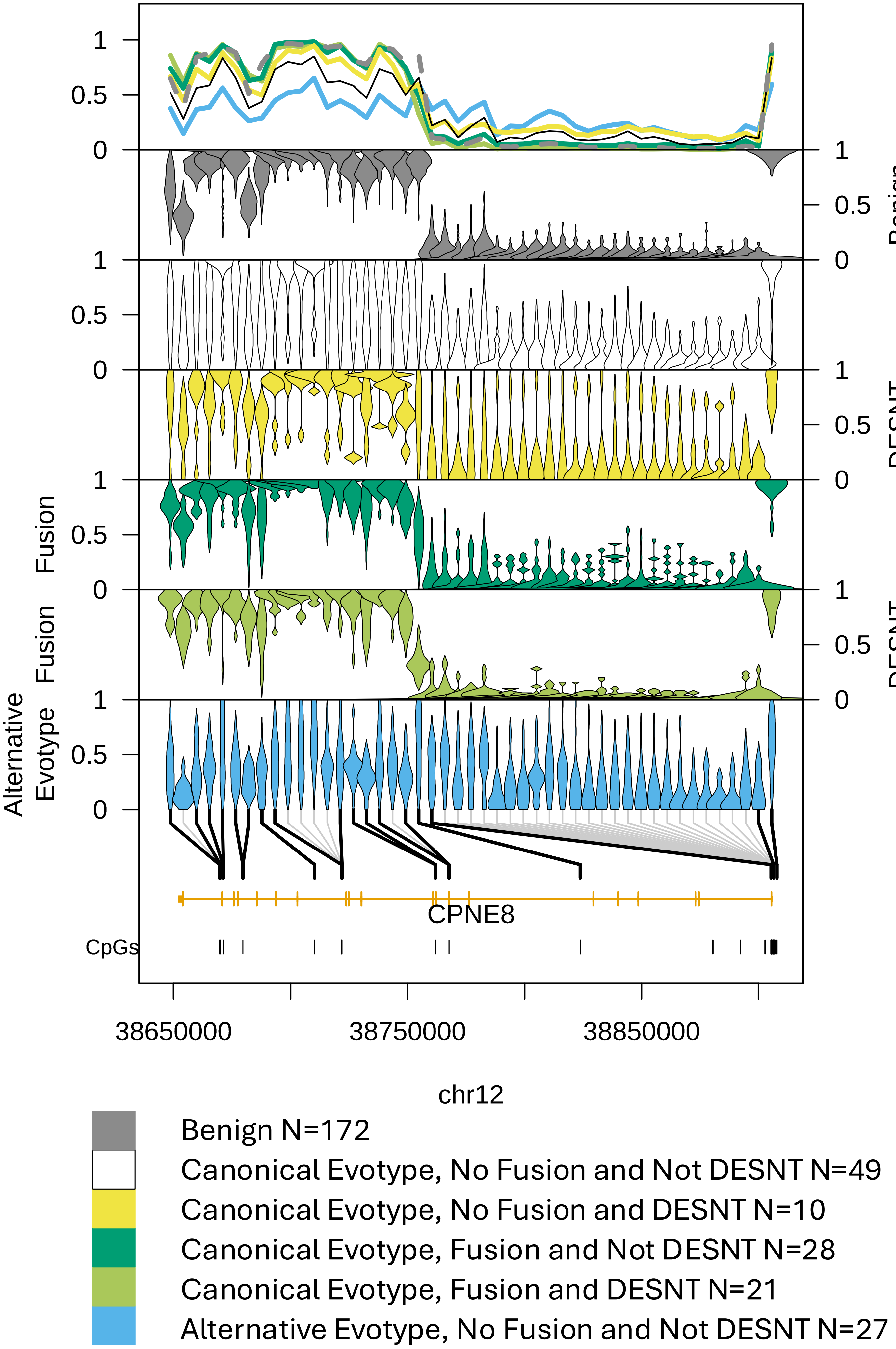

1
0.5
0
Benign
Fusion
DESNT
Alternative Evotype
CPNE8
CpGs
38650000
38750000
38850000
chr12
Benign N=172
Canonical Evotype, No Fusion and Not DESNT N=49
Canonical Evotype, No Fusion and DESNT N=10
Canonical Evotype, Fusion and Not DESNT N=28
Canonical Evotype, Fusion and DESNT N=21
Alternative Evotype, No Fusion and Not DESNT N=27

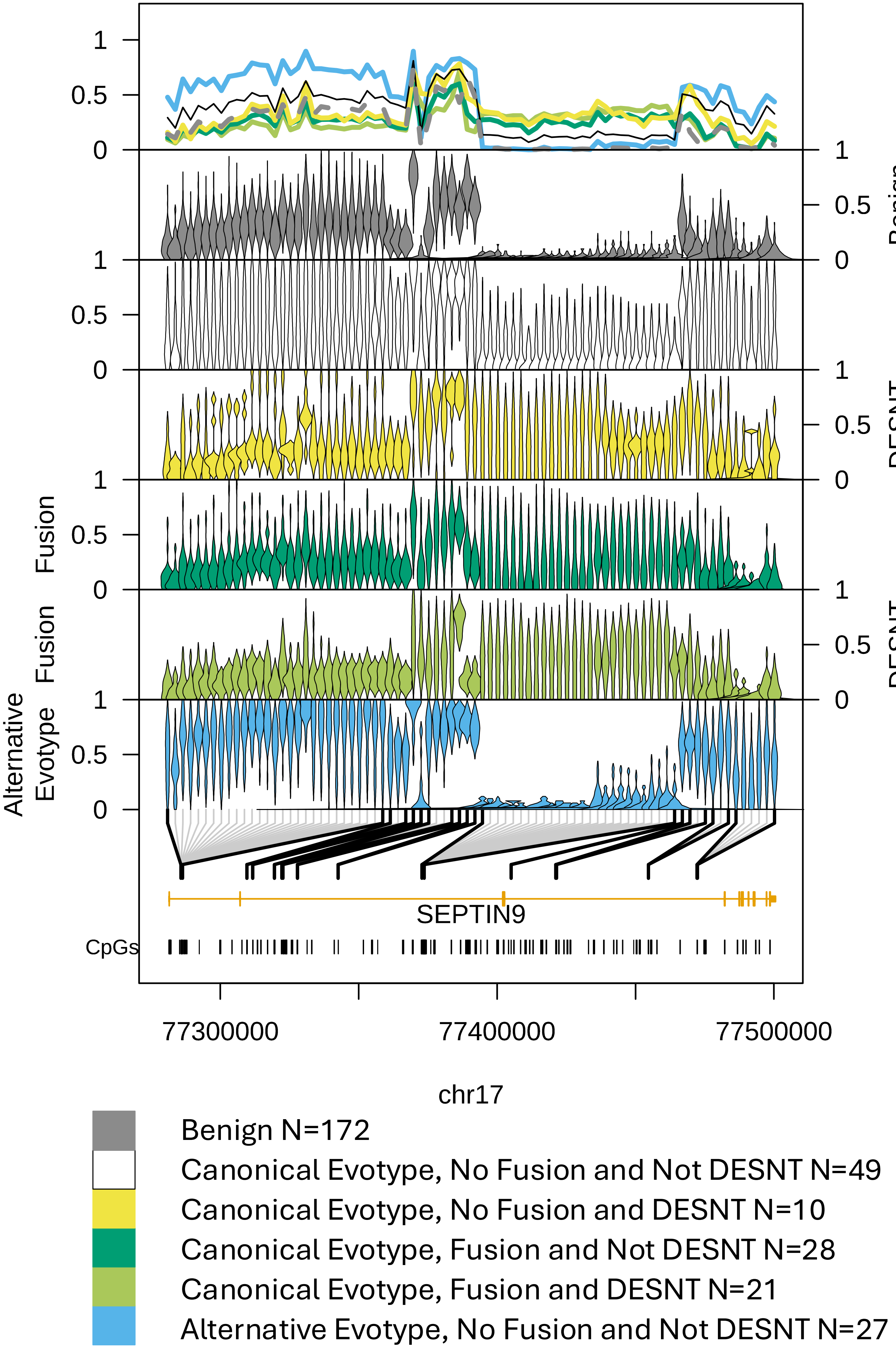

1
0.5
0
1
0.5
0
Benign
1
0.5
0
1
0.5
0
DESNT
Fusion
0.5
0
1
Fusion
1
0.5
0
DESNT
Alternative Evotype
1
0.5
0
SEPTIN9
CpGs
77300000
77400000
77500000
chr17
Benign N=172
Canonical Evotype, No Fusion and Not DESNT N=49
Canonical Evotype, No Fusion and DESNT N=10
Canonical Evotype, Fusion and Not DESNT N=28
Canonical Evotype, Fusion and DESNT N=21
Alternative Evotype, No Fusion and Not DESNT N=27

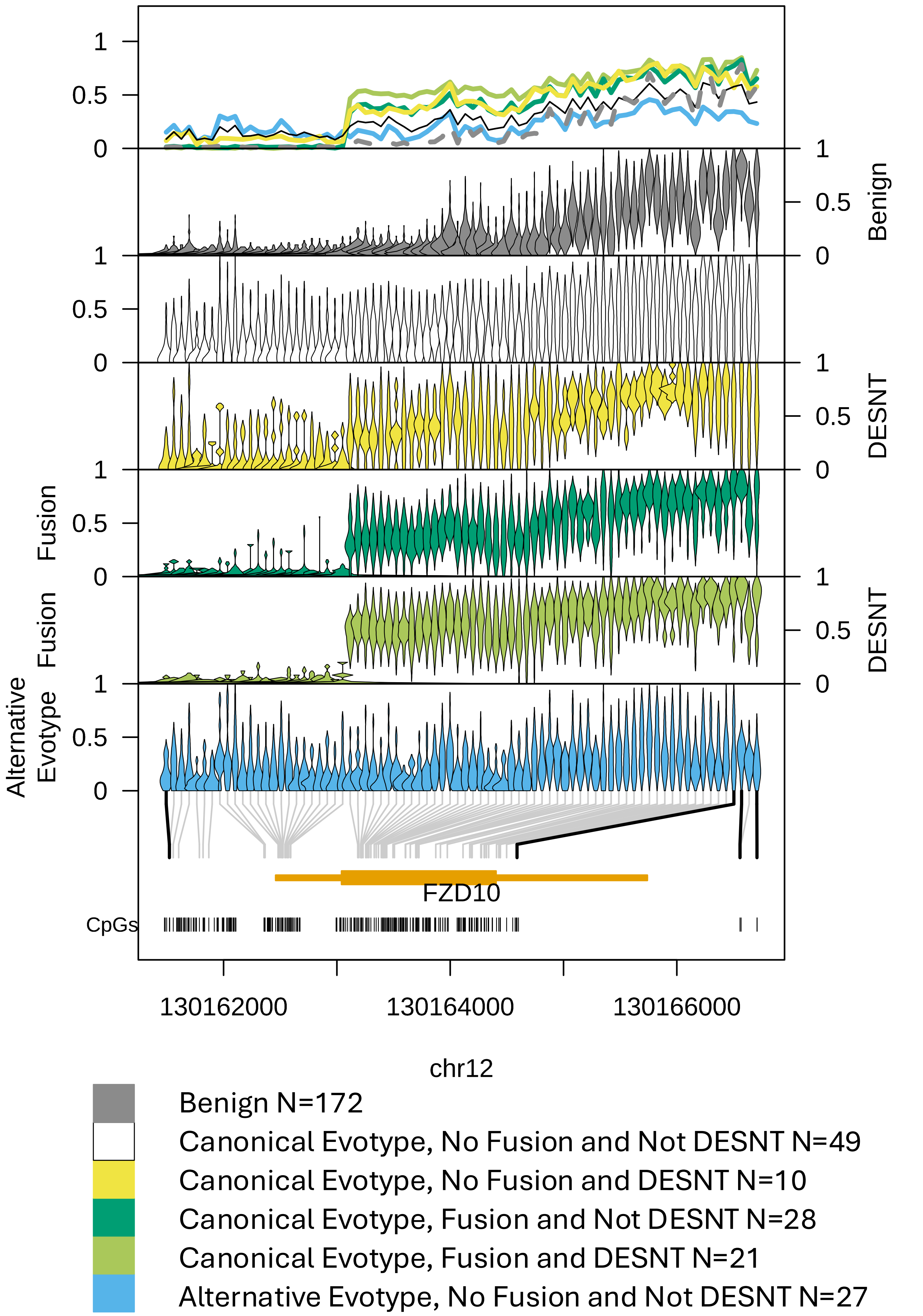

Benign
DESNT
Fusion
Fusion
DESNT
Alternative
Evotype
FZD10
CpGs
130162000
130164000
130166000
chr12
Benign N=172
Canonical Evotype, No Fusion and Not DESNT N=49
Canonical Evotype, No Fusion and DESNT N=10
Canonical Evotype, Fusion and Not DESNT N=28
Canonical Evotype, Fusion and DESNT N=21
Alternative Evotype, No Fusion and Not DESNT N=27

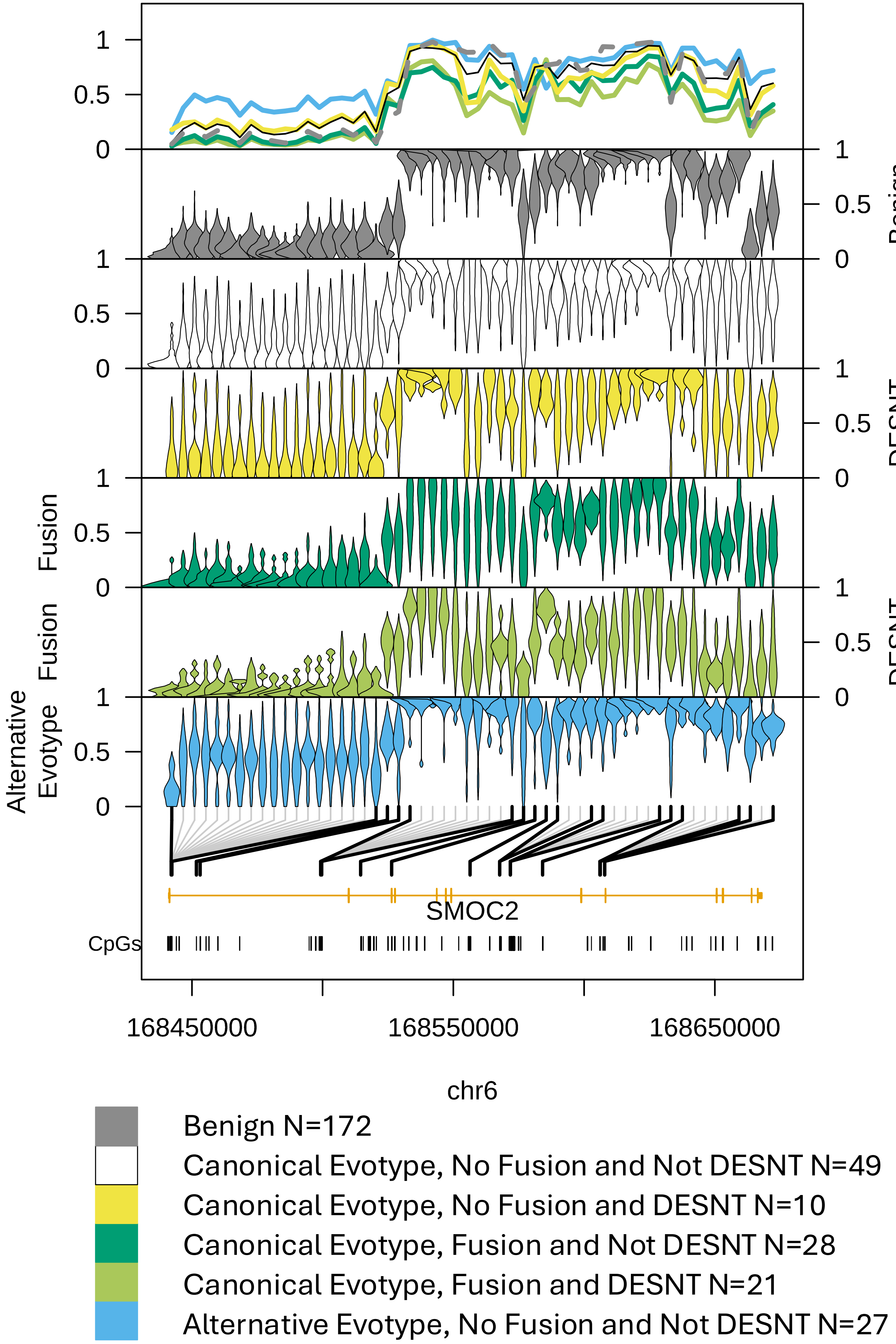

1
0.5
0
Benign
DESNT
Fusion
Fusion
DESNT
Alternative Evotype
SMOC2
CpGs
168450000
168550000
168650000
chr6
Benign N=172
Canonical Evotype, No Fusion and Not DESNT N=49
Canonical Evotype, No Fusion and DESNT N=10
Canonical Evotype, Fusion and Not DESNT N=28
Canonical Evotype, Fusion and DESNT N=21
Alternative Evotype, No Fusion and Not DESNT N=27

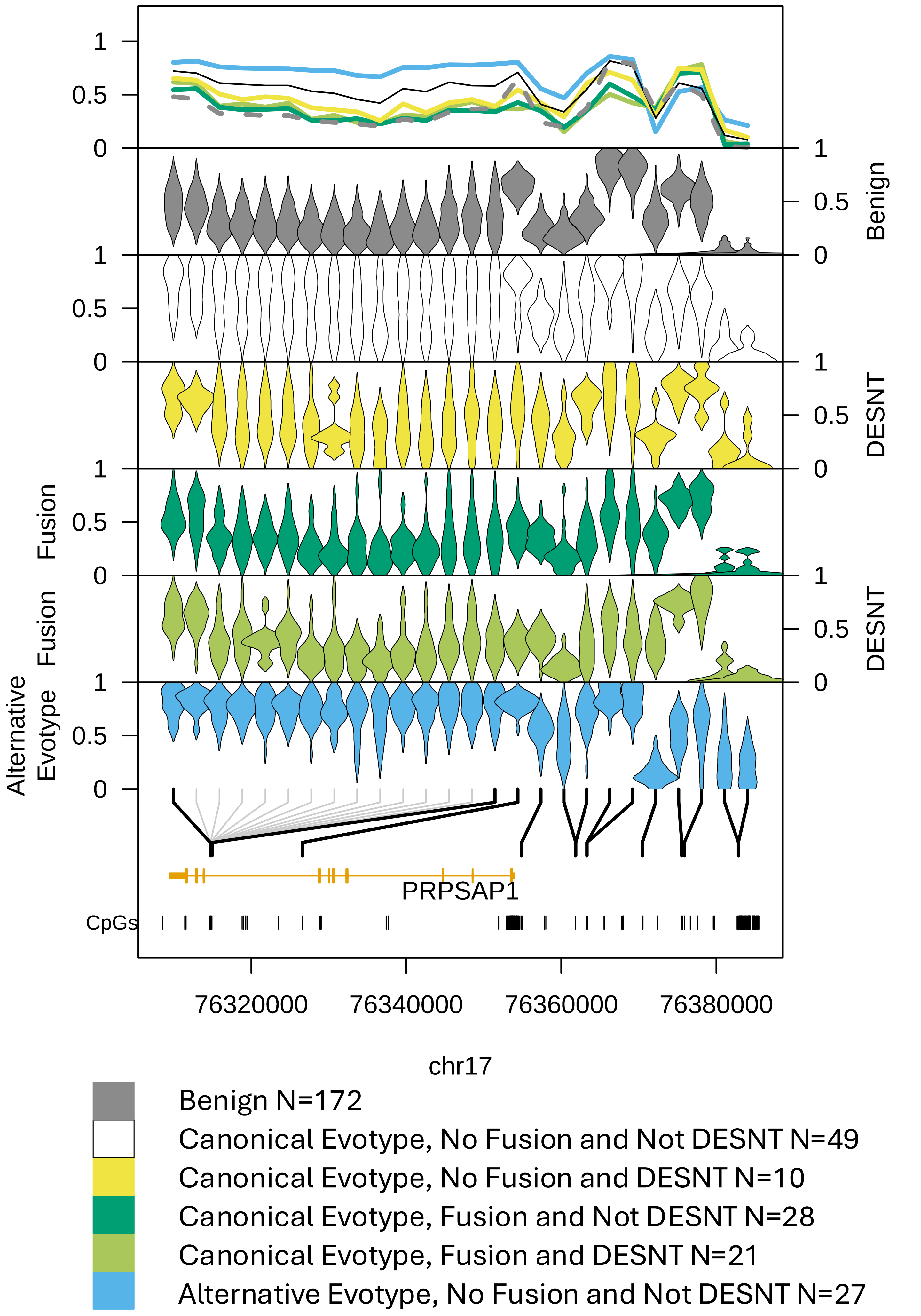

Benign
DESNT
Fusion
Fusion DESNT
Alternative Evotype
PRPSAP1
CpGs
76320000
76340000
76360000
76380000
chr17
Benign N=172
Canonical Evotype, No Fusion and Not DESNT N=49
Canonical Evotype, No Fusion and DESNT N=10
Canonical Evotype, Fusion and Not DESNT N=28
Canonical Evotype, Fusion and DESNT N=21
Alternative Evotype, No Fusion and Not DESNT N=27

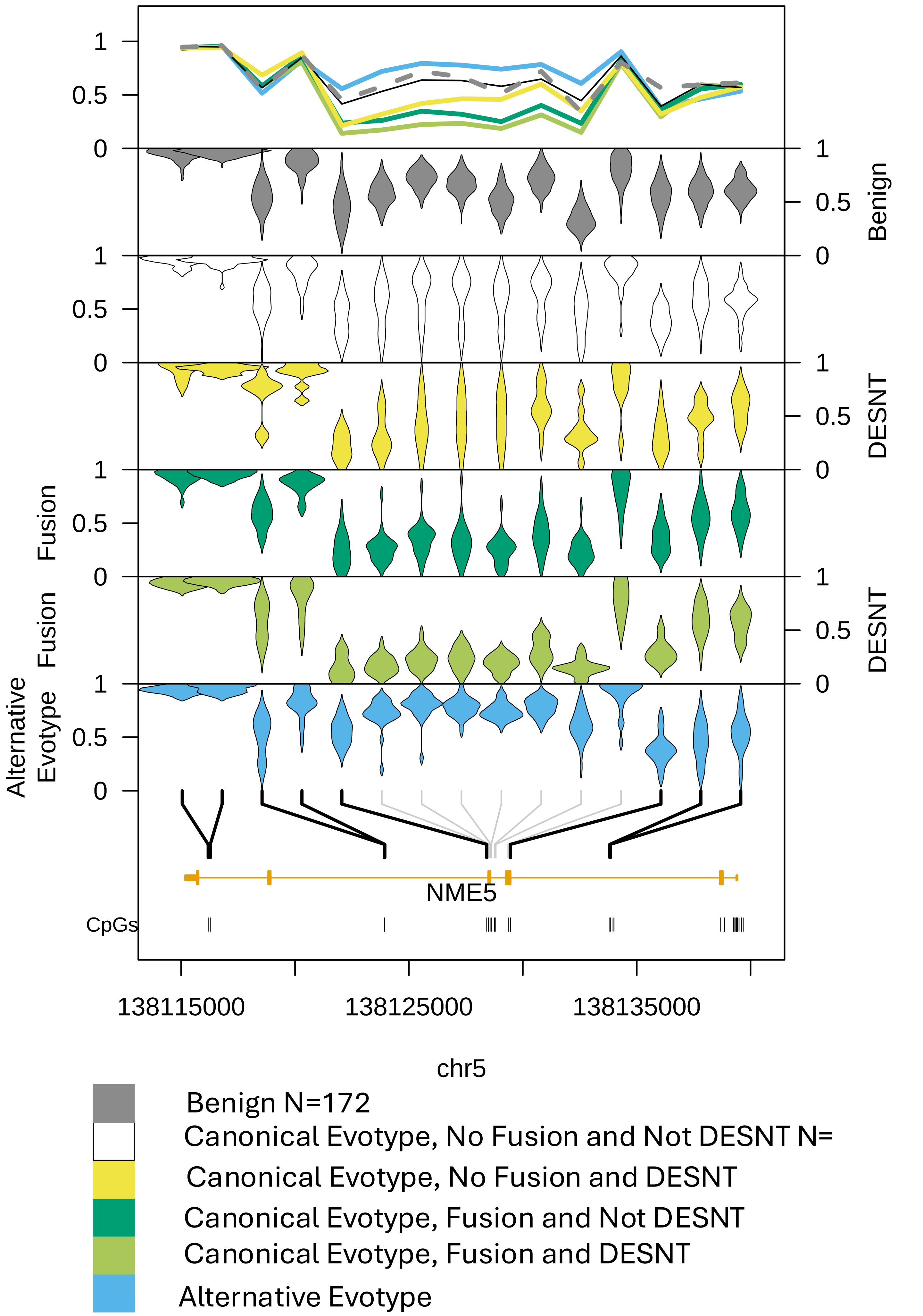

Benign
DESNT
Fusion
Fusion
DESNT
Alternative Evotype
NME5
CpGs
138115000
138125000
138135000
chr5
Benign N=172
Canonical Evotype, No Fusion and Not DESNT N=
Canonical Evotype, No Fusion and DESNT
Canonical Evotype, Fusion and Not DESNT
Canonical Evotype, Fusion and DESNT
Alternative Evotype

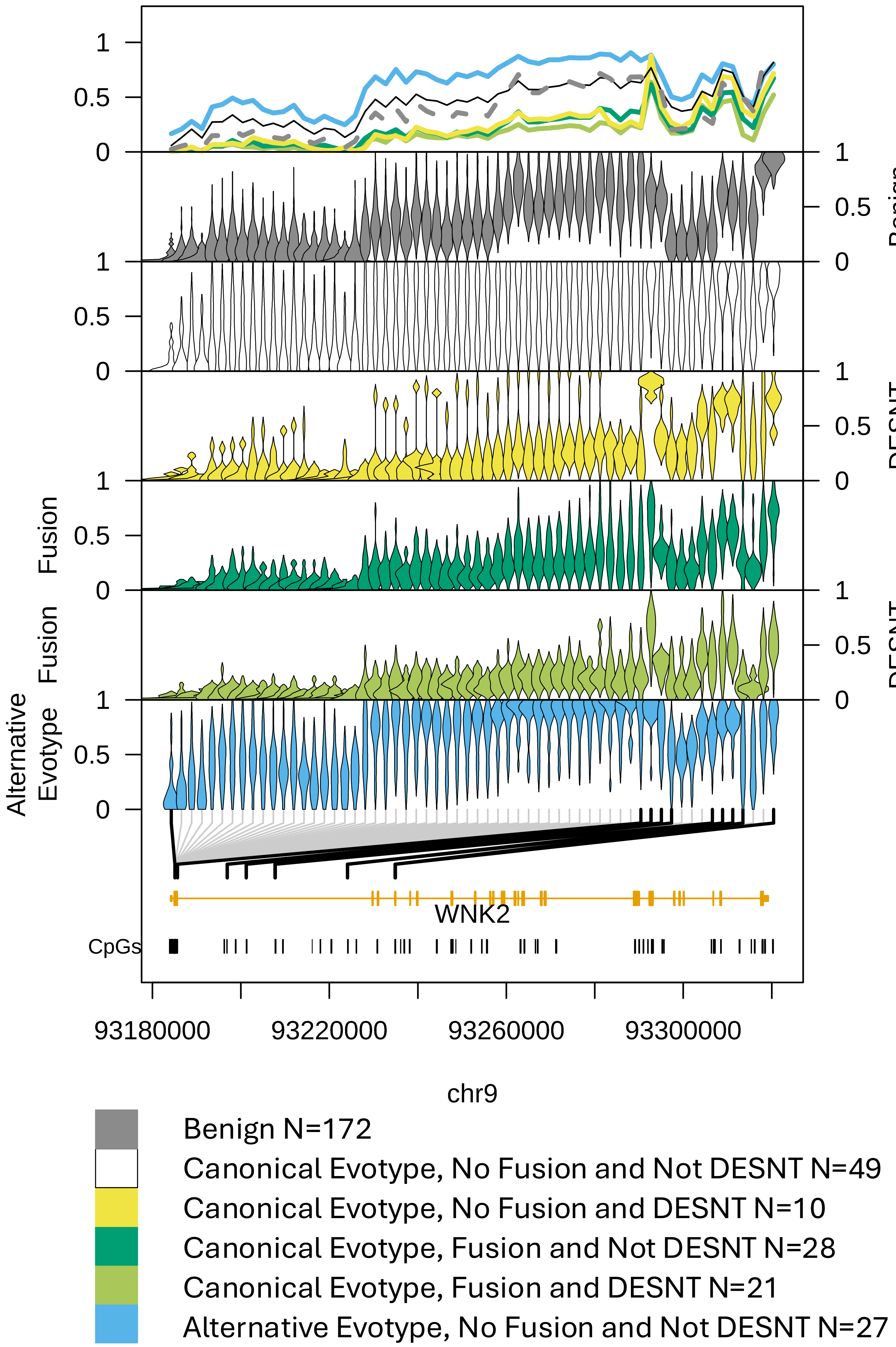

Benign
DESNT
Fusion
Fusion
DESNT
Alternative
Evotype
WNK2
CpGs
93180000
93220000
93260000
93300000
chr9
Benign N=172
Canonical Evotype, No Fusion and Not DESNT N=49
Canonical Evotype, No Fusion and DESNT N=10
Canonical Evotype, Fusion and Not DESNT N=28
Canonical Evotype, Fusion and DESNT N=21
Alternative Evotype, No Fusion and Not DESNT N=27

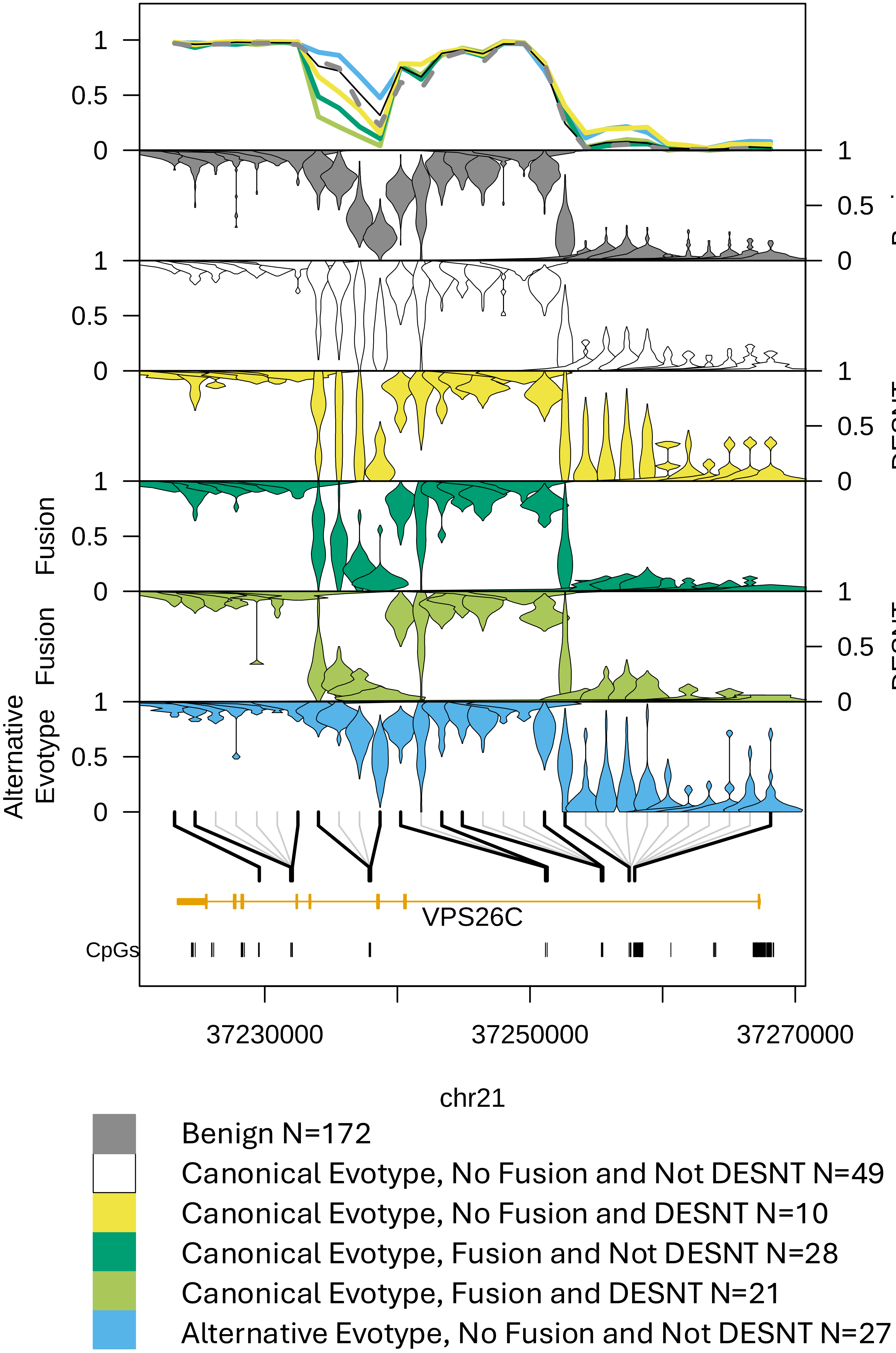

Benign
DESNT
Fusion
Fusion DESNT
Alternative Evotype
VPS26C
CpGs
37230000
37250000
37270000
chr21
Benign N=172
Canonical Evotype, No Fusion and Not DESNT N=49
Canonical Evotype, No Fusion and DESNT N=10
Canonical Evotype, Fusion and Not DESNT N=28
Canonical Evotype, Fusion and DESNT N=21
Alternative Evotype, No Fusion and Not DESNT N=27

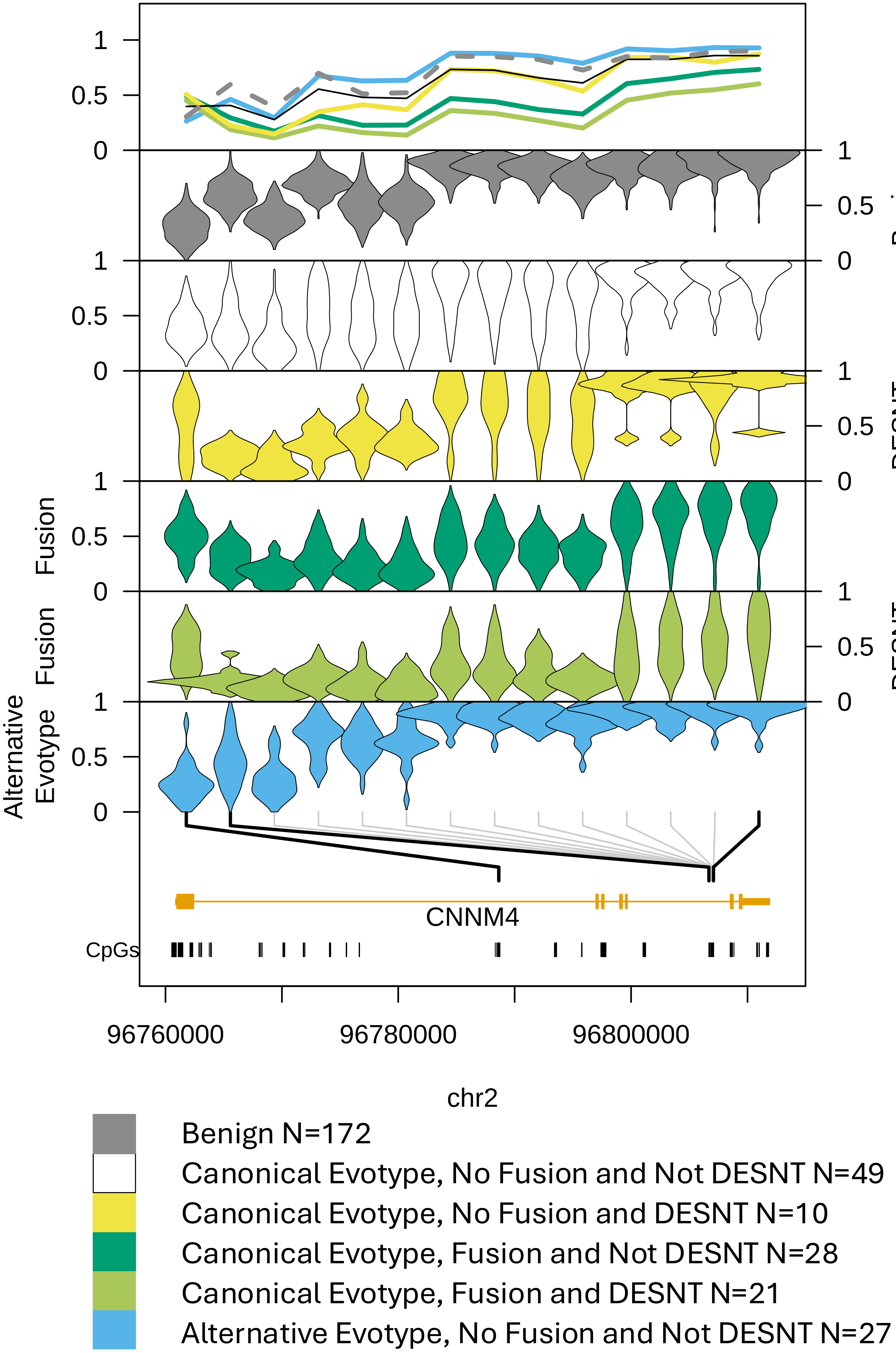

1
0.5
0
Benign
DESNT
Fusion
Fusion
DESNT
Alternative
Evotype
CNNM4
CpGs
96760000
96780000
96800000
chr2
Benign N=172
Canonical Evotype, No Fusion and Not DESNT N=49
Canonical Evotype, No Fusion and DESNT N=10
Canonical Evotype, Fusion and Not DESNT N=28
Canonical Evotype, Fusion and DESNT N=21
Alternative Evotype, No Fusion and Not DESNT N=27

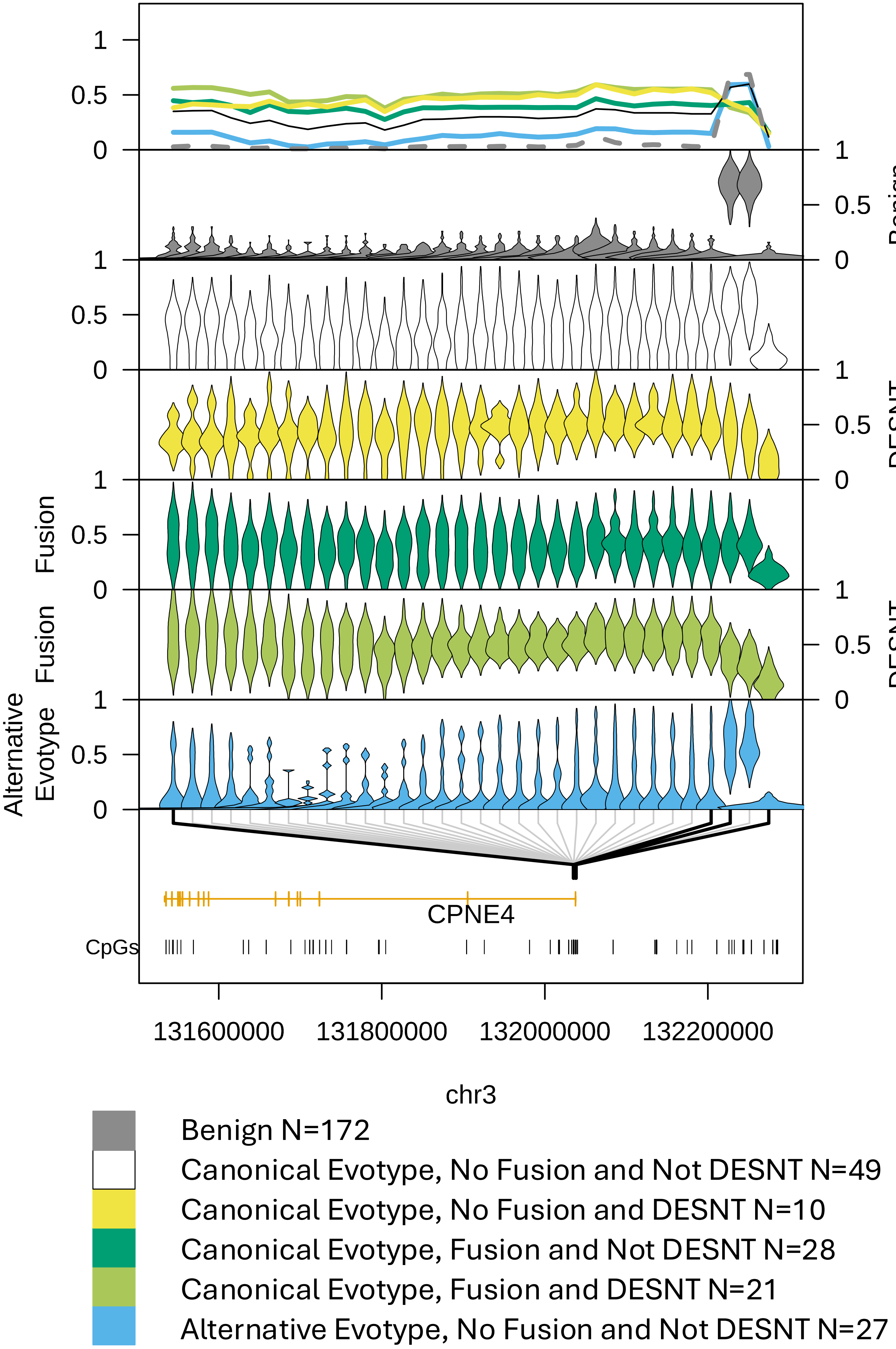

1
0.5
0
Benign
DESNT
Fusion
Fusion
DESNT
Alternative
Evotype
CPNE4
CpGs
131600000
131800000
132000000
132200000
chr3
Benign N=172
Canonical Evotype, No Fusion and Not DESNT N=49
Canonical Evotype, No Fusion and DESNT N=10
Canonical Evotype, Fusion and Not DESNT N=28
Canonical Evotype, Fusion and DESNT N=21
Alternative Evotype, No Fusion and Not DESNT N=27

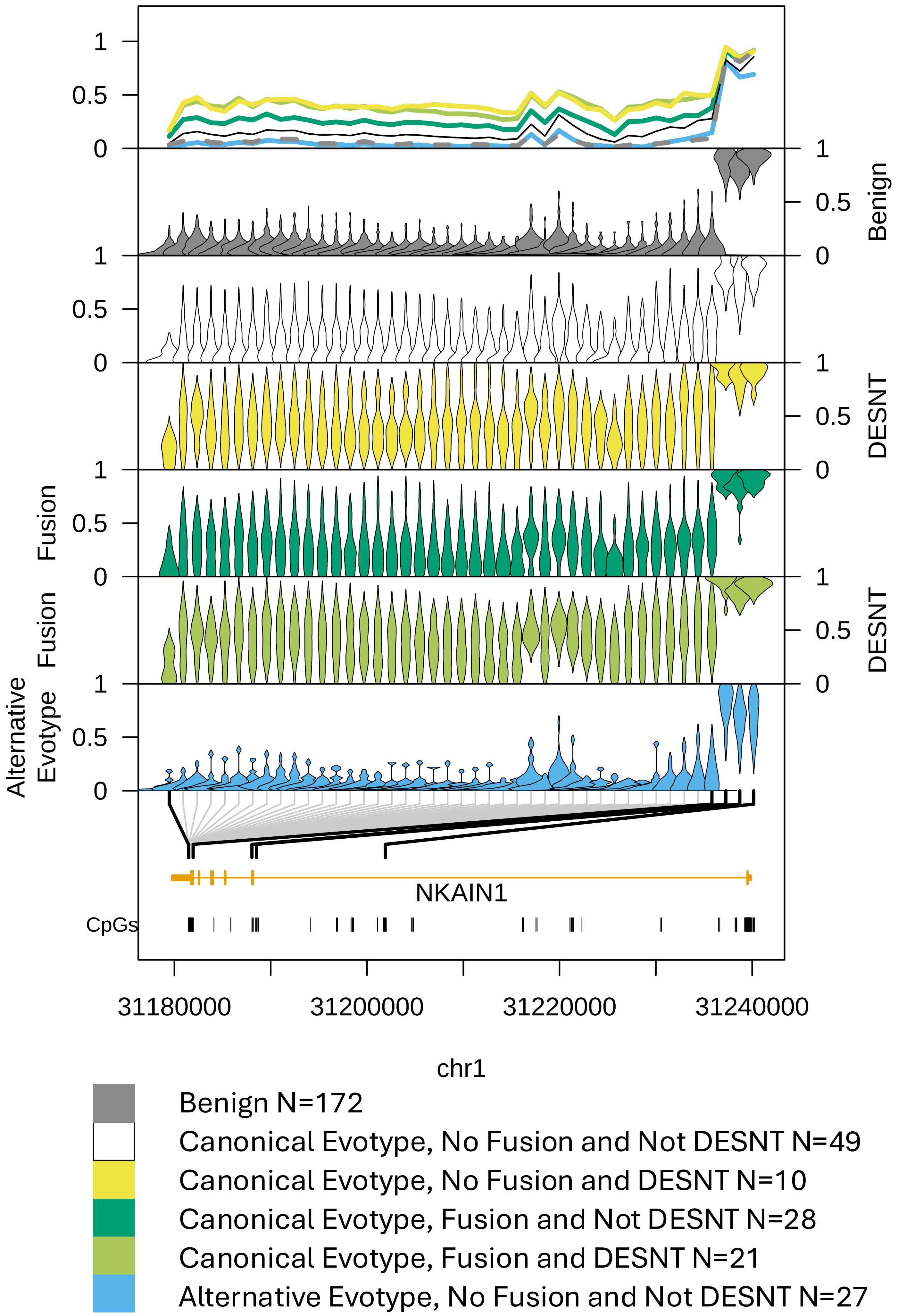

Benign
DESNT
Fusion
Fusion
DESNT
Alternative
Evotype
NKAIN1
CpGs
31180000
31200000
31220000
31240000
chr1
Benign N=172
Canonical Evotype, No Fusion and Not DESNT N=49
Canonical Evotype, No Fusion and DESNT N=10
Canonical Evotype, Fusion and Not DESNT N=28
Canonical Evotype, Fusion and DESNT N=21
Alternative Evotype, No Fusion and Not DESNT N=27

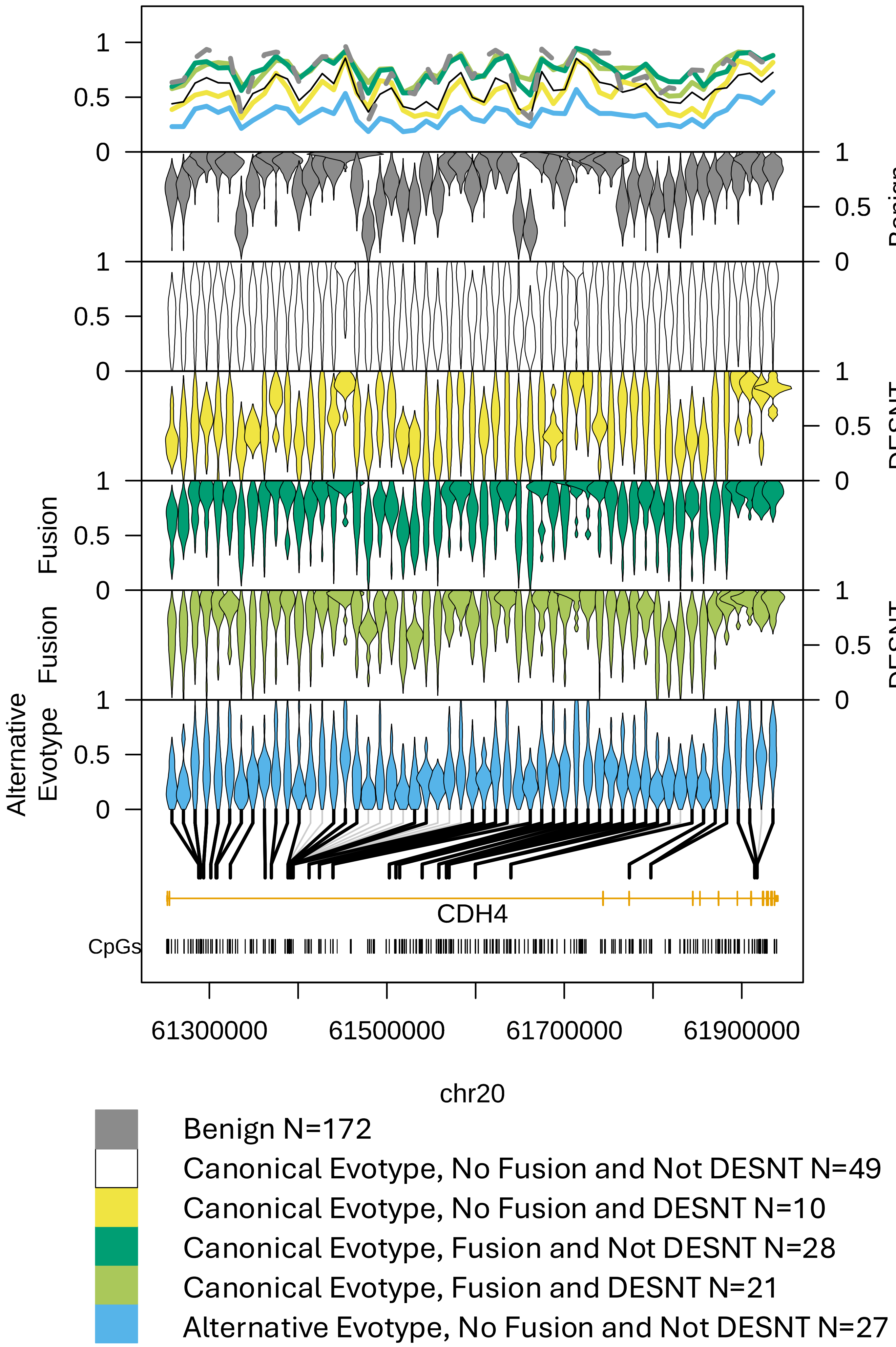

1
0.5
0
Benign
DESNT
Fusion
Alternative Evotype
CDH4
CpGs
61300000
61500000
61700000
61900000
chr20
Benign N=172
Canonical Evotype, No Fusion and Not DESNT N=49
Canonical Evotype, No Fusion and DESNT N=10
Canonical Evotype, Fusion and Not DESNT N=28
Canonical Evotype, Fusion and DESNT N=21
Alternative Evotype, No Fusion and Not DESNT N=27

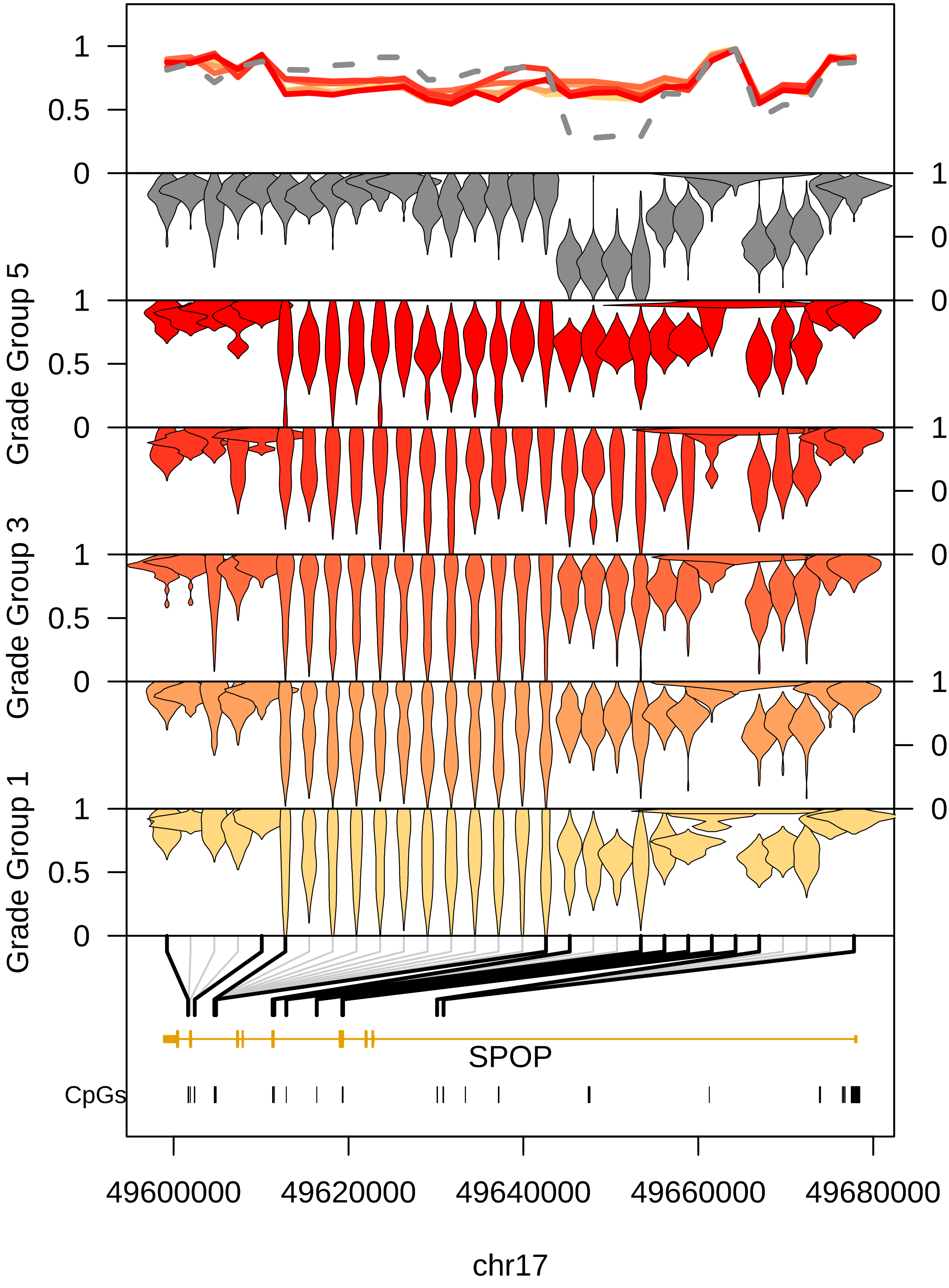

Grade Group 5
Grade Group 3
Grade Group 1
Benign
Grade Group 4
Grade Group 2
SPOP
CpGs
49600000
49620000
49640000
49660000
49680000
chr17

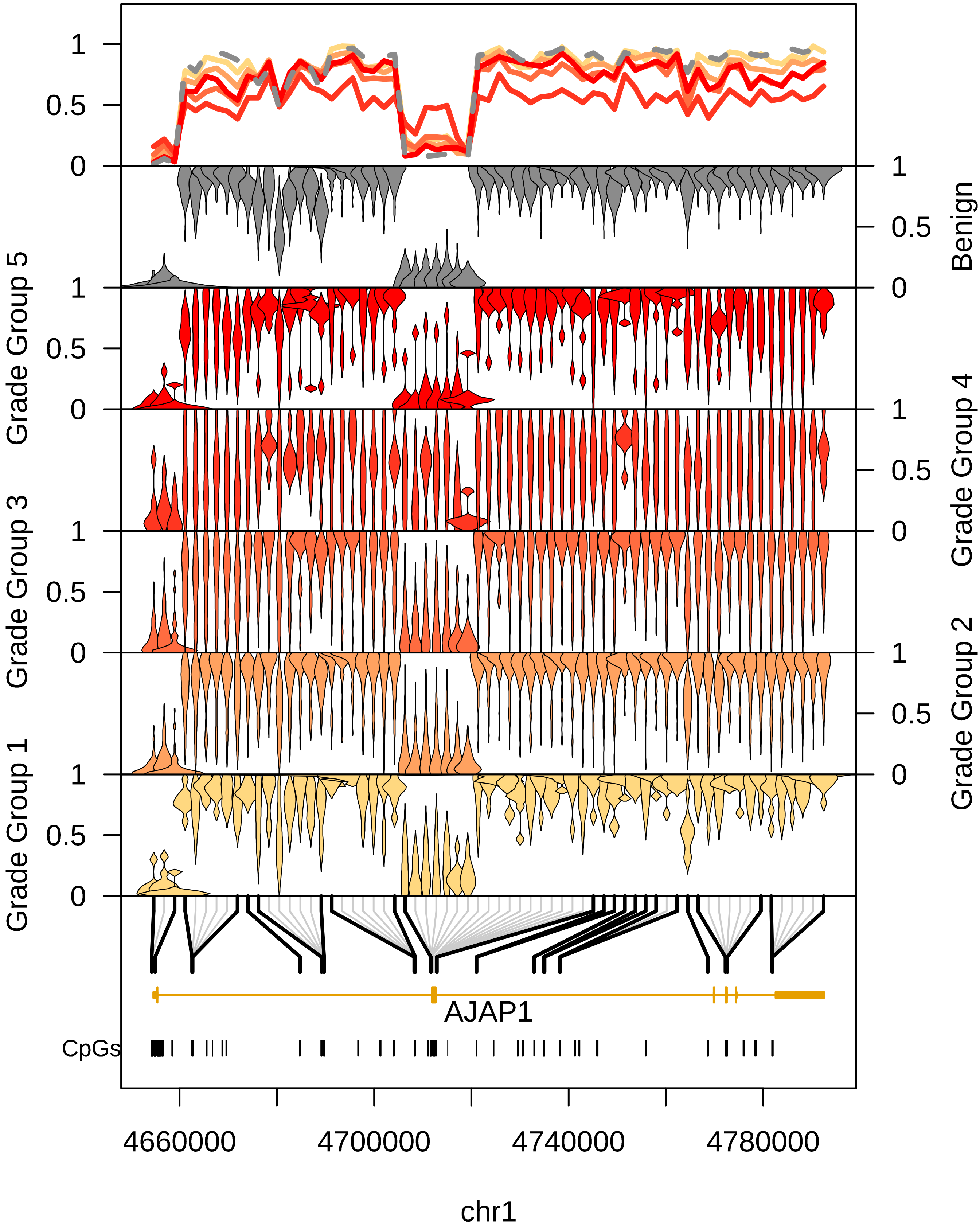

Benign
Grade Group 5
Grade Group 4
Grade Group 3
Grade Group 2
Grade Group 1
AJAP1
CpGs
4660000
4700000
4740000
4780000
chr1

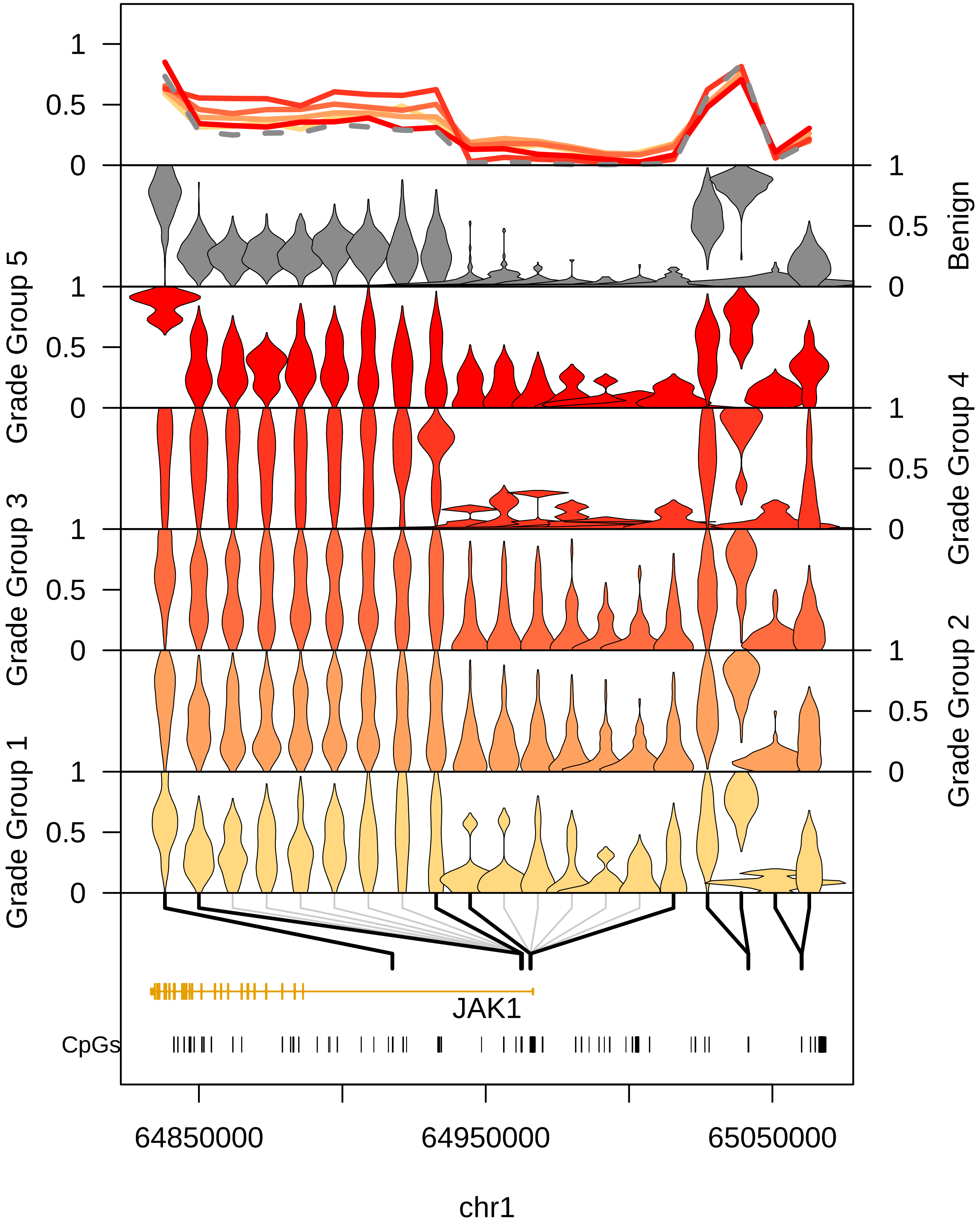

1
0.5
0
Benign
Grade Group 5
Grade Group 4
Grade Group 3
Grade Group 2
Grade Group 1
JAK1
CpGs
64850000
64950000
65050000
chr1

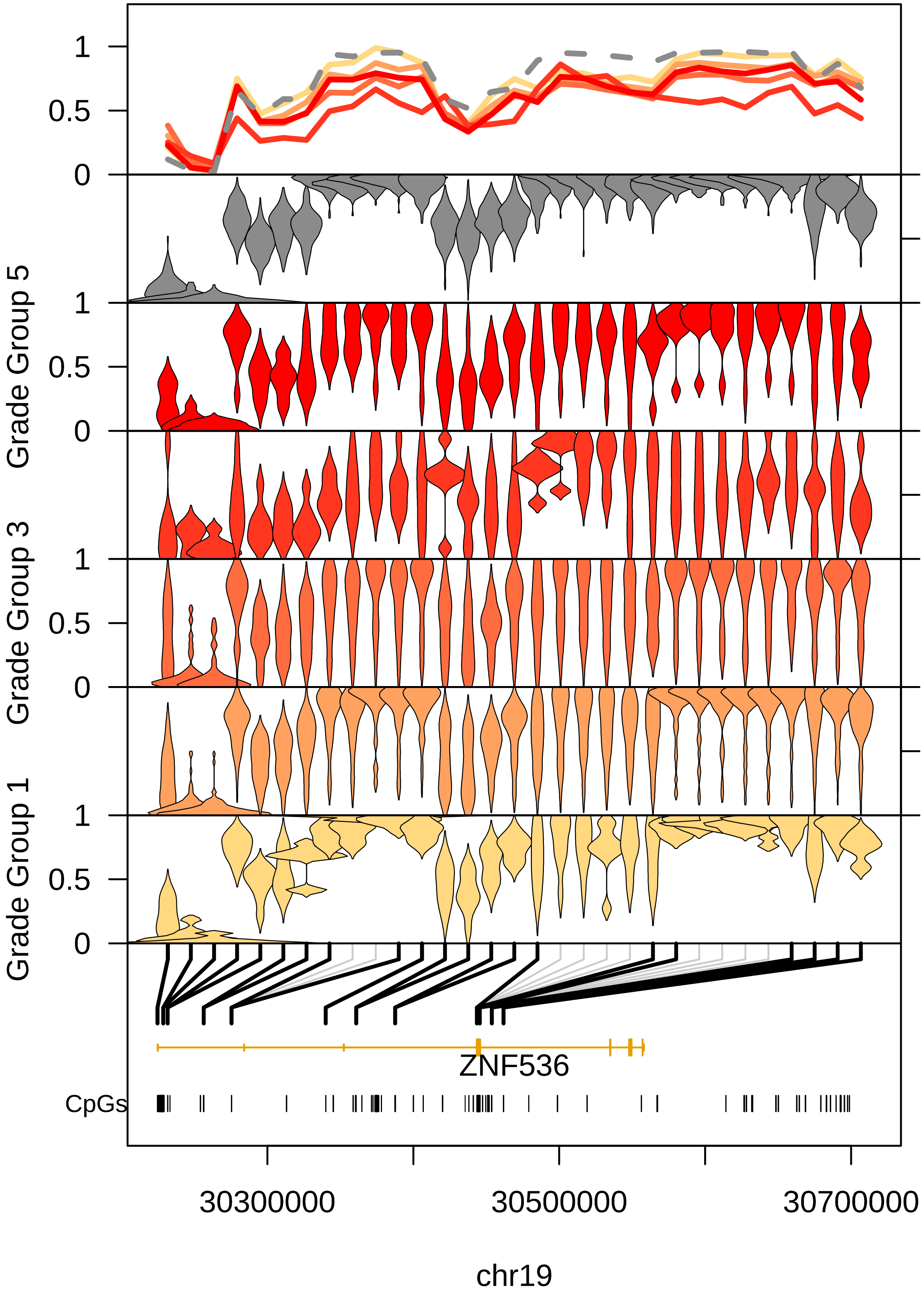

1
0.5
0
Benign
Grade Group 5
Grade Group 4
Grade Group 3
Grade Group 2
Grade Group 1
ZNF536
CpGs
30300000
30500000
30700000
chr19

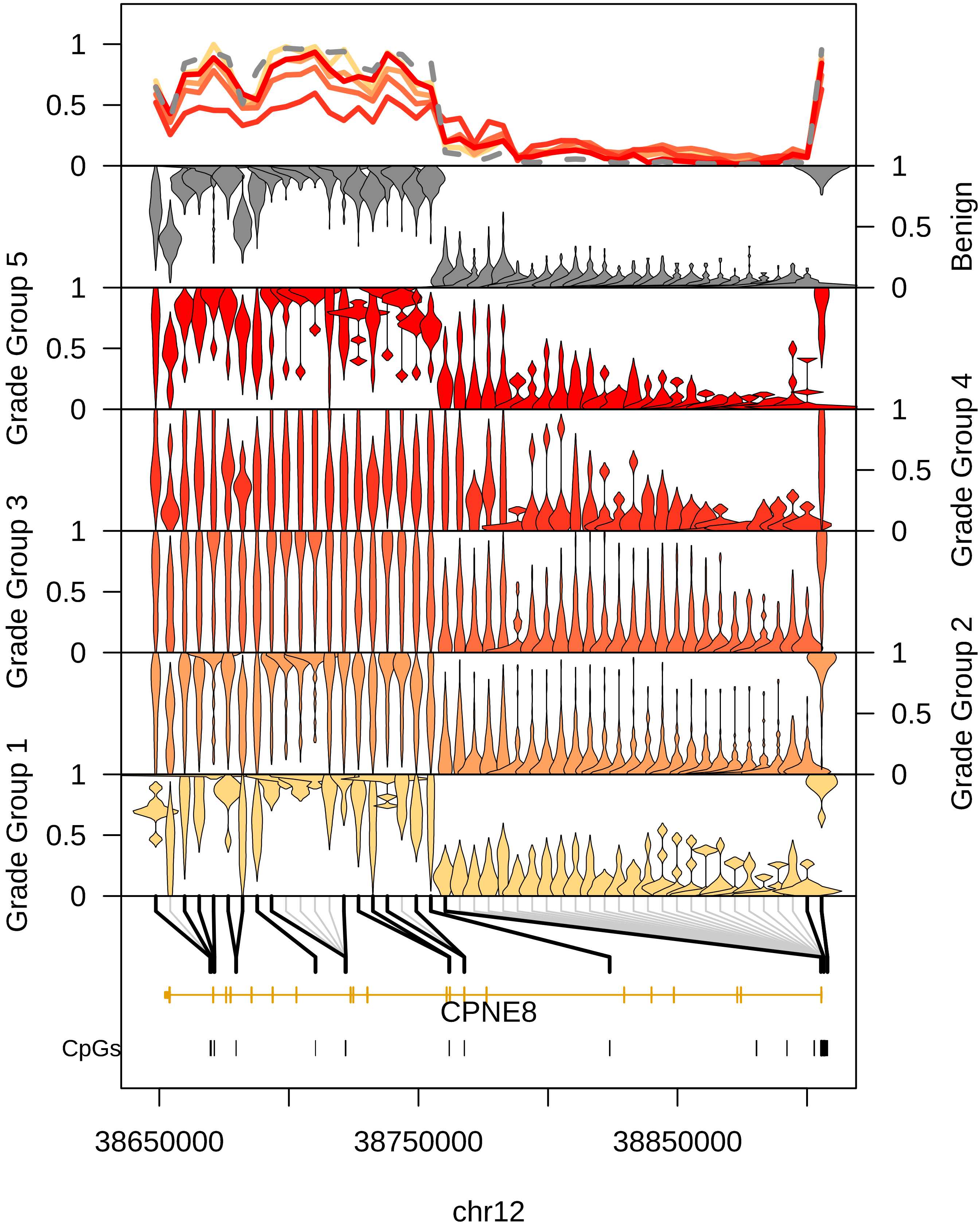

1
0.5
0
Benign
Grade Group 5
Grade Group 4
Grade Group 3
Grade Group 2
Grade Group 1
CPNE8
CpGs
38650000
38750000
38850000
chr12

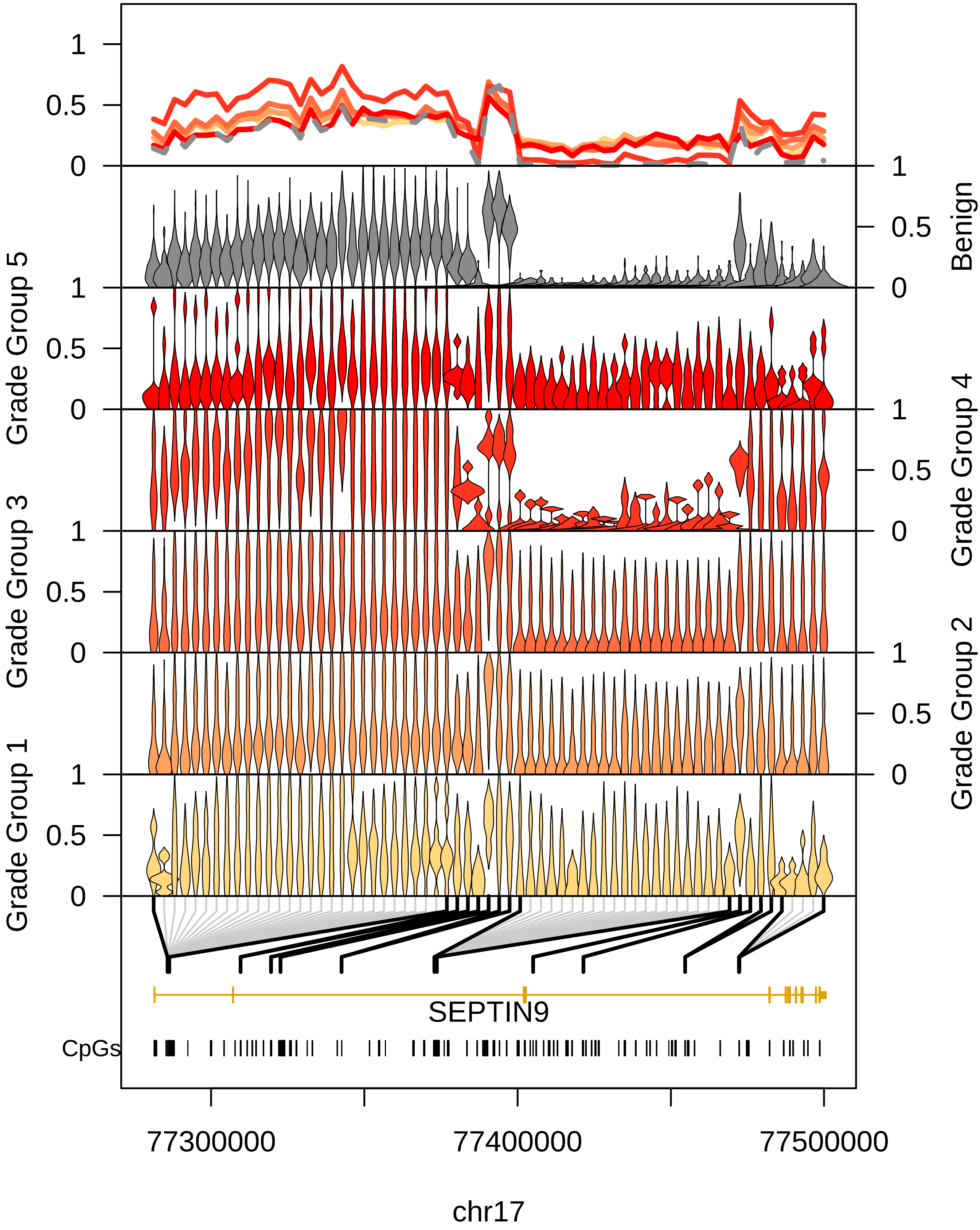

Benign
Grade Group 5
Grade Group 4
Grade Group 3
Grade Group 2
Grade Group 1
1
0.5
0
SEPTIN9
CpGs
77300000
77400000
77500000
chr17

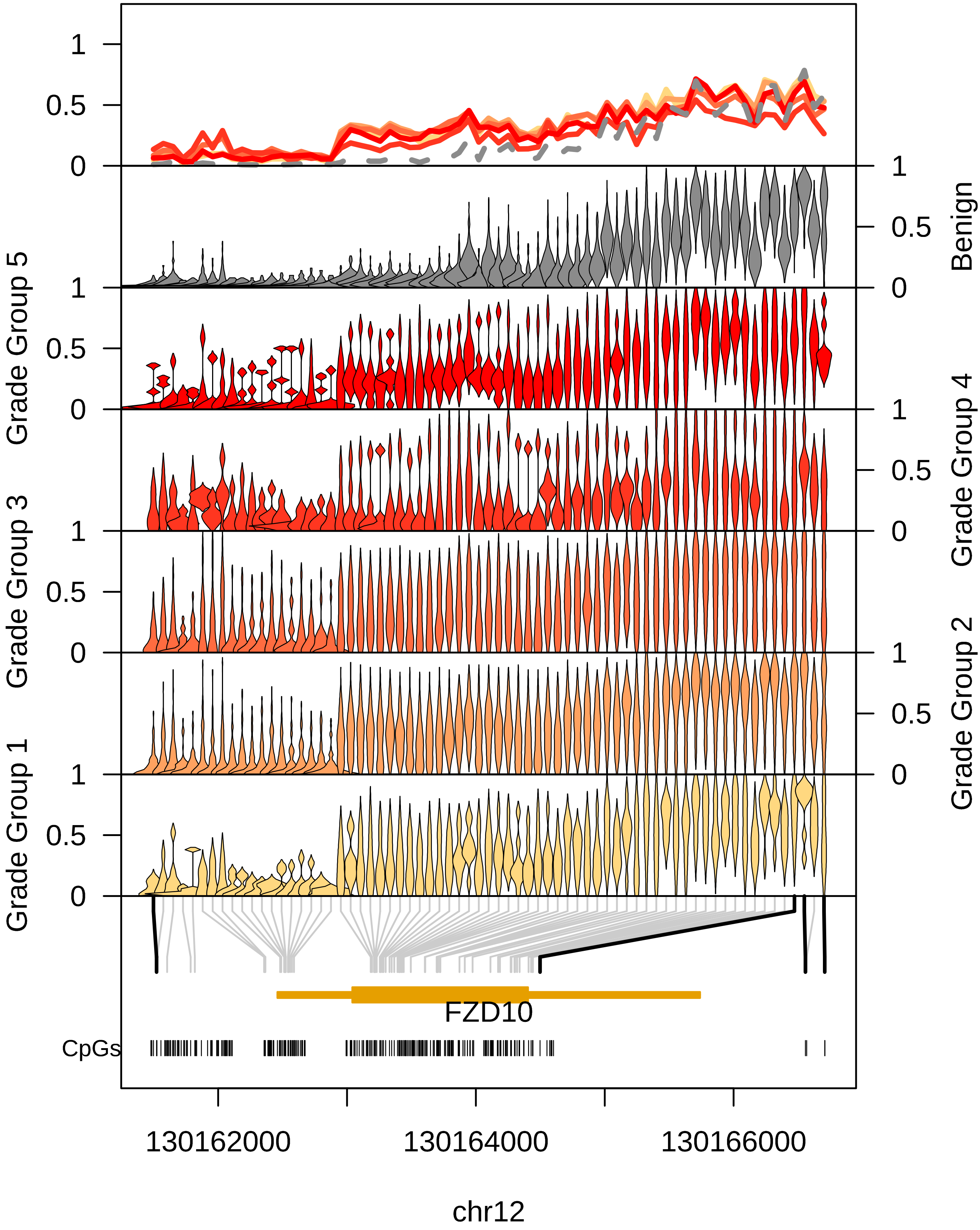

1
0.5
0
Benign
Grade Group 5
Grade Group 4
Grade Group 3
Grade Group 2
Grade Group 1
FZD10
CpGs
130162000
130164000
130166000
chr12

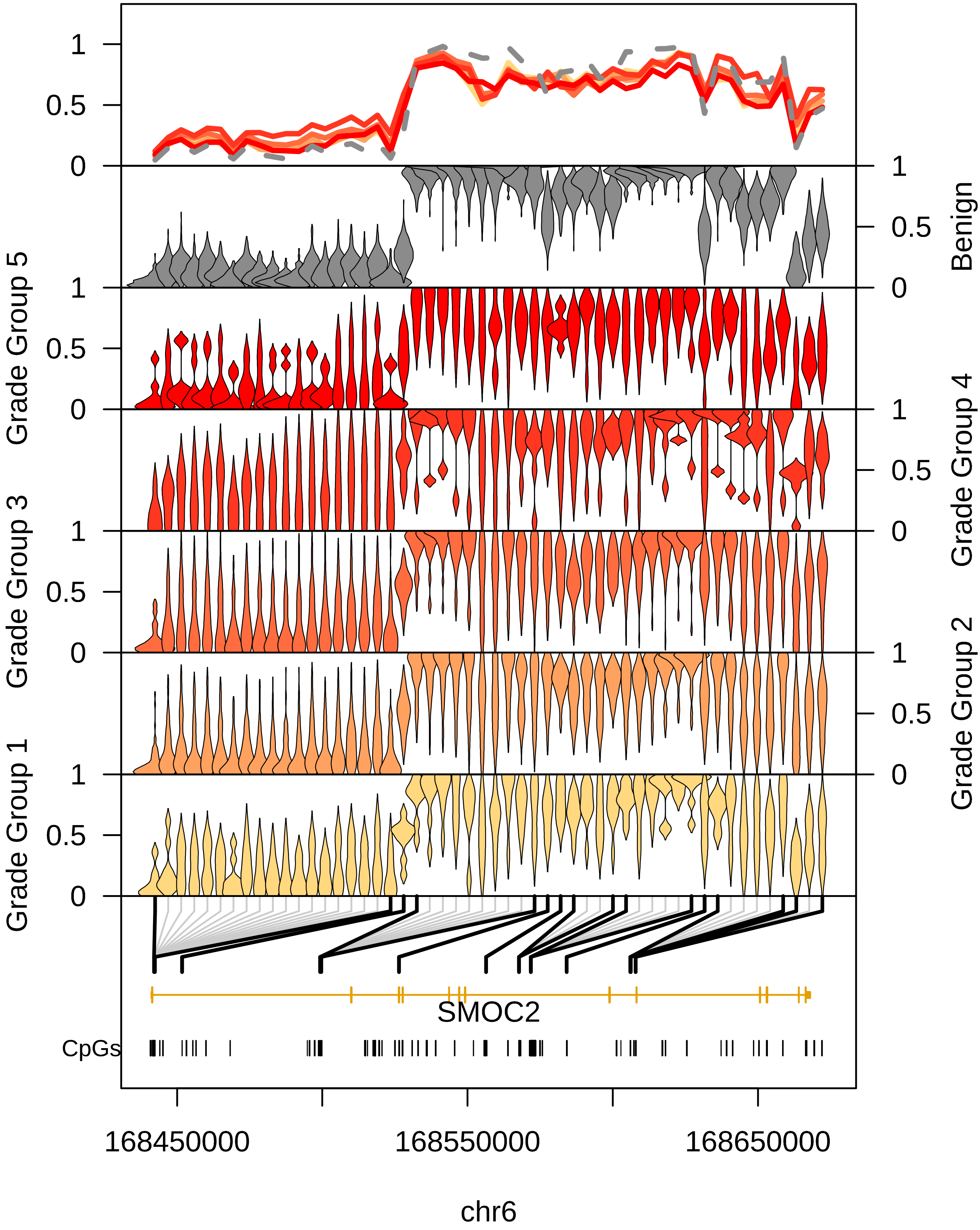

1
0.5
0
Benign
Grade Group 5
Grade Group 4
Grade Group 3
Grade Group 2
Grade Group 1
SMOC2
CpGs
168450000
168550000
168650000
chr6

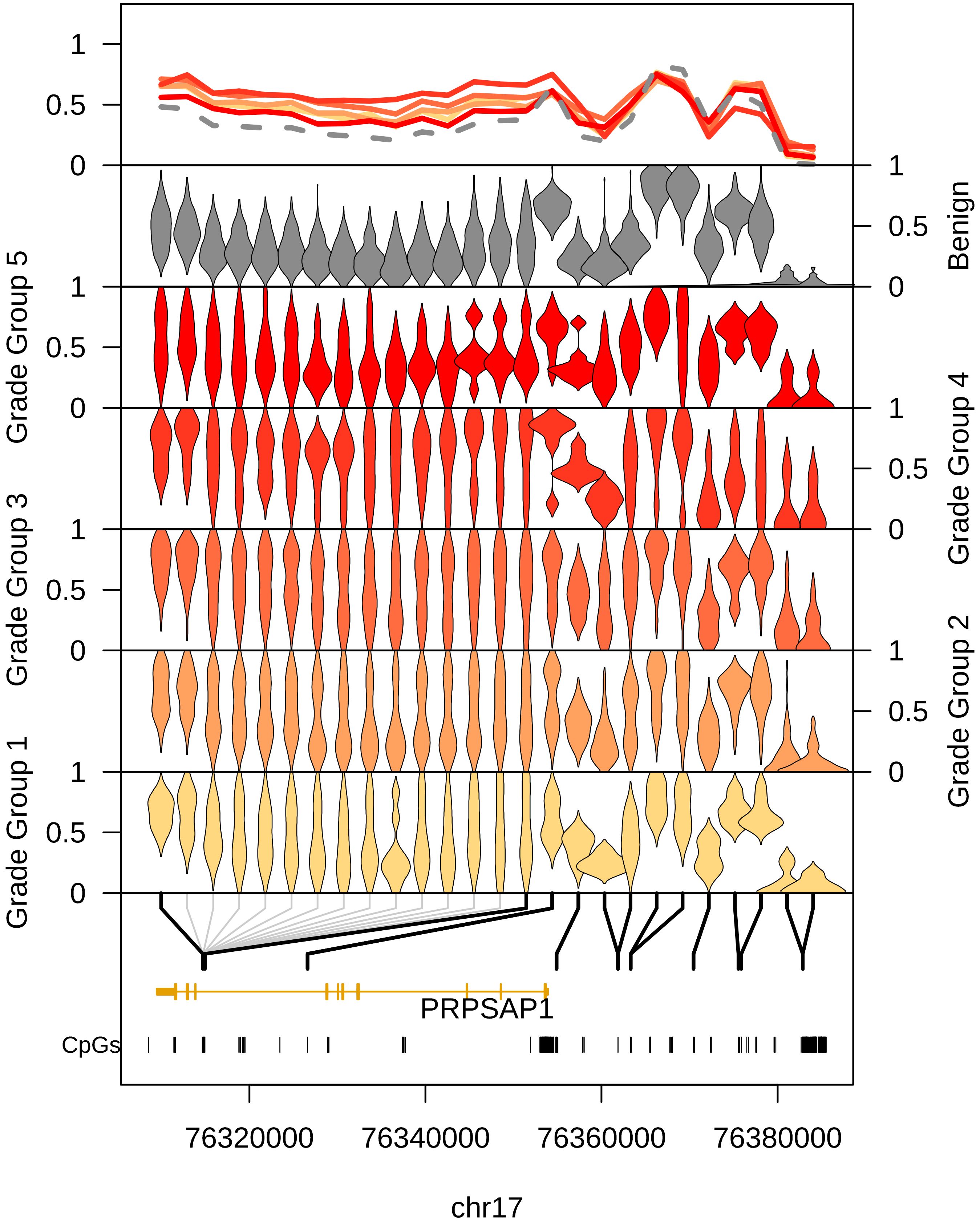

1
0.5
0
Benign
Grade Group 5
Grade Group 4
Grade Group 3
Grade Group 2
Grade Group 1
PRPSAP1
CpGs
76320000
76340000
76360000
76380000
chr17

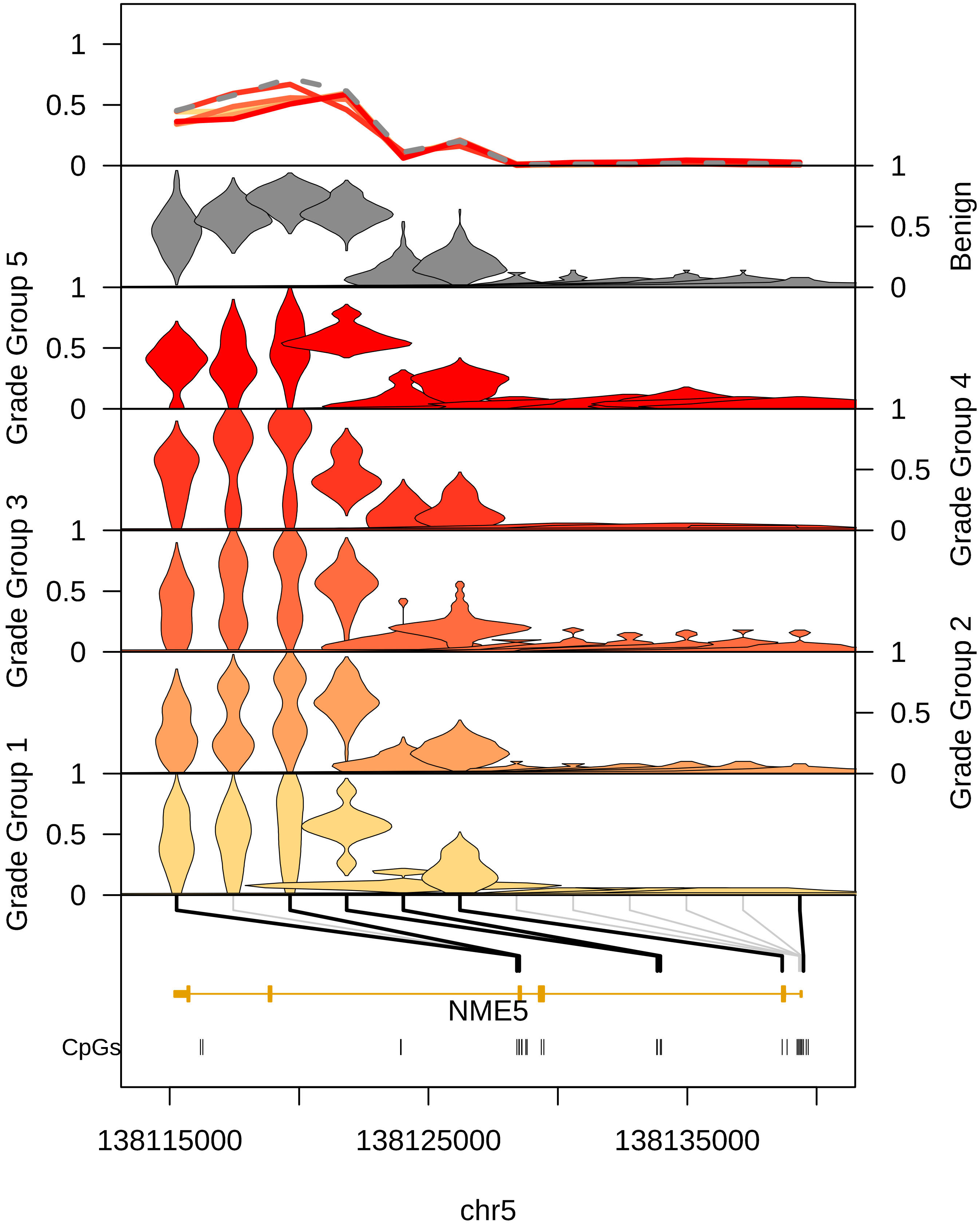

1
0.5
0
Benign
Grade Group 5
Grade Group 4
Grade Group 3
Grade Group 2
Grade Group 1
NME5
CpGs
138115000
138125000
138135000
chr5

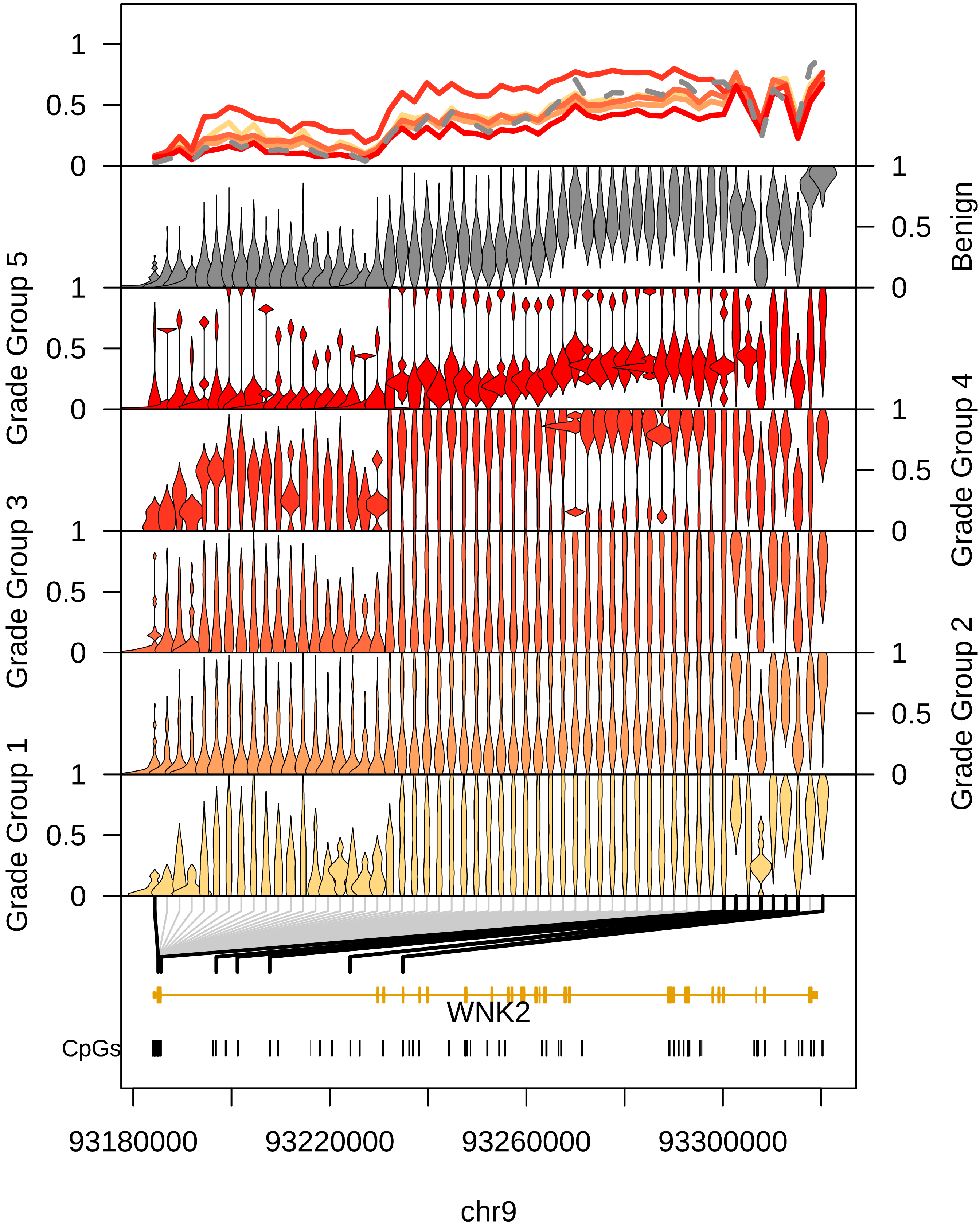

1
0.5
0
Benign
Grade Group 5
Grade Group 4
Grade Group 3
Grade Group 2
Grade Group 1
WNK2
CpGs
93180000
93220000
93260000
93300000
chr9

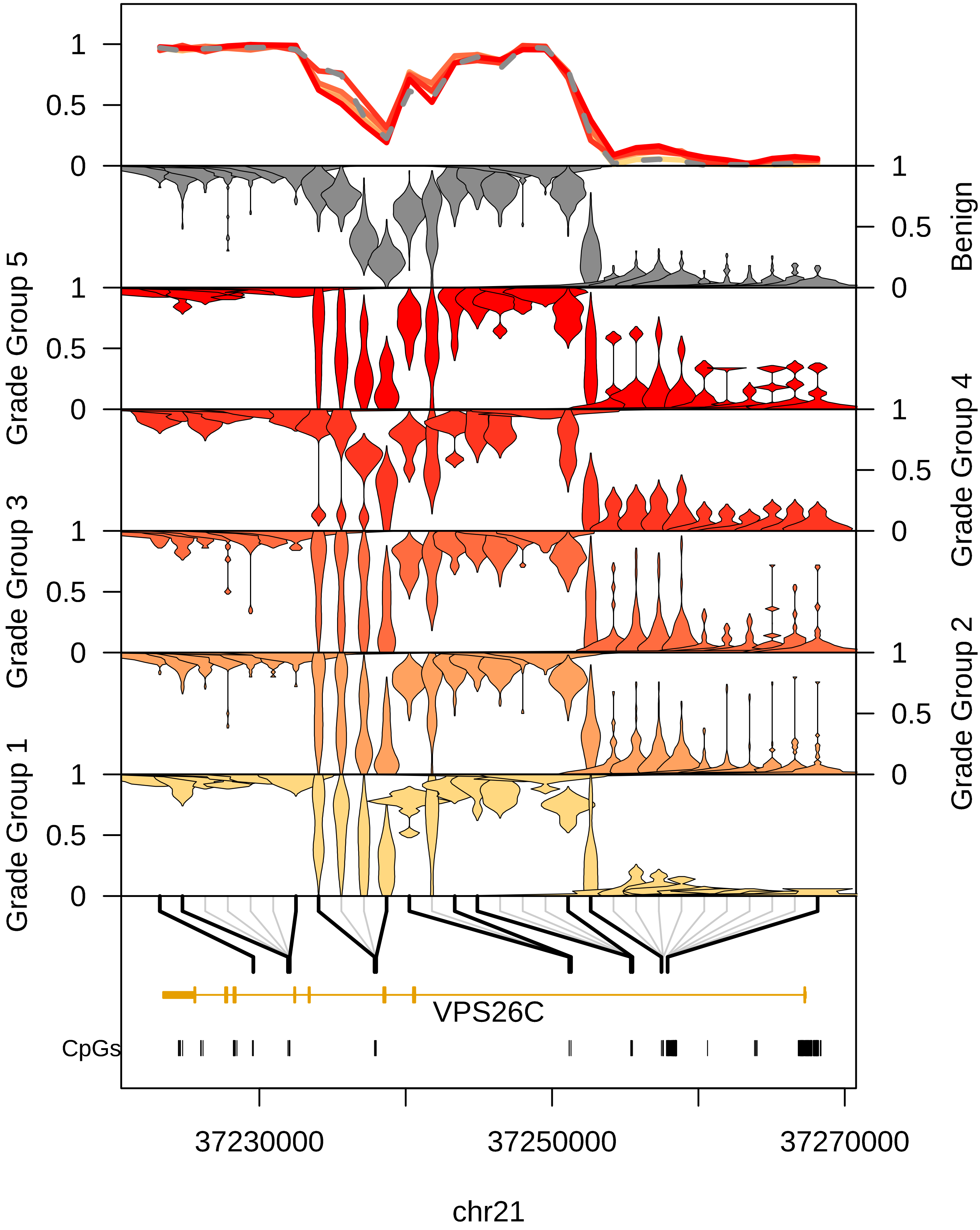

1
0.5
0
Benign
Grade Group 5
Grade Group 4
Grade Group 3
Grade Group 2
Grade Group 1
VPS26C
CpGs
37230000
37250000
37270000
chr21

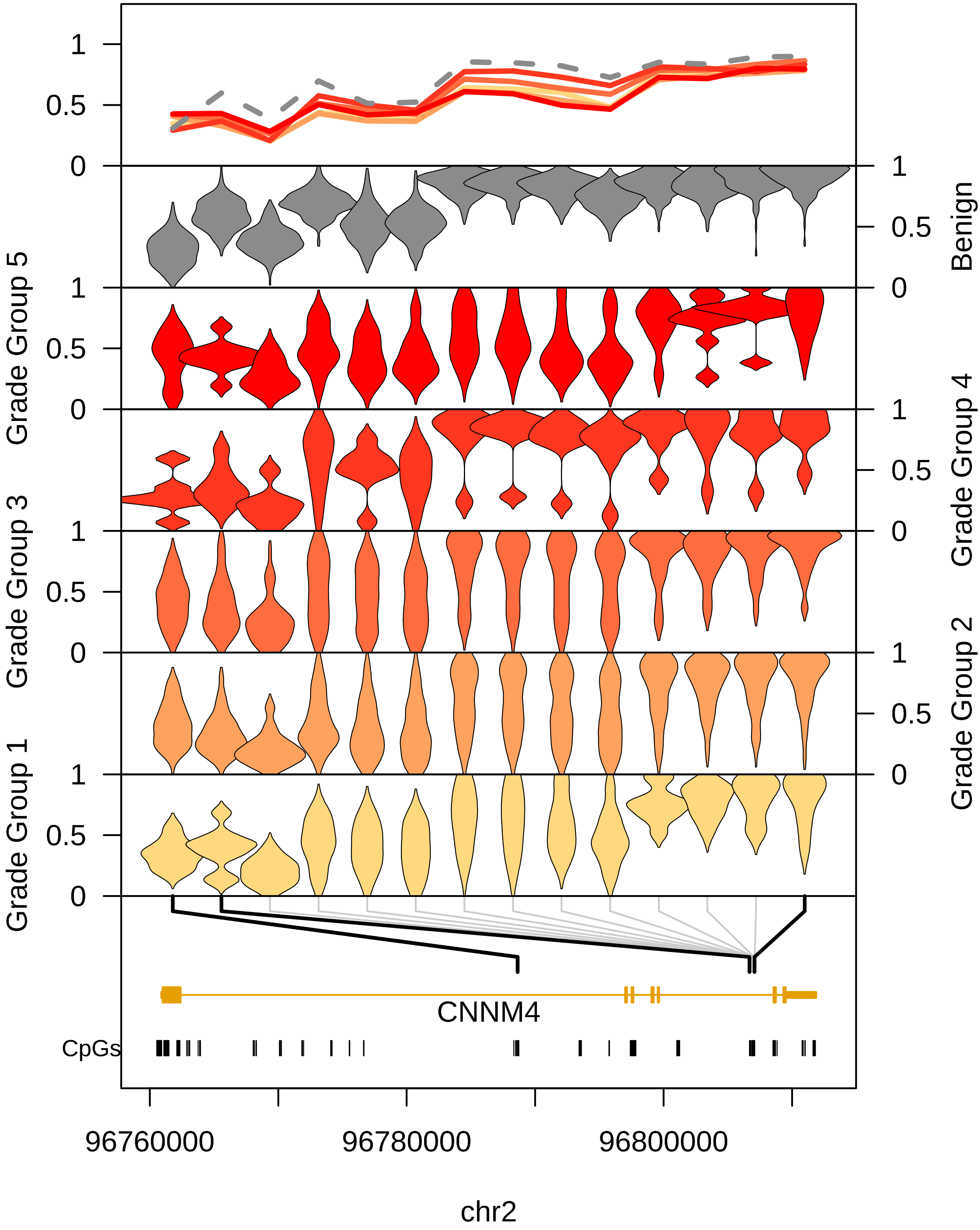

Benign
Grade Group 5
Grade Group 4
Grade Group 3
Grade Group 2
Grade Group 1
CNNM4
CpGs
96760000
96780000
96800000
chr2

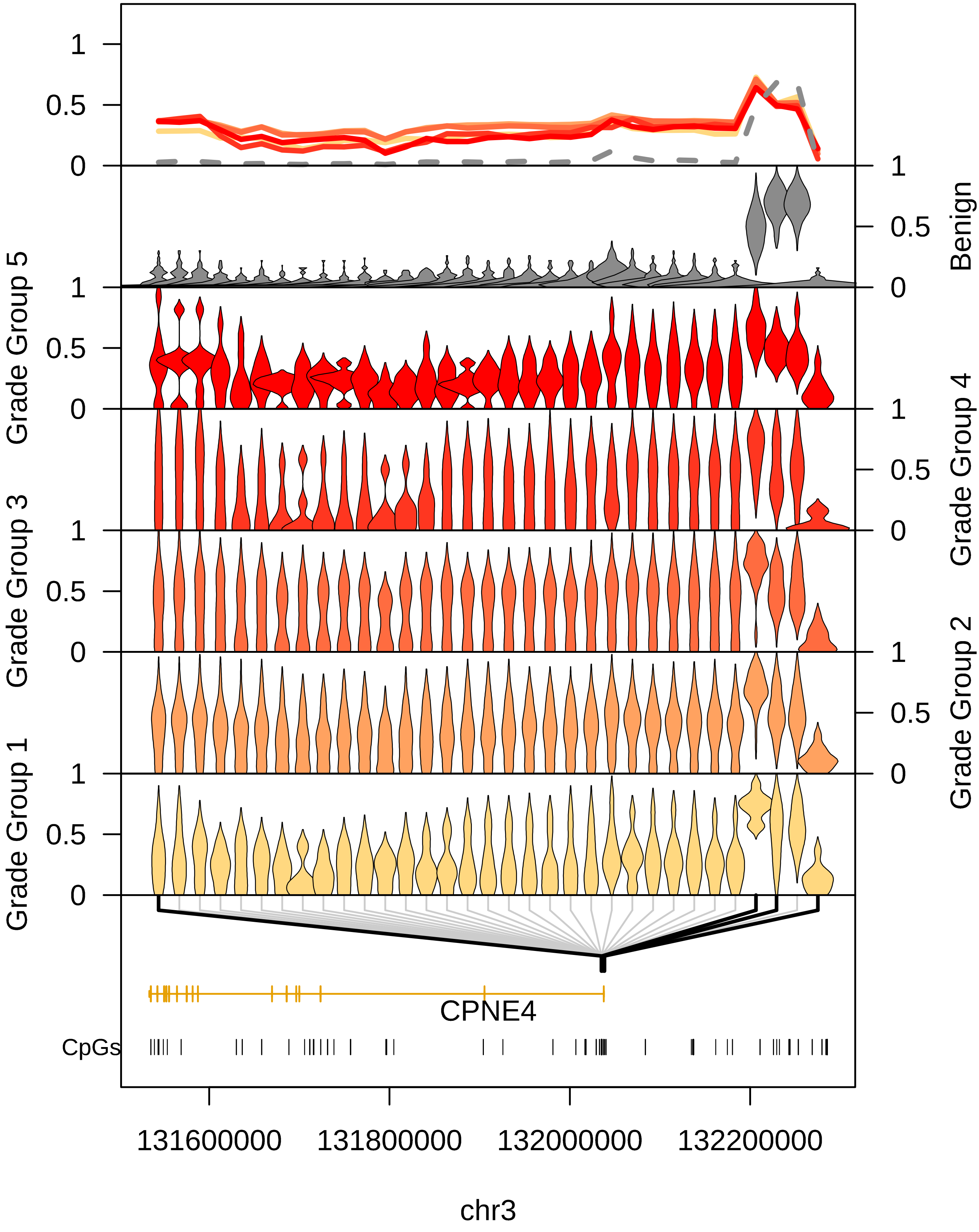

1
0.5
0
Benign
Grade Group 5
Grade Group 4
Grade Group 3
Grade Group 2
Grade Group 1
CPNE4
CpGs
131600000
131800000
132000000
132200000
chr3

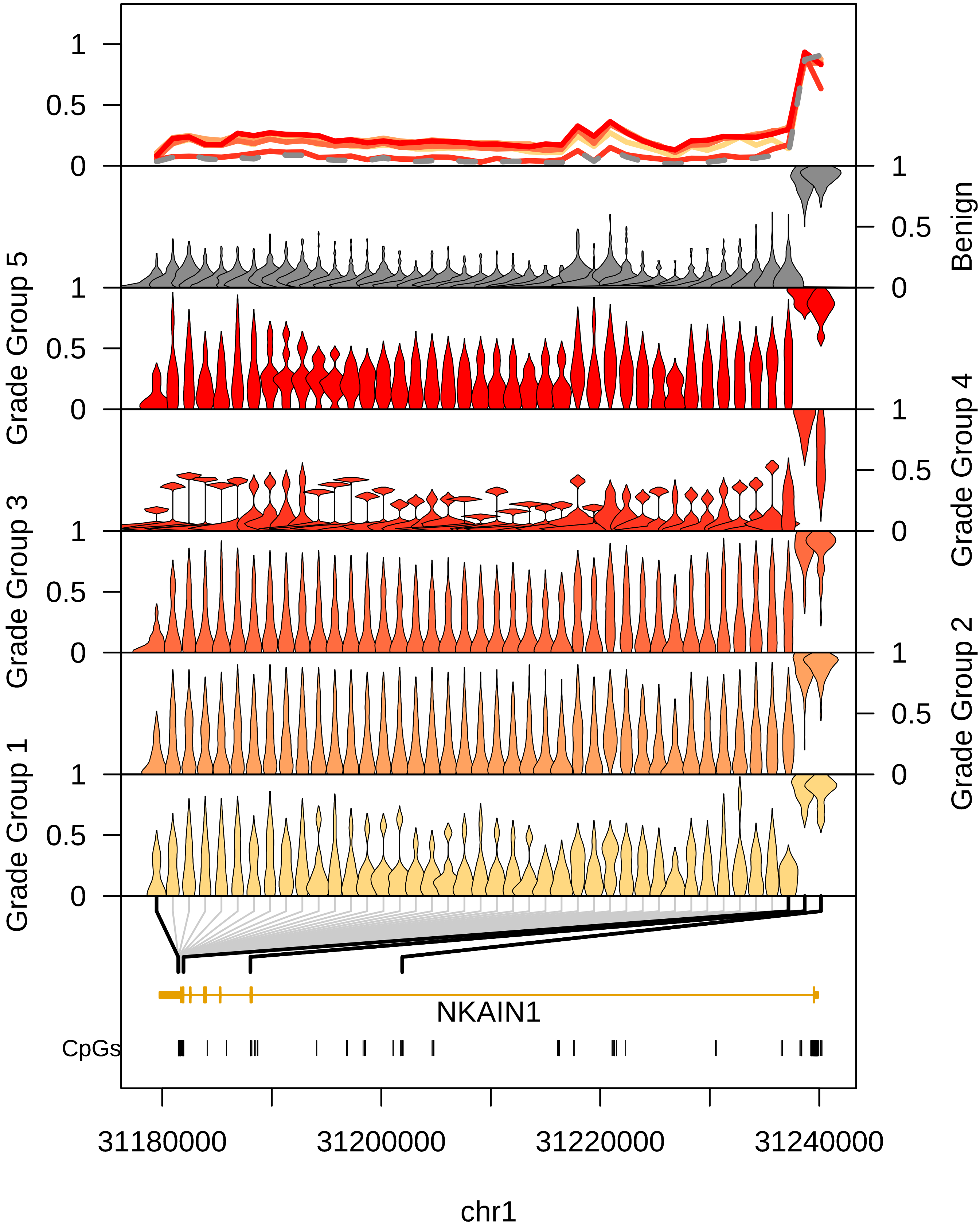

Benign
Grade Group 5
Grade Group 4
Grade Group 3
Grade Group 2
Grade Group 1
NKAIN1
CpGs
31180000
31200000
31220000
31240000
chr1

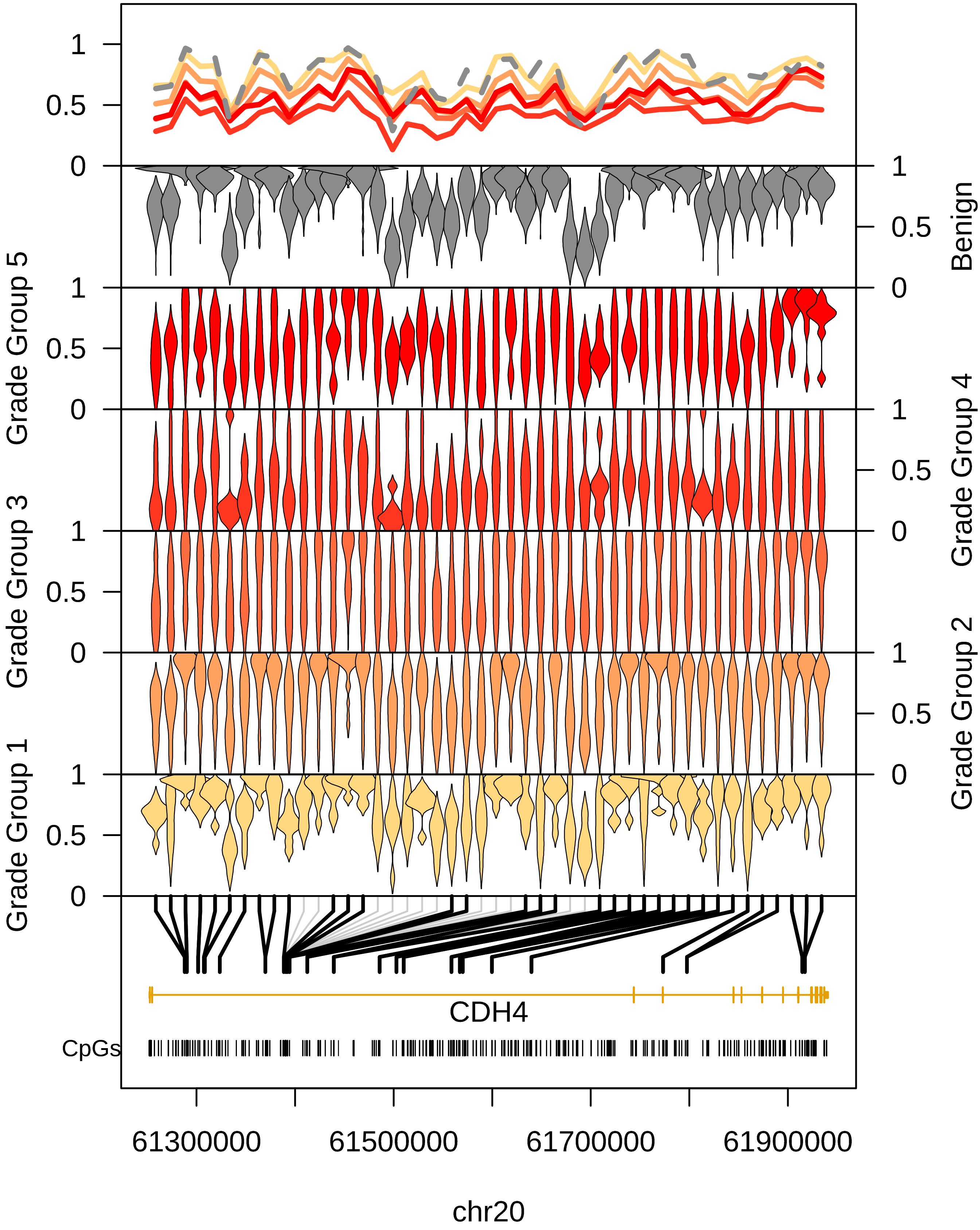

Benign
Grade Group 5
Grade Group 4
Grade Group 3
Grade Group 2
Grade Group 1
CDH4
CpGs
61300000
61500000
61700000
61900000
chr20

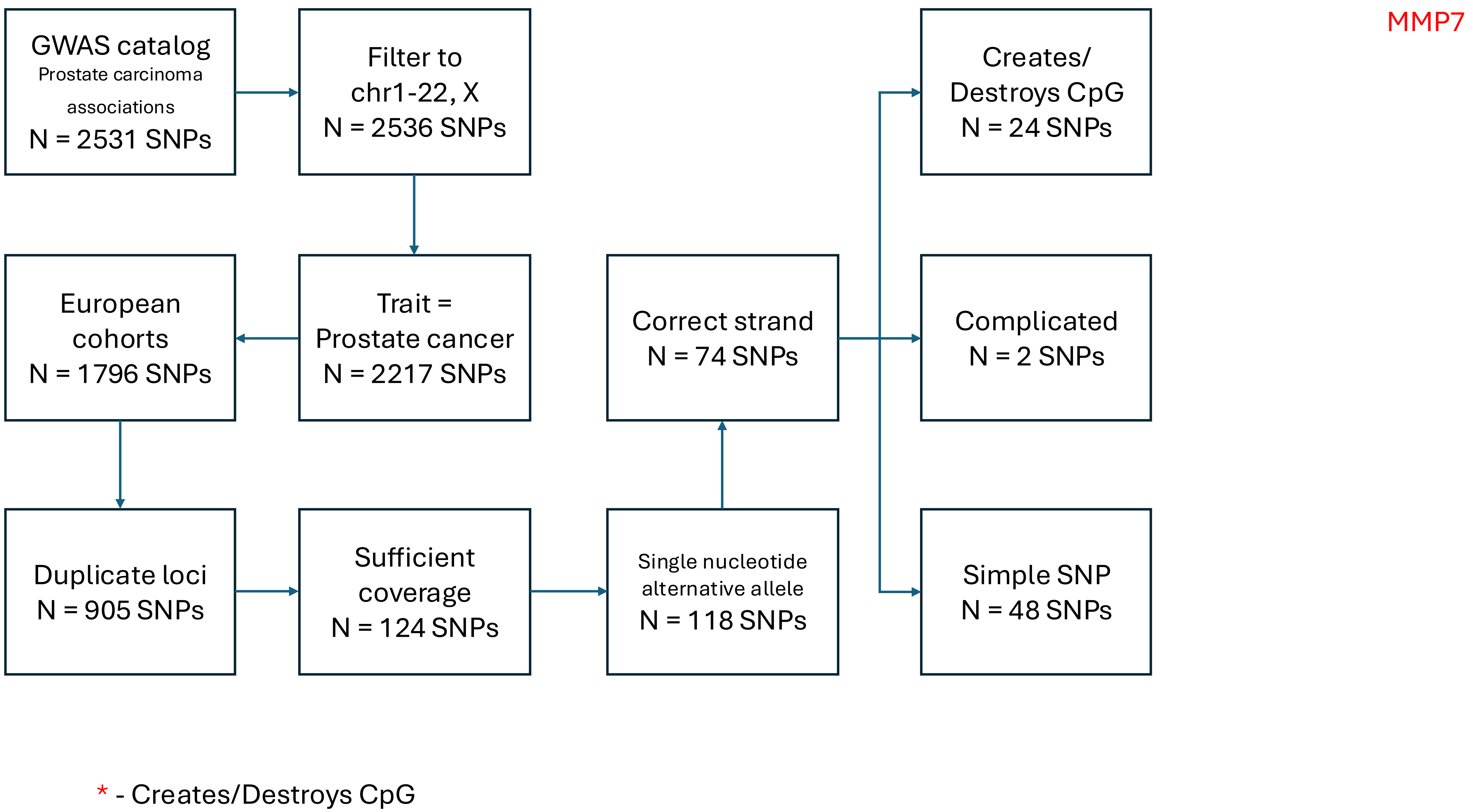

MMP7
GWAS catalog
Prostate carcinoma associations
N = 2531 SNPs
Filter to chr1-22, X
N = 2536 SNPs
Trait = Prostate cancer
N = 2217 SNPs
European cohorts
N = 1796 SNPs
Duplicate loci
N = 905 SNPs
Sufficient coverage
N = 124 SNPs
Single nucleotide alternative allele
N = 118 SNPs
Correct strand
N = 74 SNPs
Creates/ Destroys CpG
N = 24 SNPs
Complicated
N = 2 SNPs
Simple SNP
N = 48 SNPs
* - Creates/Destroys CpG

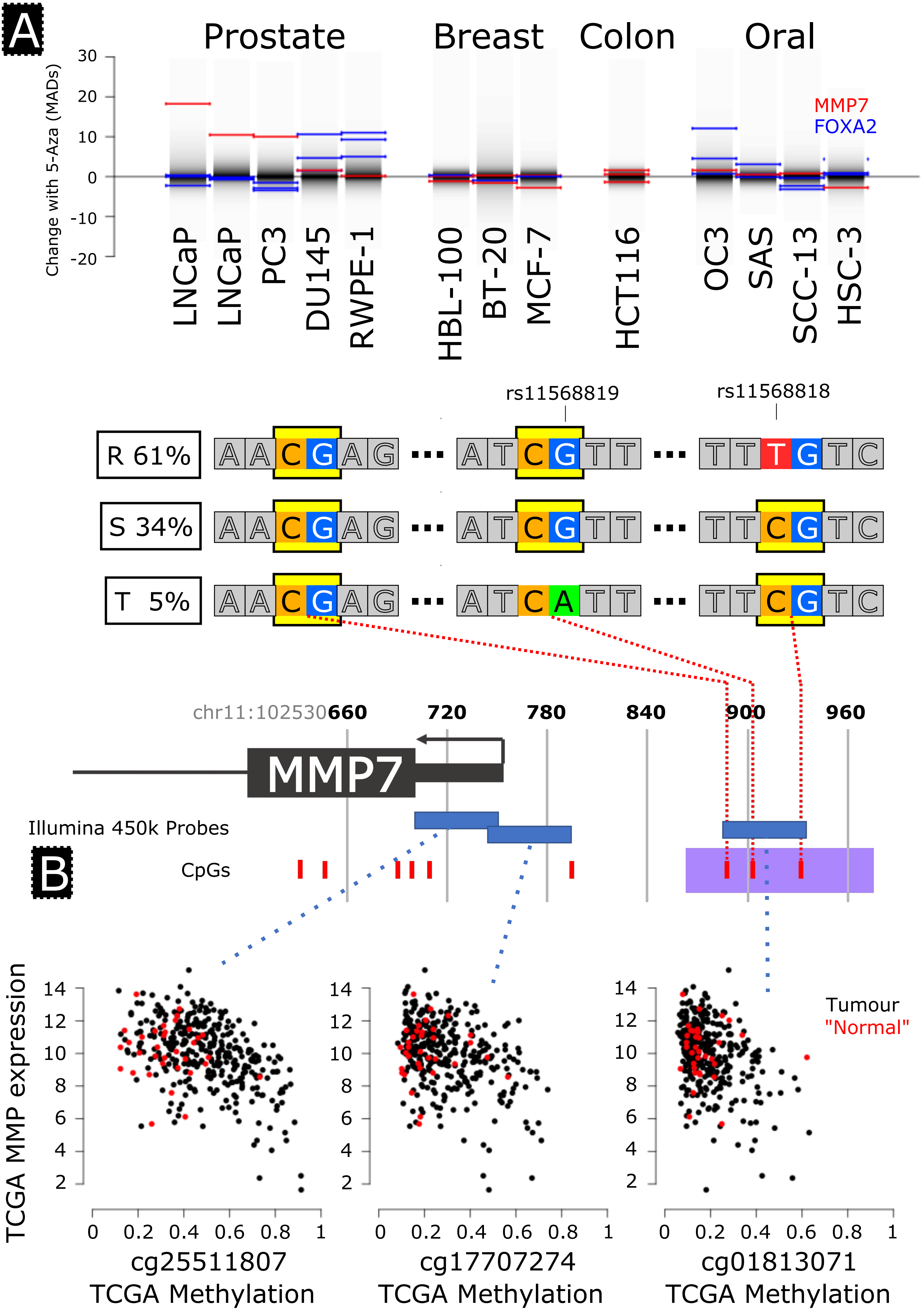
A
Prostate
Breast
Colon
Oral
Change with 5-Aza (MADs)
30
20
10
0
-10
-20
MMP7
FOXA2
LNCaP
LNCaP
PC3
DU145
RWPE-1
HBL-100
BT-20
MCF-7
HCT116
OC3
SAS
SCC-13
HSC-3
rs11568819
rs11568818
R 61%
S 34%
T 5%
A A C G A G
A T C G T T
T T T G T C
A A C G A G
A T C G T T
T T C G T C
A A C G A G
A T C A T T
T T C G T C
chr11:102530660
720
780
840
900
960
MMP7
Illumina 450k Probes
B
CpGs
TCGA MMP expression
Tumour
"Normal"
cg25511807
cg17707274
cg01813071
TCGA Methylation
TCGA Methylation
TCGA Methylation

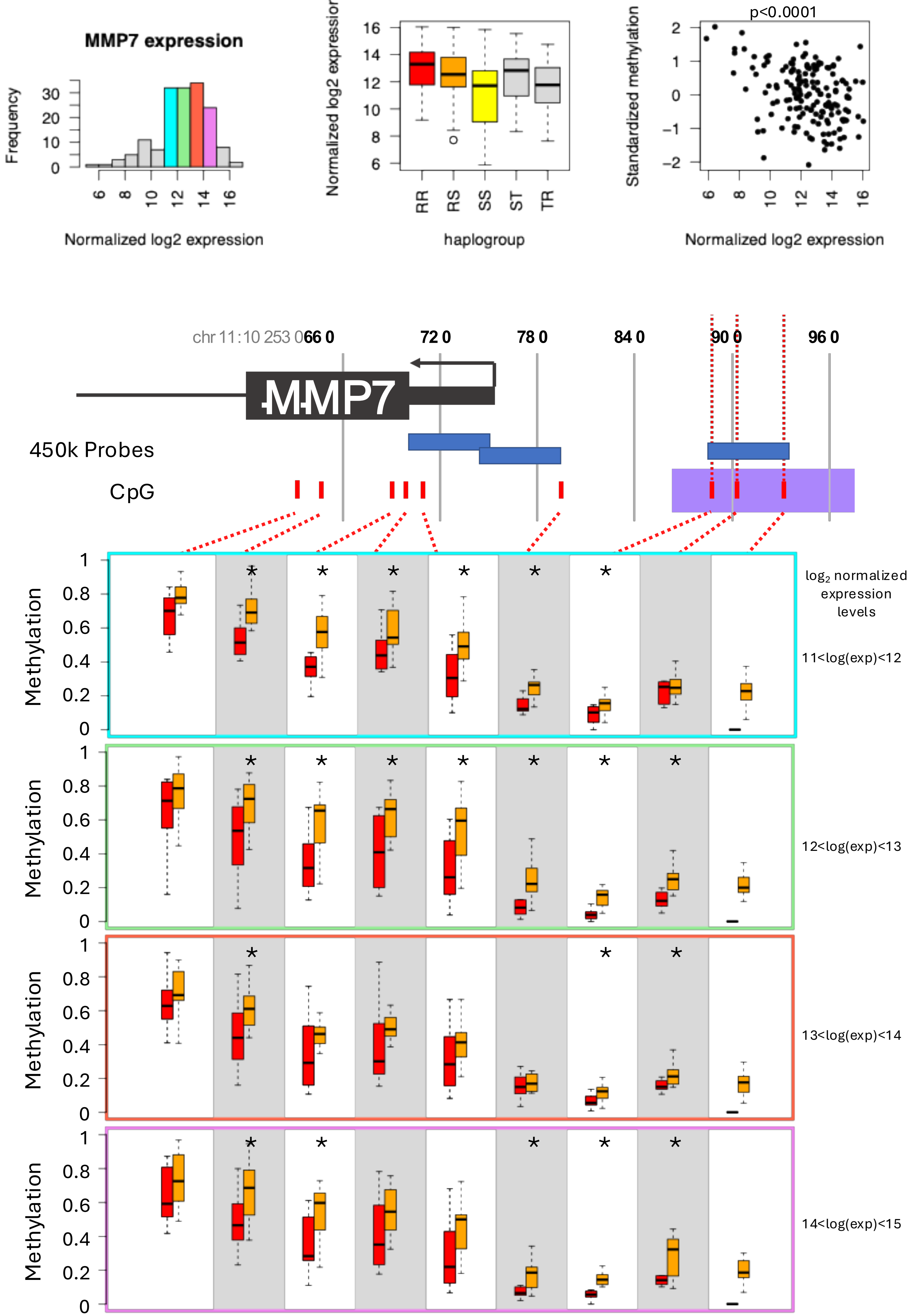

MMP7 expression
Frequency
Normalized log2 expression
haplogroup
RR
RS
SS
ST
TR
Standardized methylation
p<0.0001
chr 11:10 253 066 0
72 0
78 0
84 0
90 0
96 0
MMP7
450k Probes
CpG
Methylation
log2 normalized expression levels
11<log(exp)<12
12<log(exp)<13
13<log(exp)<14
14<log(exp)<15

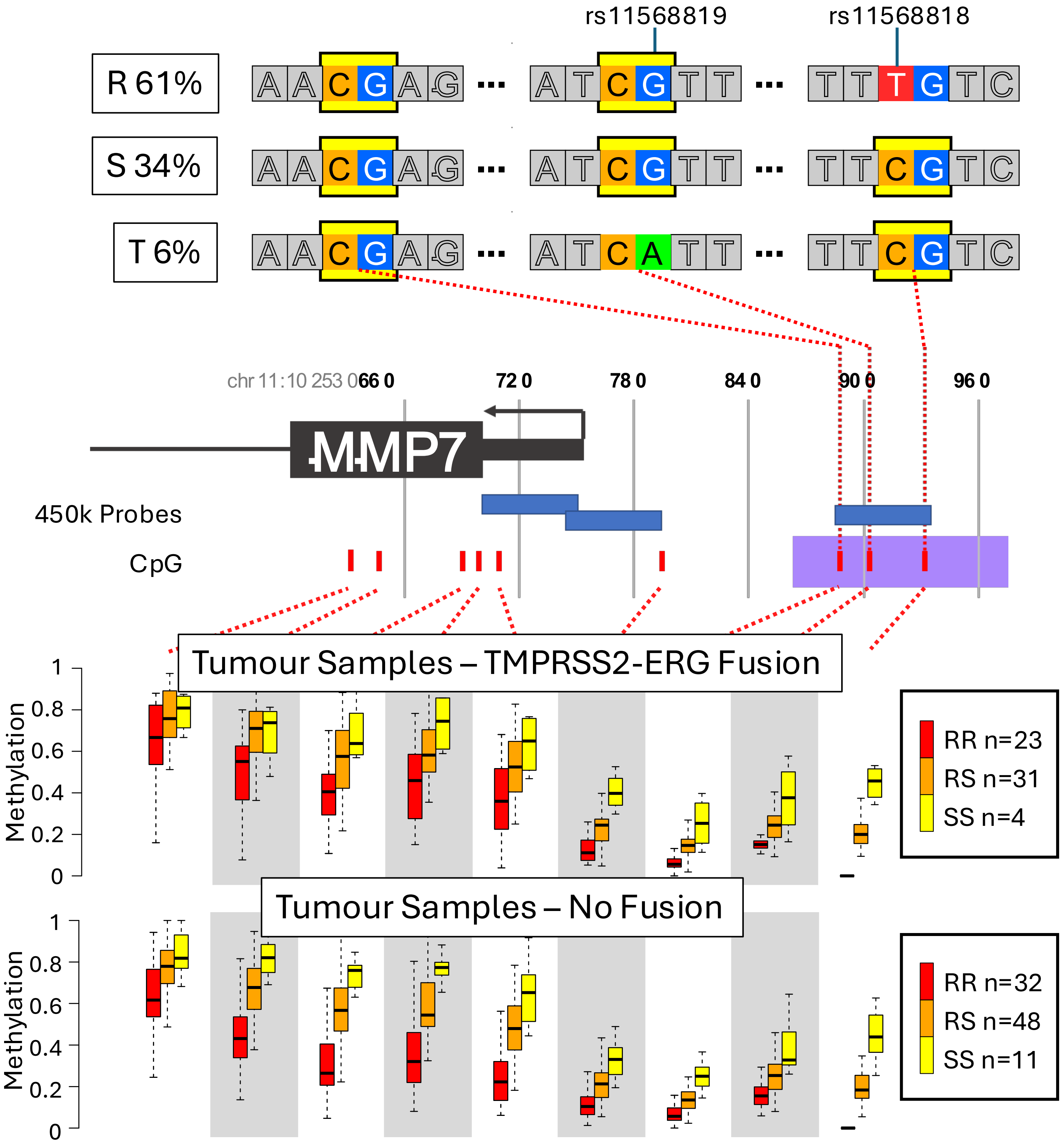

rs11568819
rs11568818
R 61%
S 34%
T 6%
chr 11:10 253 066 0
72 0
78 0
84 0
90 0
96 0
MMP7
450k Probes
CpG
Tumour Samples – TMPRSS2-ERG Fusion
Methylation
RR n=23
RS n=31
SS n=4
Tumour Samples – No Fusion
Methylation
RR n=32
RS n=48
SS n=11

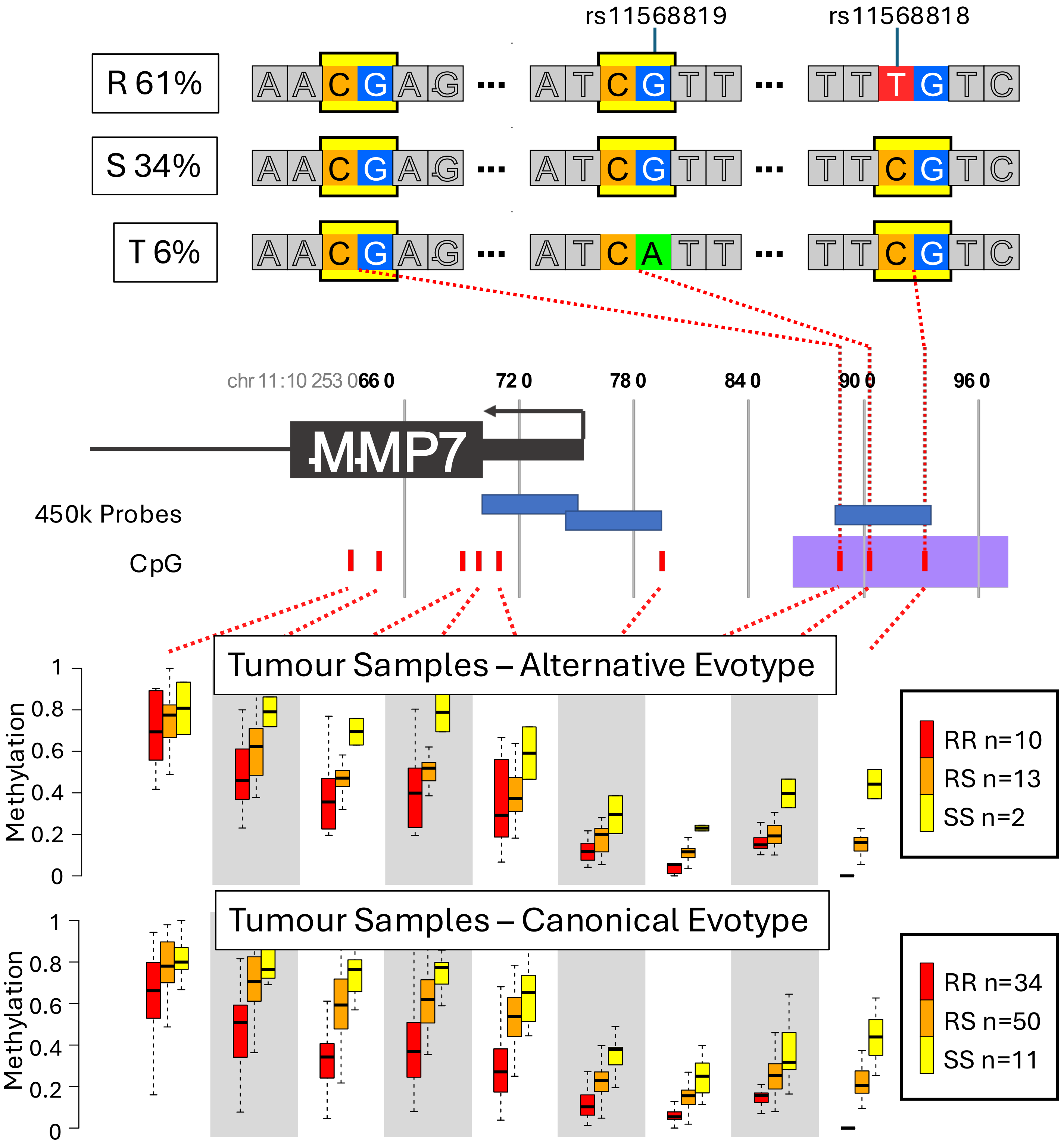
rs11568819
rs11568818
R 61%
S 34%
T 6%
chr 11:10 253 066 0
72 0
78 0
84 0
90 0
96 0
MMP7
450k Probes
CpG
Tumour Samples – Alternative Evotype
Methylation
RR n=10
RS n=13
SS n=2
Tumour Samples – Canonical Evotype
RR n=34
RS n=50
SS n=11

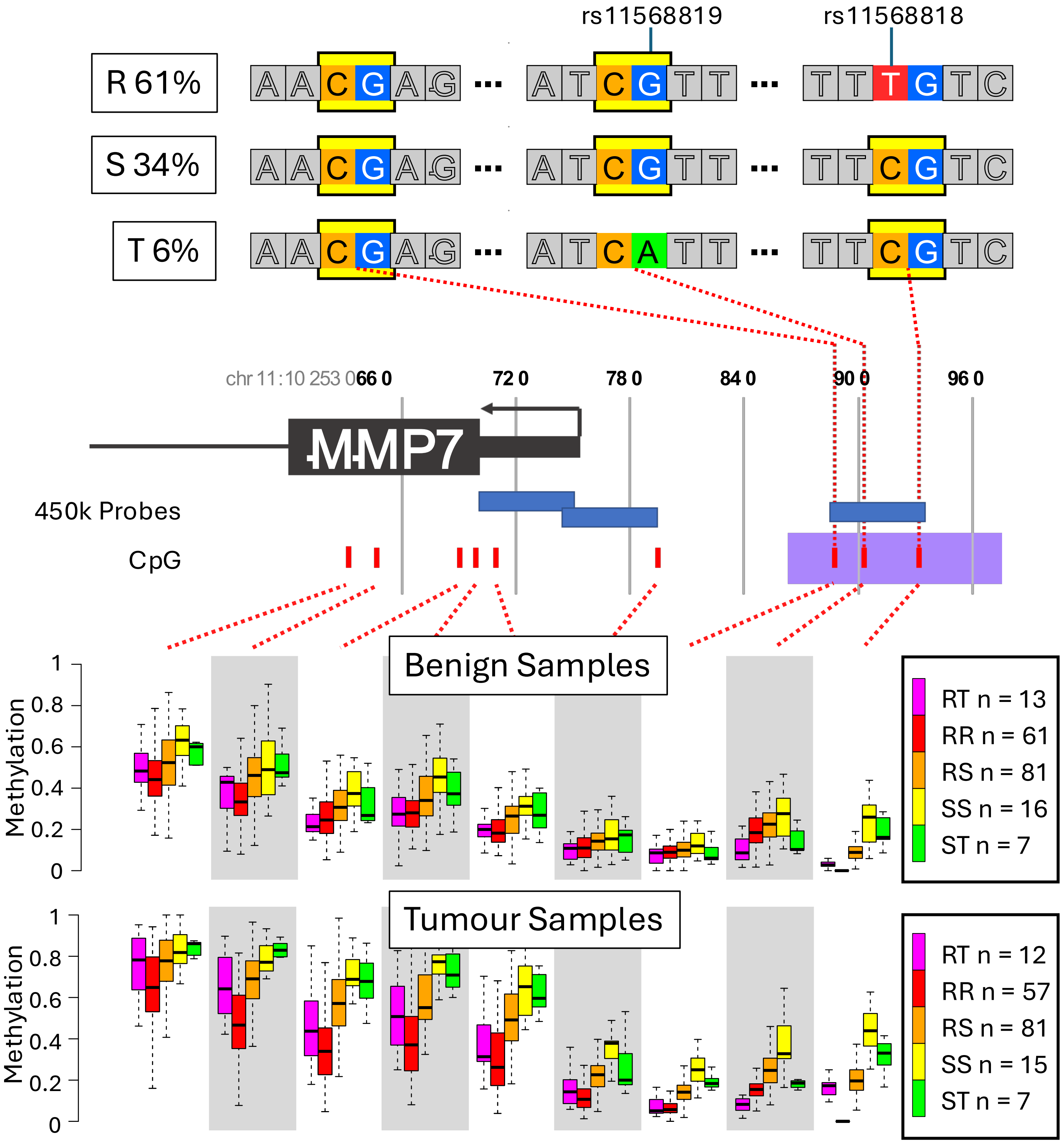

rs11568819
rs11568818
R 61%
S 34%
T 6%
chr 11:10 253 066 0
72 0
78 0
84 0
90 0
96 0
MMP7
450k Probes
CpG
Benign Samples
Methylation
RT n = 13
RR n = 61
RS n = 81
SS n = 16
ST n = 7
Tumour Samples
RT n = 12
RR n = 57
RS n = 81
SS n = 15
ST n = 7

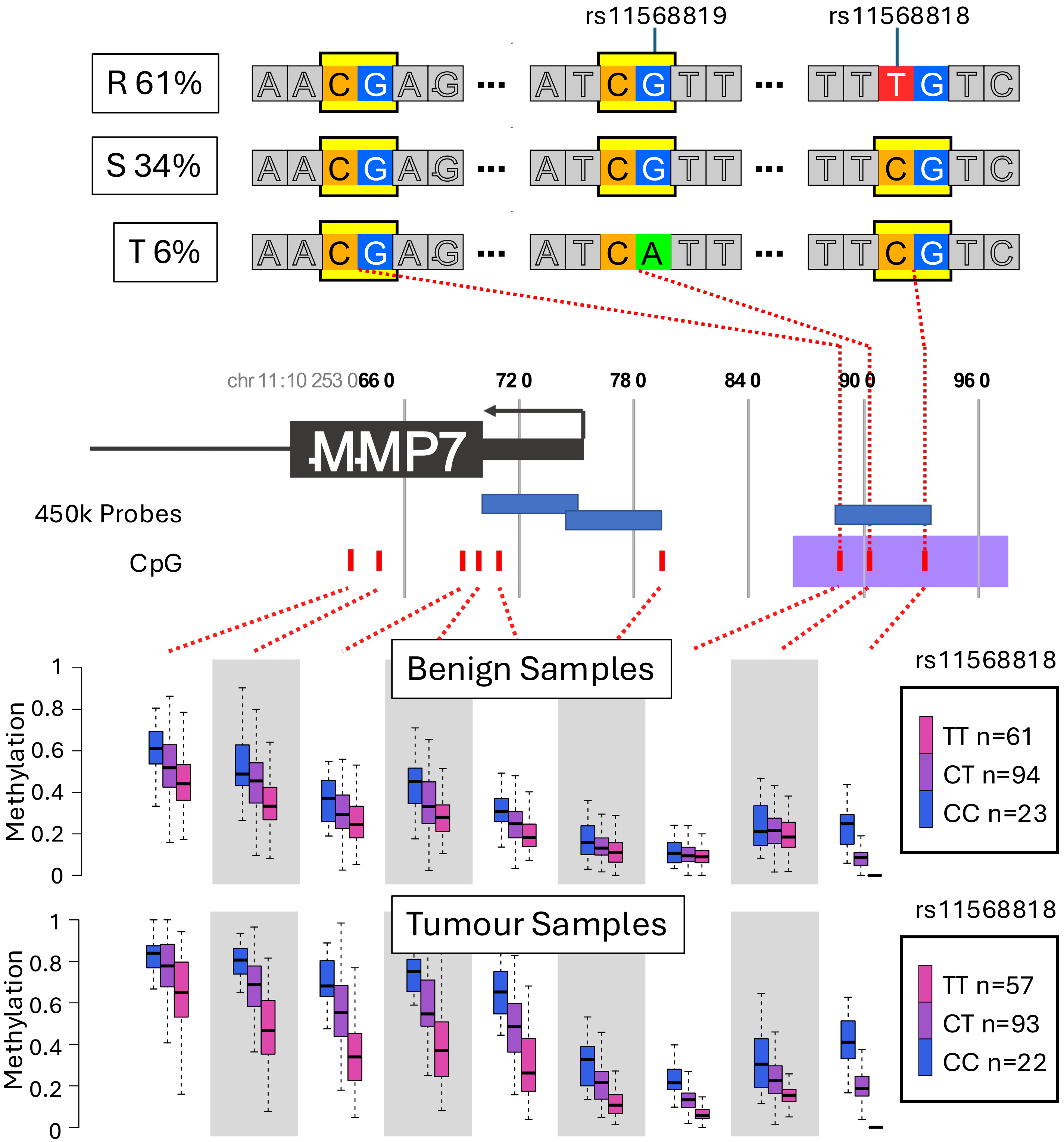

rs11568819
rs11568818
R 61%
S 34%
T 6%
chr 11:10 253 066 0
72 0
78 0
84 0
90 0
96 0
MMP7
450k Probes
CpG
Benign Samples
Tumour Samples
Methylation
rs11568818
TT n=61
CT n=94
CC n=23
TT n=57
CT n=93
CC n=22

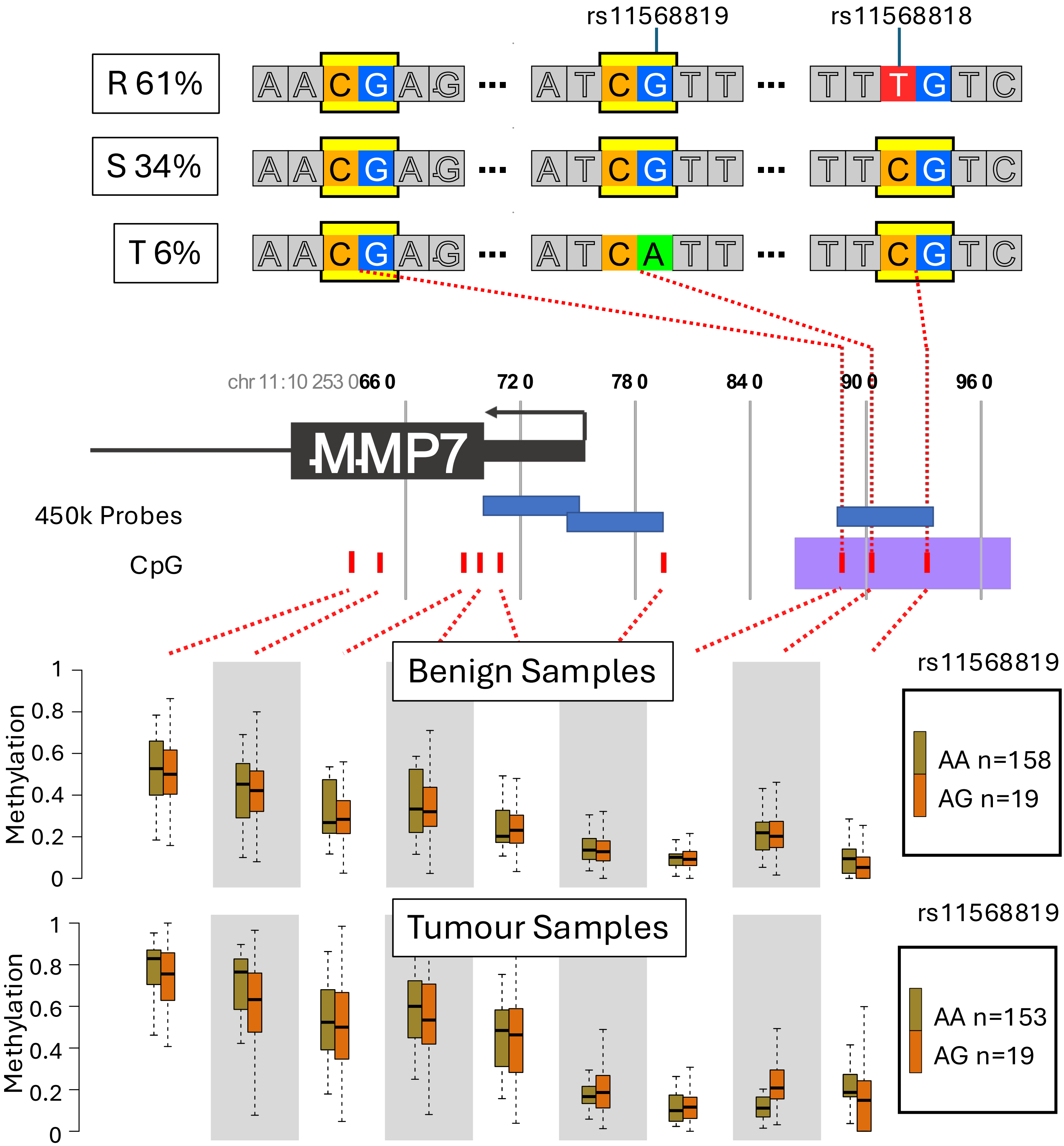
rs11568819
rs11568818
R 61%
S 34%
T 6%
A A C G A G ··· A T C G T T ··· T T T G T C
A A C G A G ··· A T C G T T ··· T T C G T C
A A C G A G ··· A T C A T T ··· T T C G T C
chr 11:10 253 066 0
72 0
78 0
84 0
90 0
96 0
MMP7
450k Probes
CpG
Benign Samples
Methylation
1
0.8
0.6
0.4
0.2
0
rs11568819
AA n=158
AG n=19
Tumour Samples
rs11568819
AA n=153
AG n=19

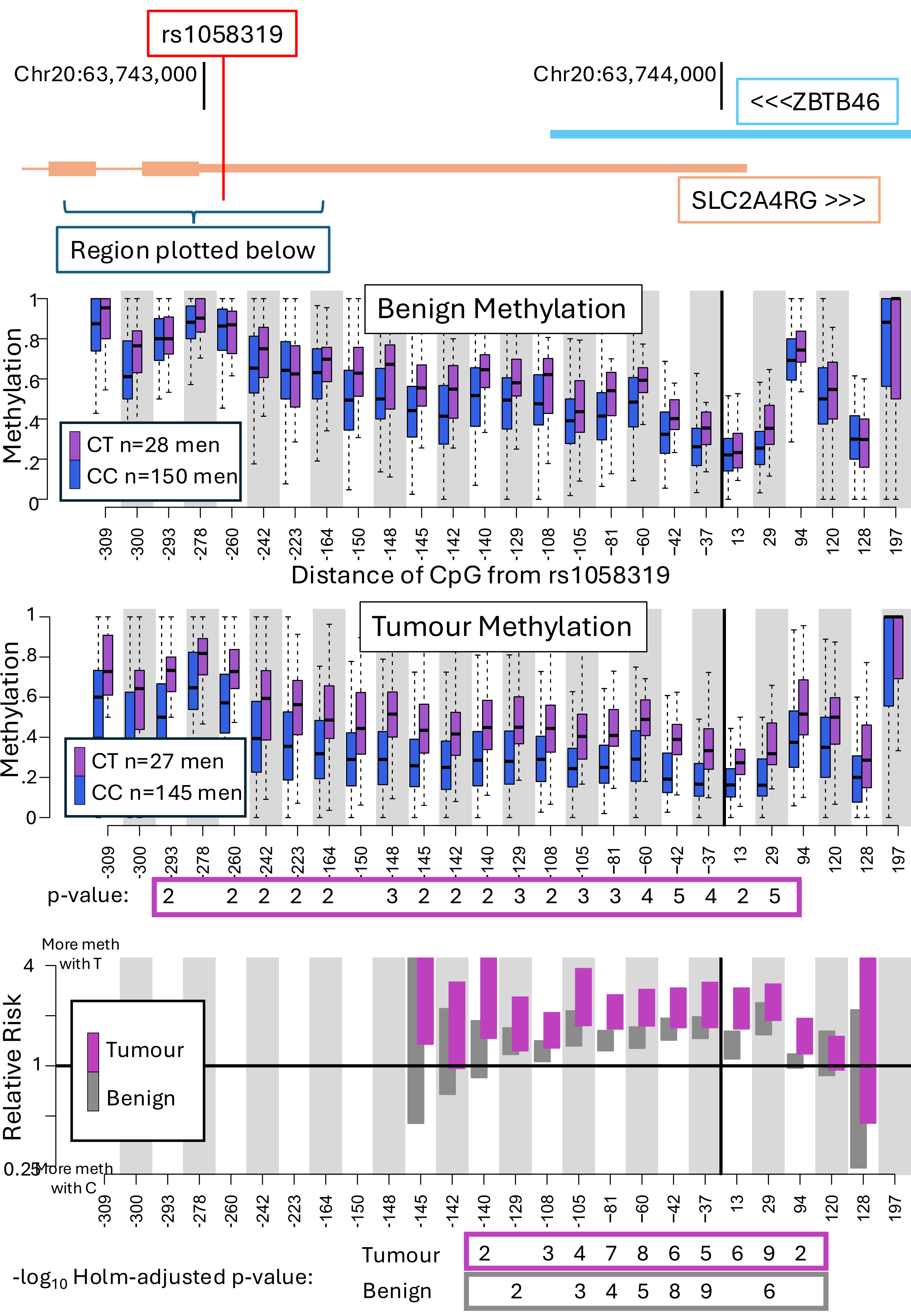
rs1058319
Chr20:63,743,000
Chr20:63,744,000
<<<ZBTB46
SLC2A4RG >>>
Region plotted below
Benign Methylation
Methylation
CT n=28 men
CC n=150 men
-309 -300 -293 -278 -260 -242 -223 -164 -150 -148 -145 -142 -140 -129 -108 -105 -81 -60 -42 -37 13 29 94 120 128 197
Distance of CpG from rs1058319
Tumour Methylation
CT n=27 men
CC n=145 men
p-value: 2 2 2 2 2 3 2 2 2 3 2 3 3 4 5 4 2 5
More meth with T
More meth with C
Relative Risk
Tumour
Benign
-log10 Holm-adjusted p-value:
Tumour 2 3 4 7 8 6 5 6 9 2
Benign 2 3 4 5 8 9 6

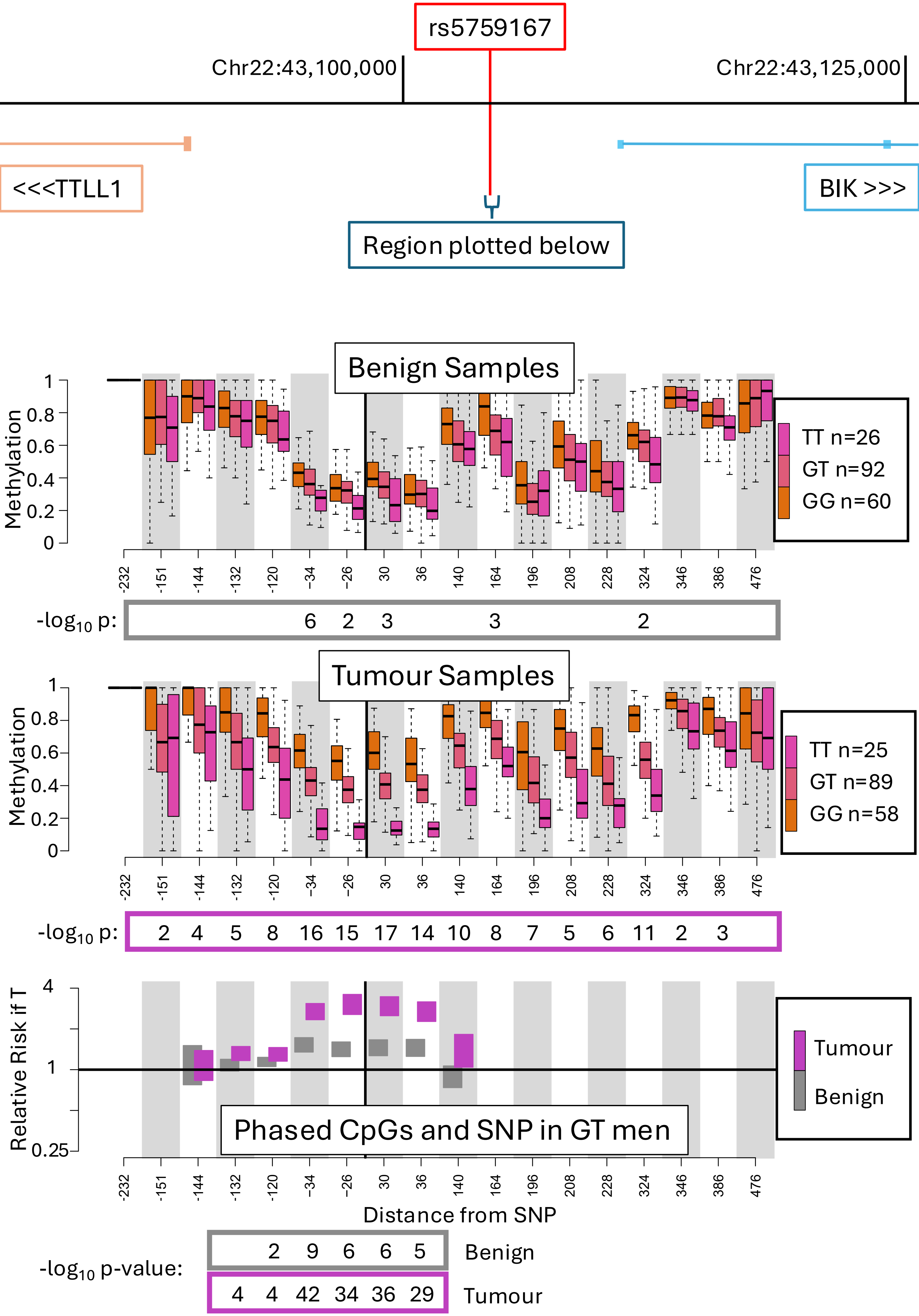

rs5759167
Chr22:43,100,000
Chr22:43,125,000
<<<TTLL1
BIK >>>
Region plotted below
Benign Samples
Methylation
TT n=26
GT n=92
GG n=60
-log10 p:
6 2 3 3 2
Tumour Samples
TT n=25
GT n=89
GG n=58
-log10 p:
2 4 5 8 16 15 17 14 10 8 7 5 6 11 2 3
Relative Risk if T
Phased CpGs and SNP in GT men
Tumour
Benign
Distance from SNP
-232 -151 -144 -132 -120 -34 -26 30 36 140 164 196 208 228 324 346 386 476
-log10 p-value:
2 9 6 6 5
Benign
4 4 42 34 36 29
Tumour

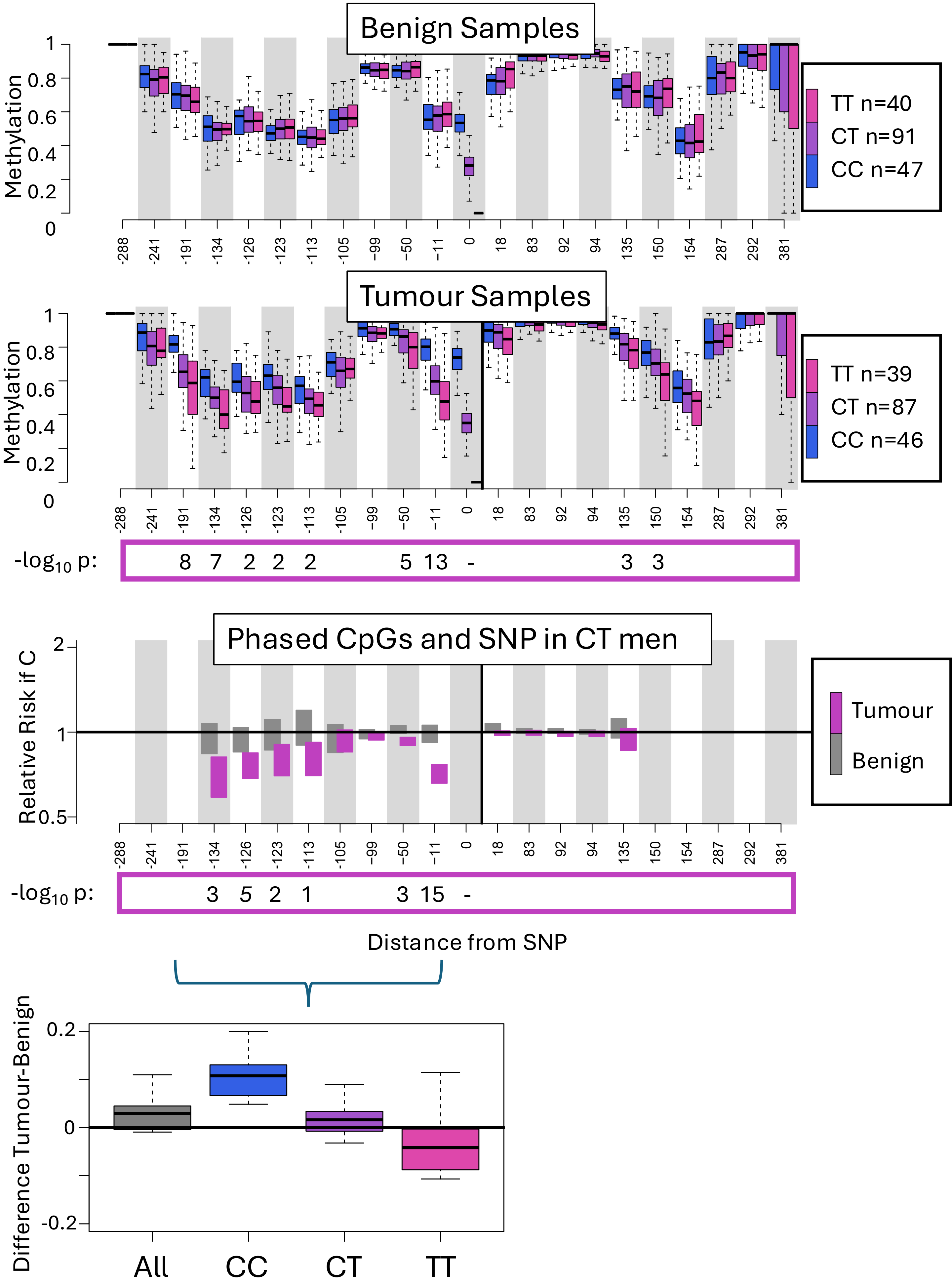


rs10866527 at chr5-1891686 (CTD-2194D22, IRX4?)

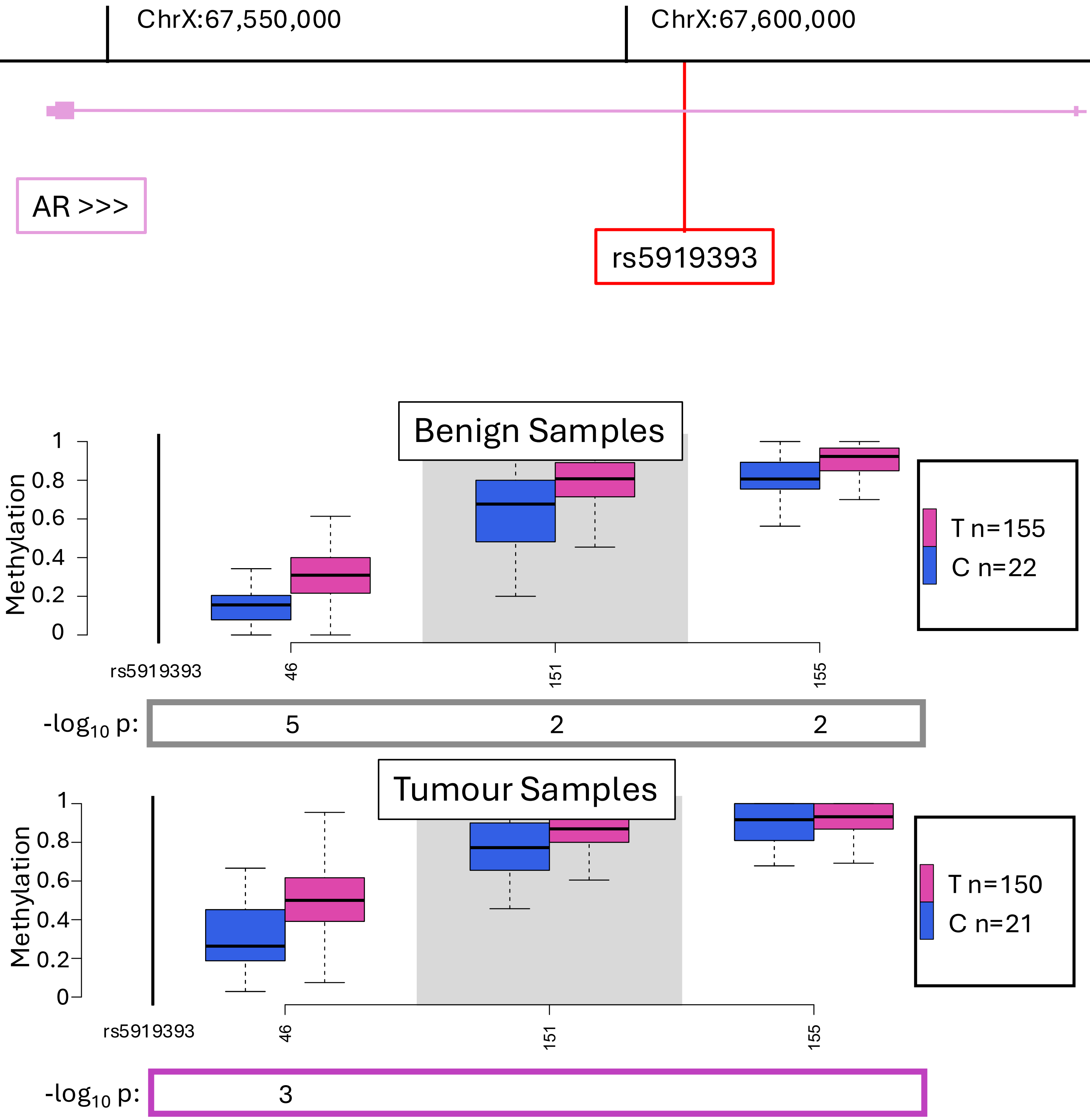
ChrX:67,550,000
ChrX:67,600,000
AR >>>
rs5919393
Benign Samples
Methylation
1
0.8
0.6
0.4
0.2
0
T n=155
C n=22
rs5919393
46
151
155
-log10 p:
5
2
2
Tumour Samples
Methylation
1
0.8
0.6
0.4
0.2
0
T n=150
C n=21
rs5919393
46
151
155
-log10 p:
3

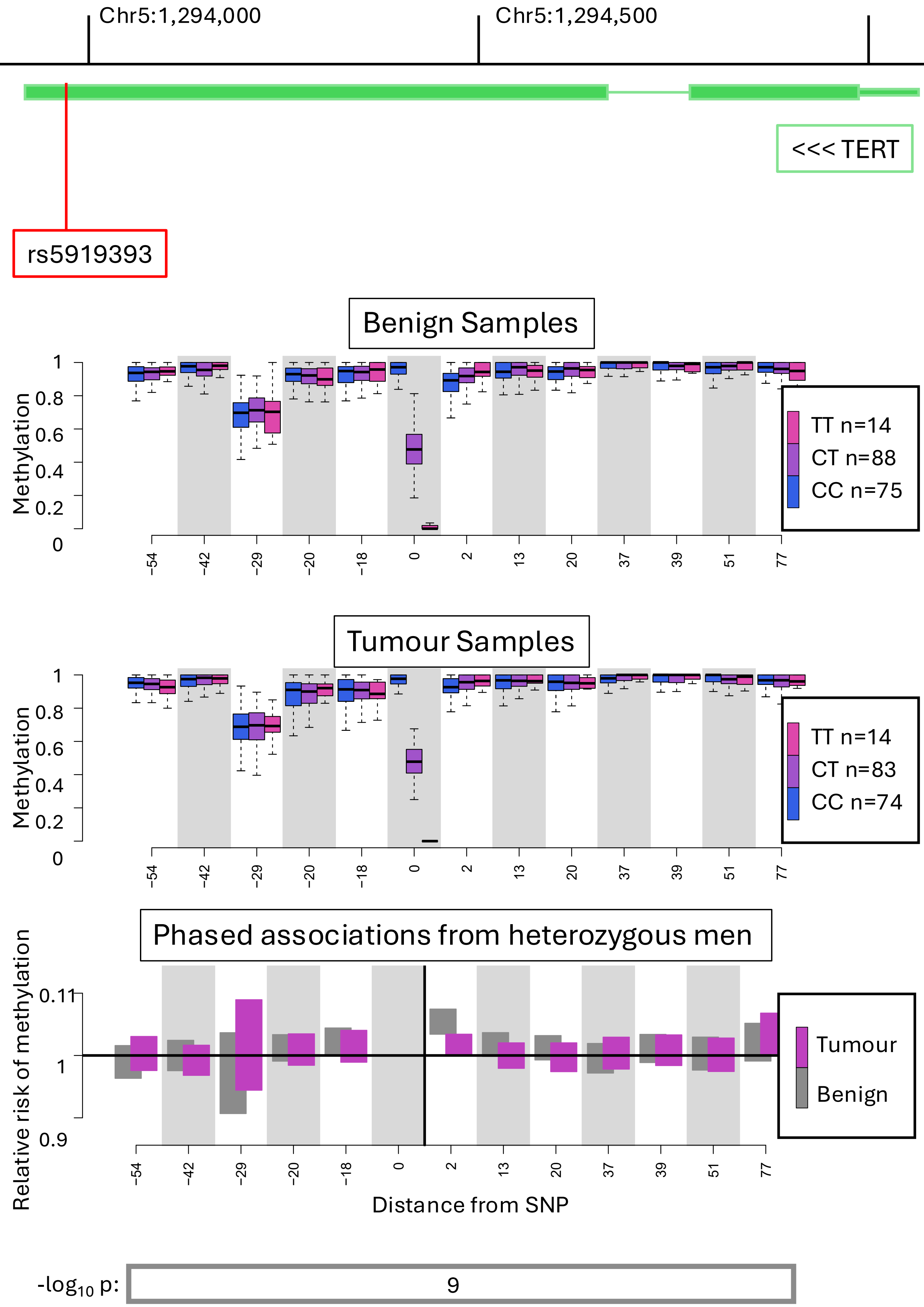

Chr5:1,294,000
Chr5:1,294,500
<<< TERT
rs5919393
Benign Samples
Methylation
1
0.8
0.6
0.4
0.2
0
-54 -42 -29 -20 -18 0 2 13 20 37 39 51 77
TT n=14
CT n=88
CC n=75
Tumour Samples
TT n=14
CT n=83
CC n=74
Phased associations from heterozygous men
Relative risk of methylation
0.11
1
0.9
Tumour
Benign
Distance from SNP
-log10 p:
9

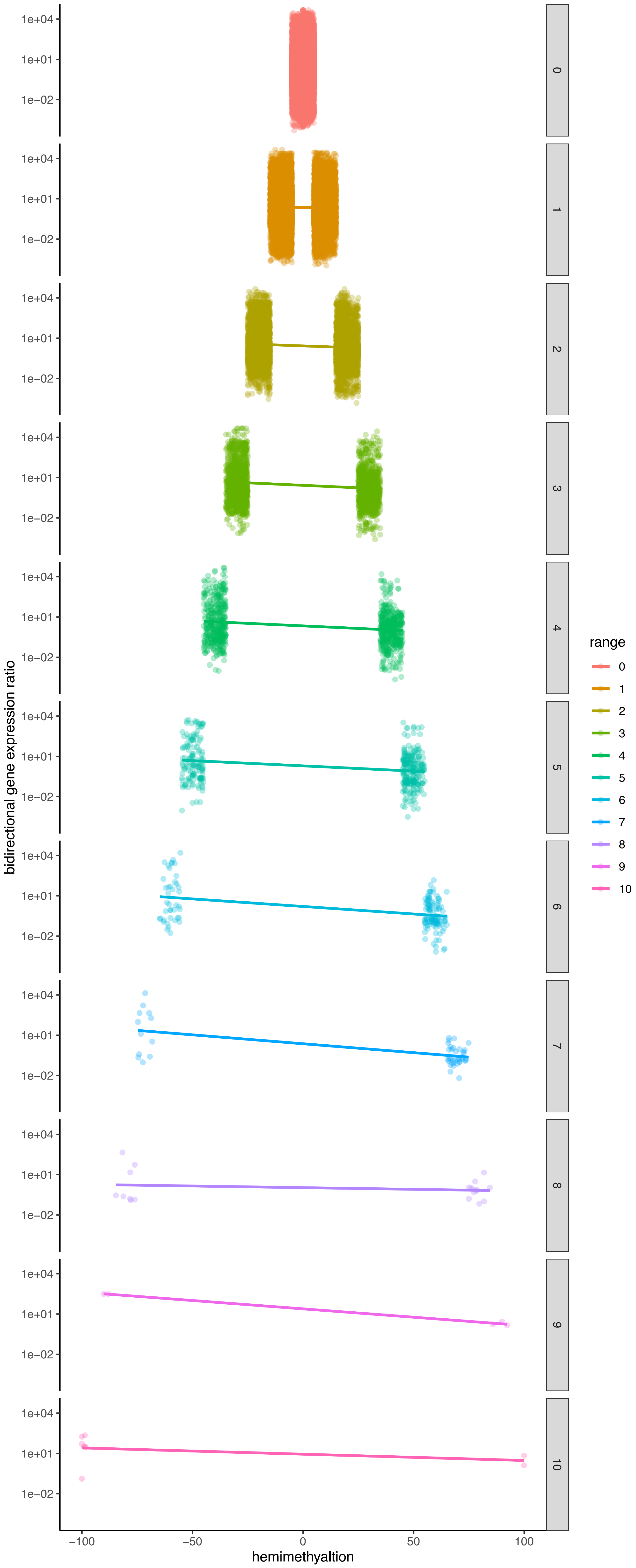

bidirectional gene expression ratio
hemimethyaltion
1e+04
1e+01
1e−02
−100
−50
0
50
100
0
1
2
3
4
5
6
7
8
9
10
range
0
1
2
3
4
5
6
7
8
9
10

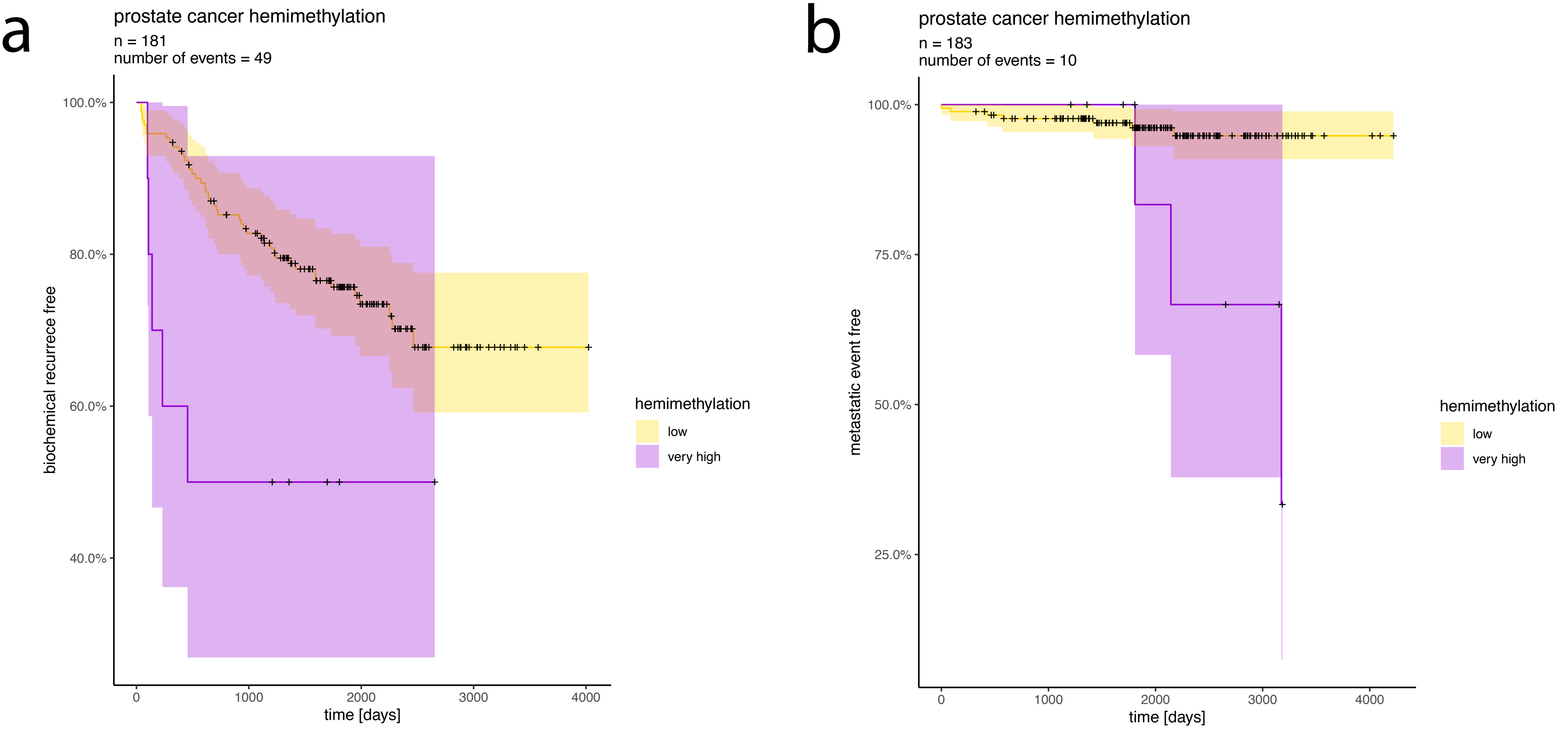
a
prostate cancer hemimethylation
n = 181
number of events = 49
biochemical recurrece free
100.0%
80.0%
60.0%
40.0%
0
1000
2000
3000
4000
time [days]
hemimethylation
low
very high
b
prostate cancer hemimethylation
n = 183
number of events = 10
metastatic event free
100.0%
75.0%
50.0%
25.0%
0
1000
2000
3000
4000
time [days]
hemimethylation
low
very high

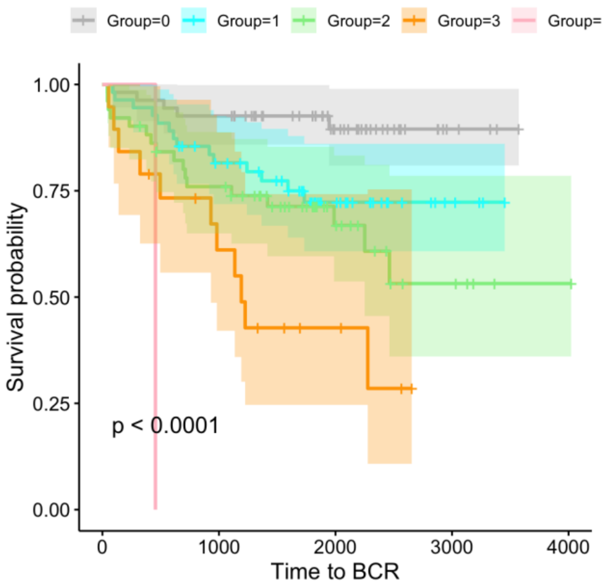


| Model | df | Log-lik |
|---|---|---|
| $\log(T_i) = \alpha + \epsilon$ | 1 | -467.6 |
| $\log(T_i) = \beta_{n_i} + \epsilon$ | 5 | -454.2 |
| $\log(T_i) = \alpha + \beta n_i + \epsilon$ | 2 | -455.1 |

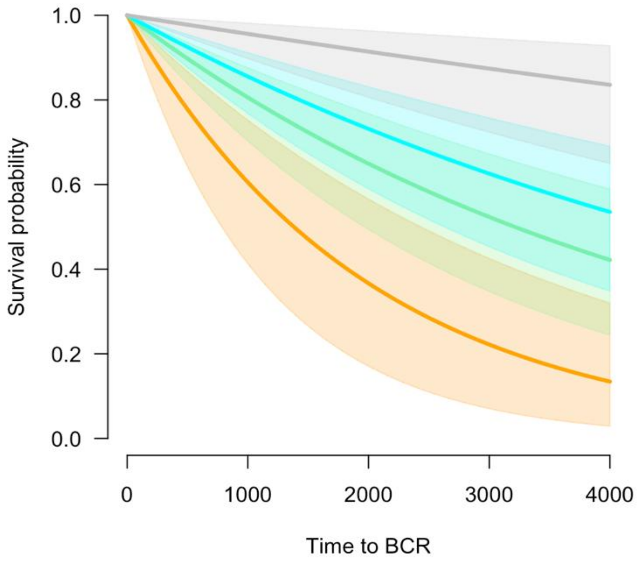


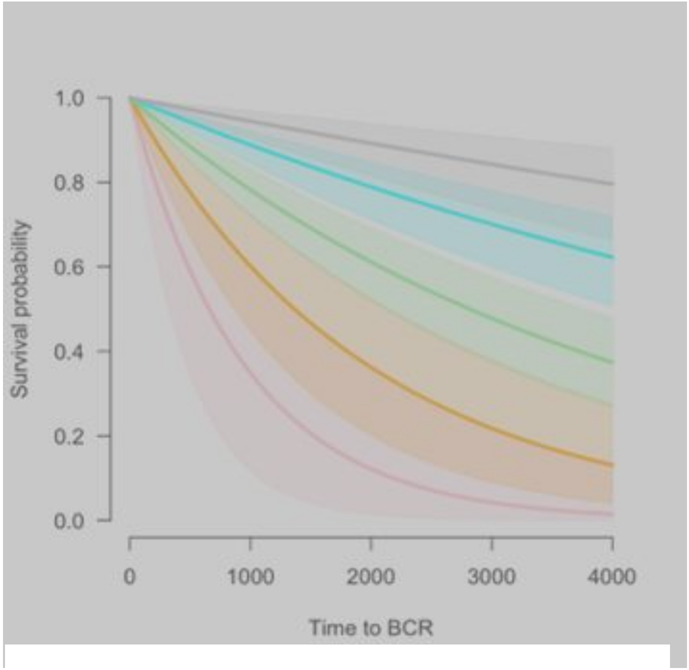


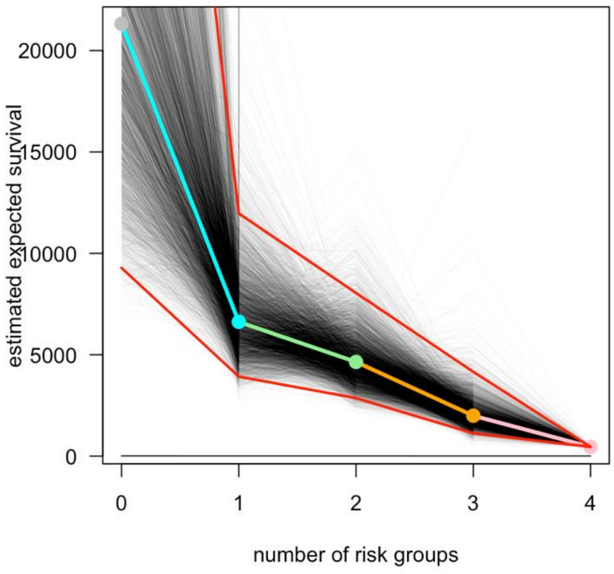


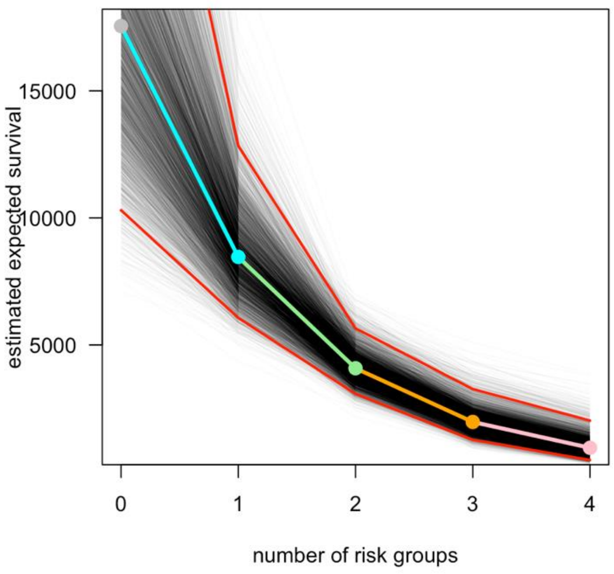

#### Estimate of parameter for number of risk groups in three exponential survival models with different covariates

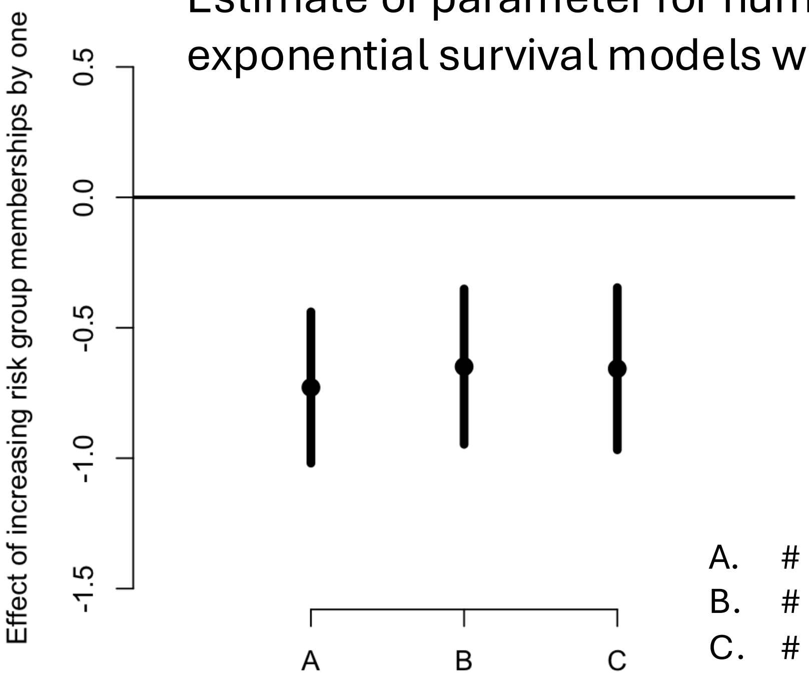


A. # Risk Groups
B. # Risk Groups + Grade Group
C. # Risk Groups + Grade Group + Age

#### Surival estimates for different numbers of risk group memberships if we restrict ourselves to 113 Grade Group 2 cases.

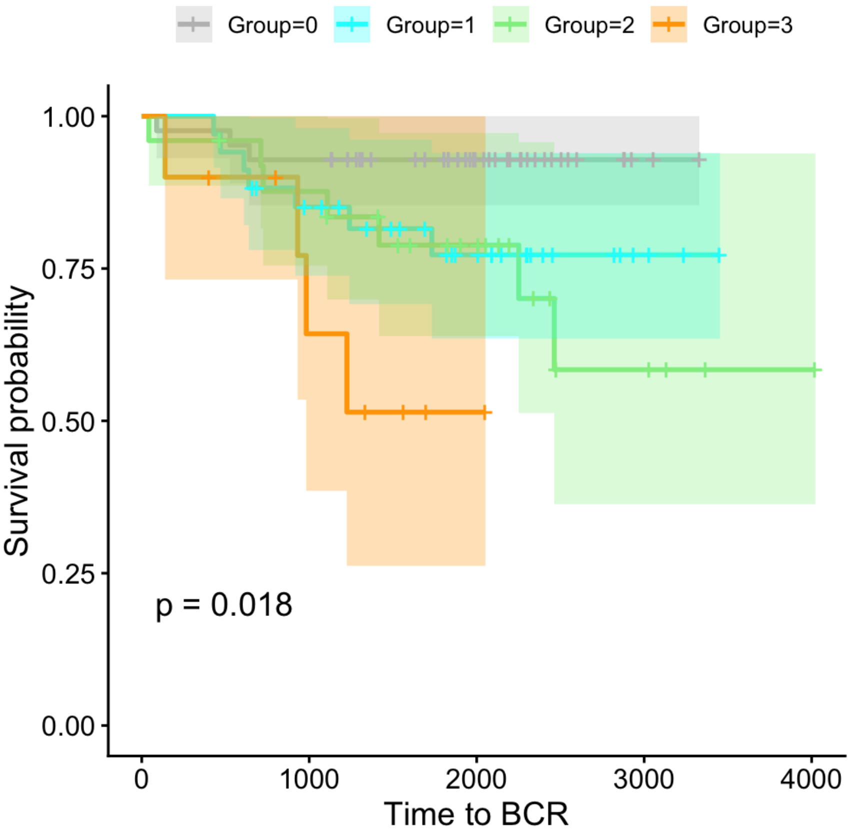

CRUK_PC_0045_Tumour

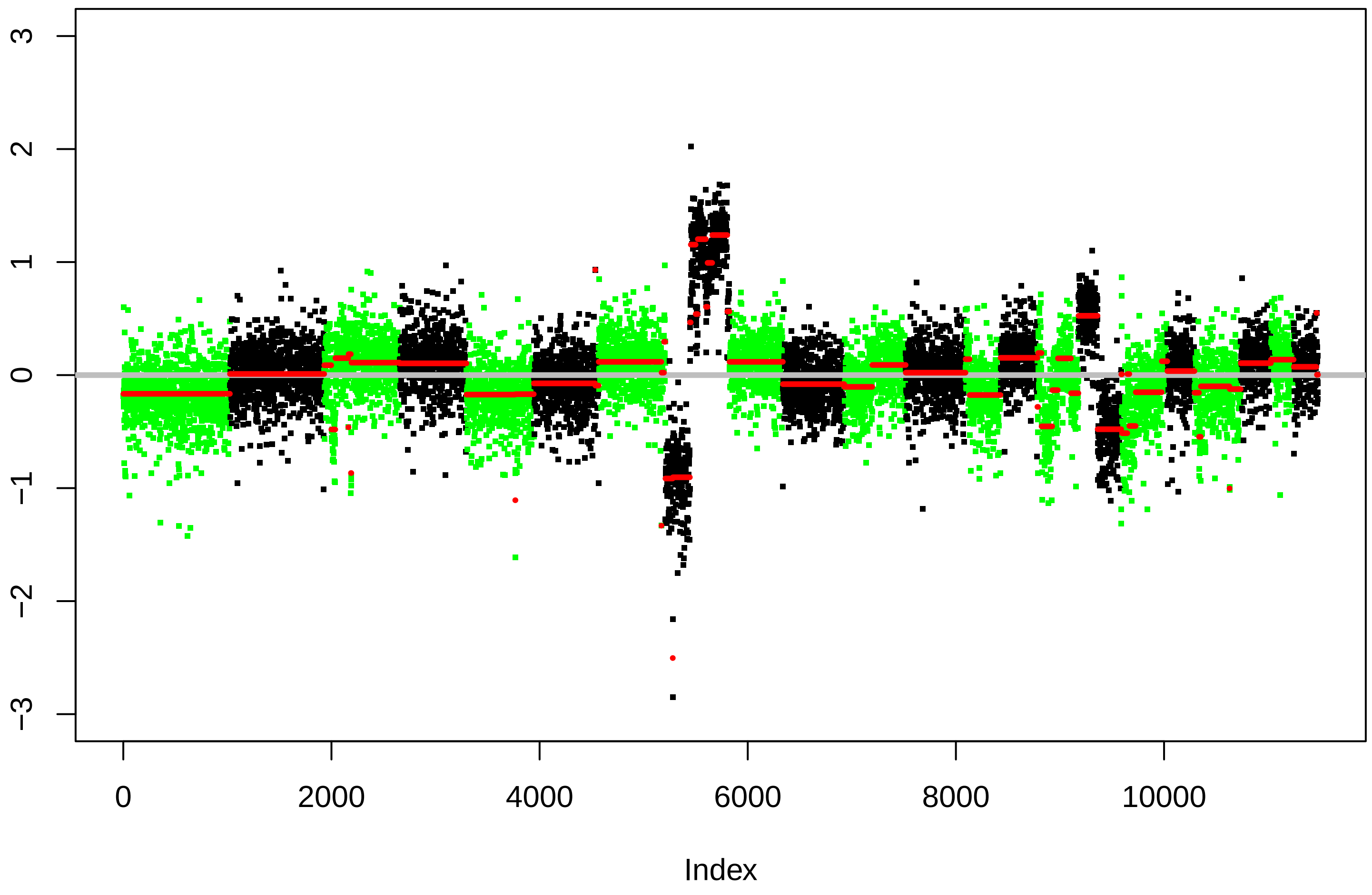


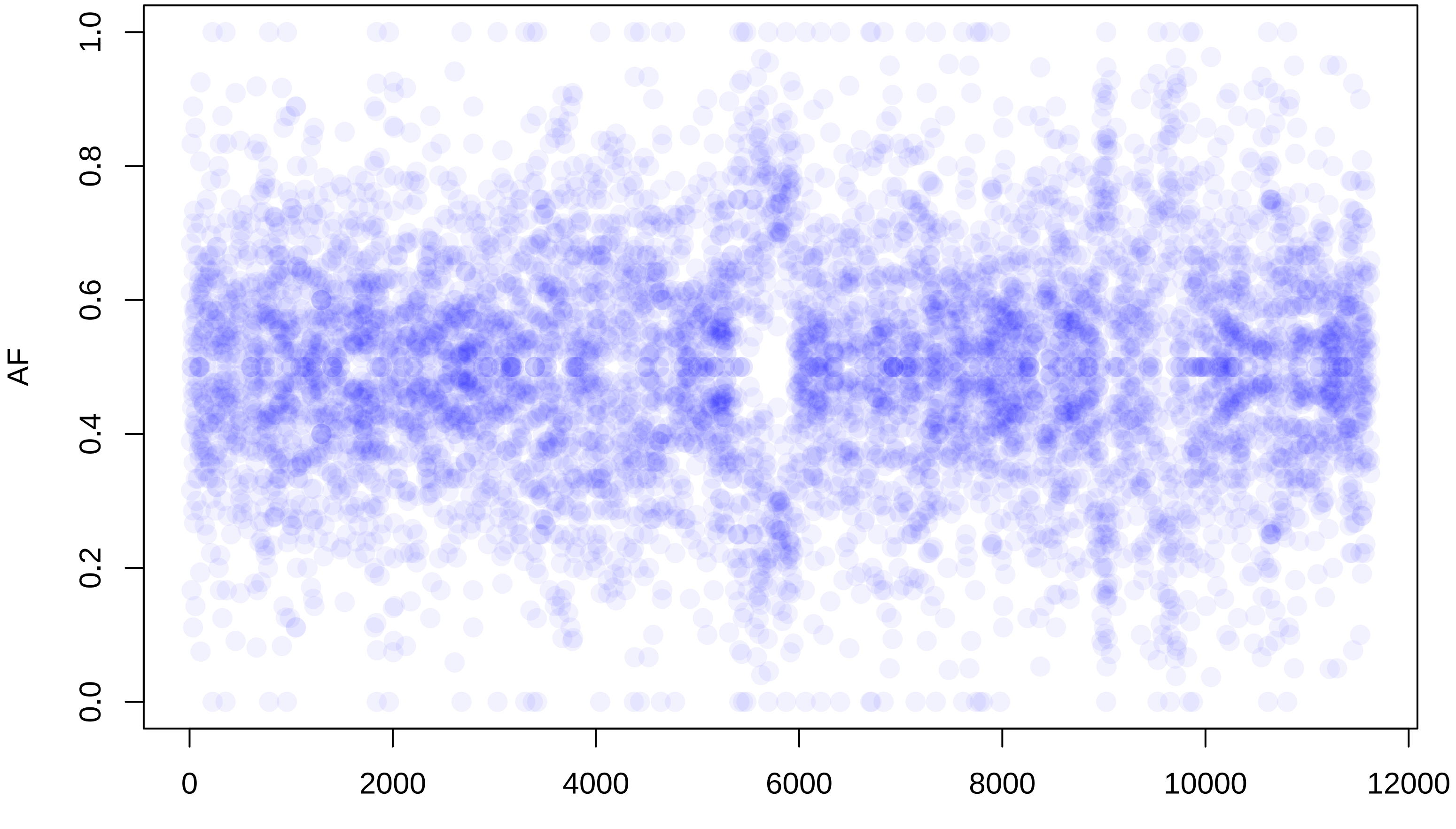

**CRUK_PC_0045_Benign**

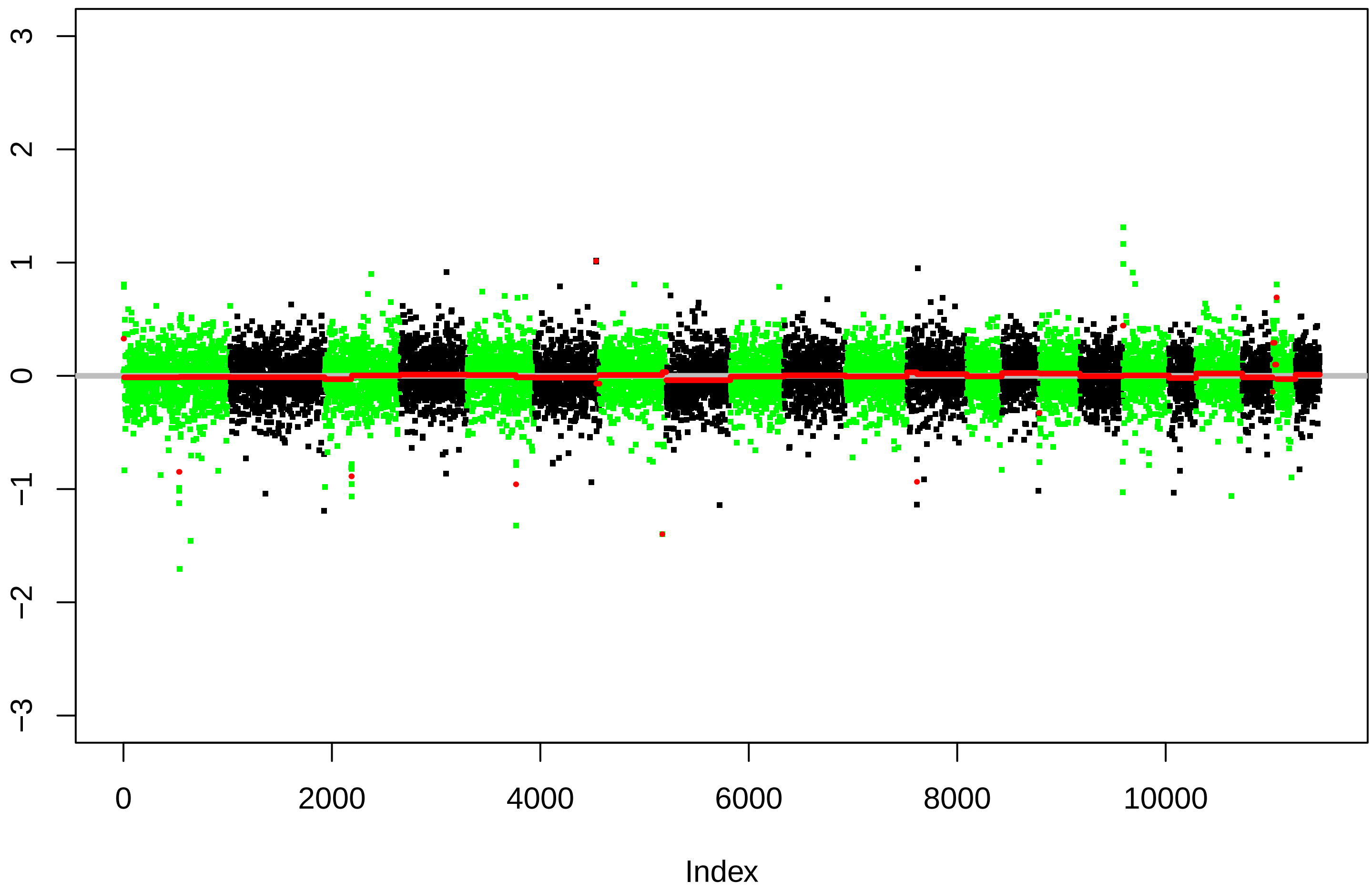


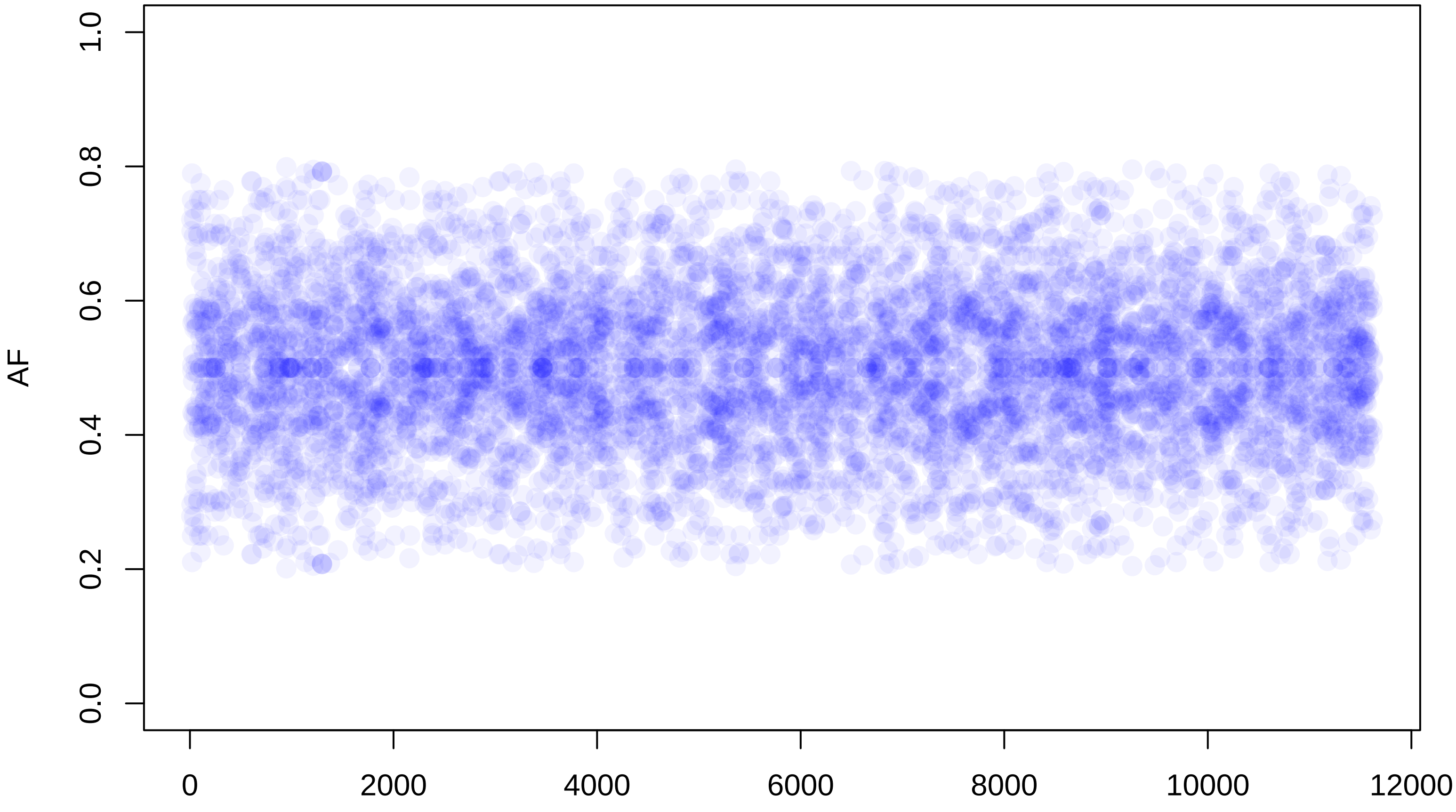

### CRUK_PC_0160_Tumour

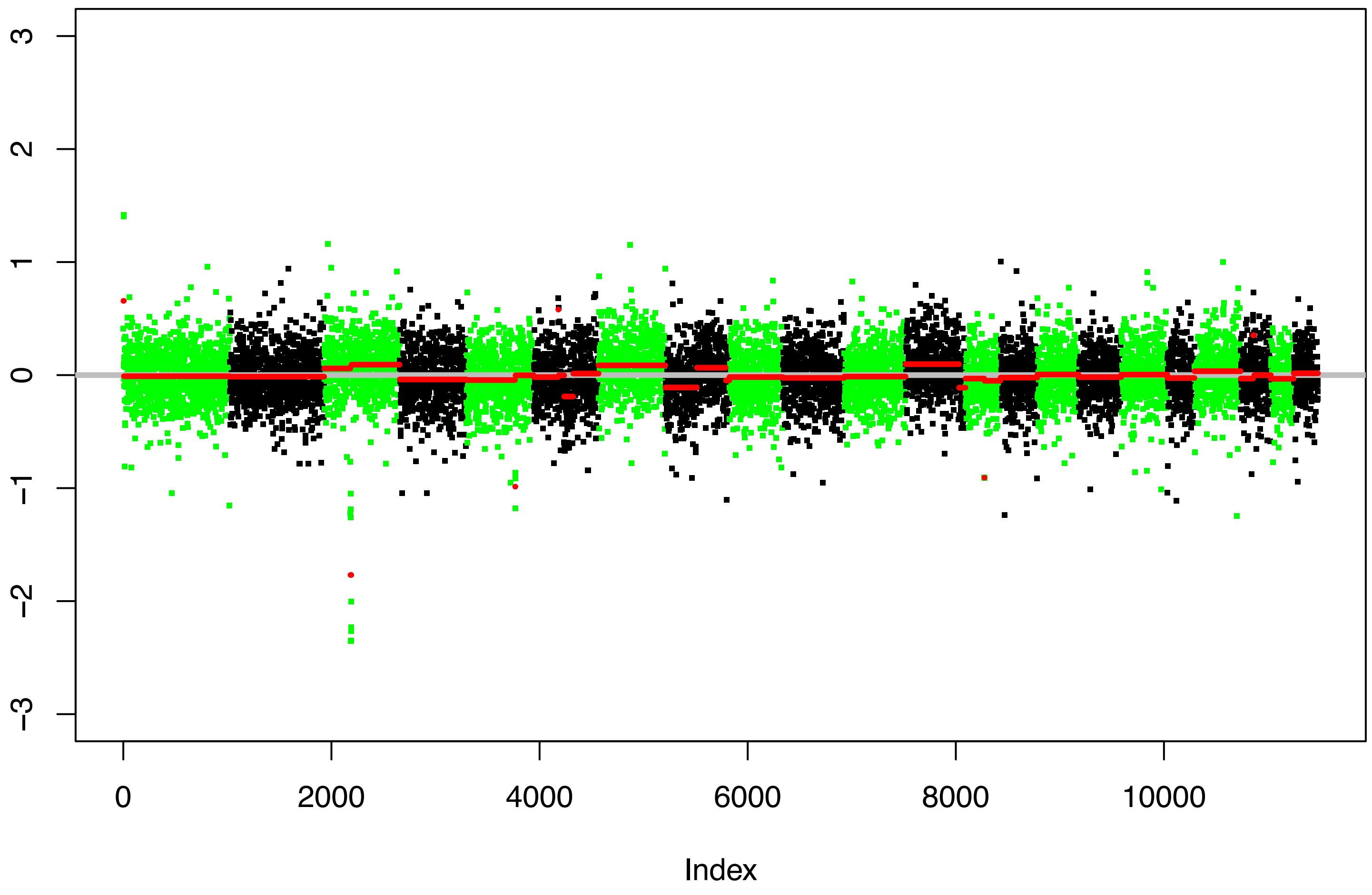


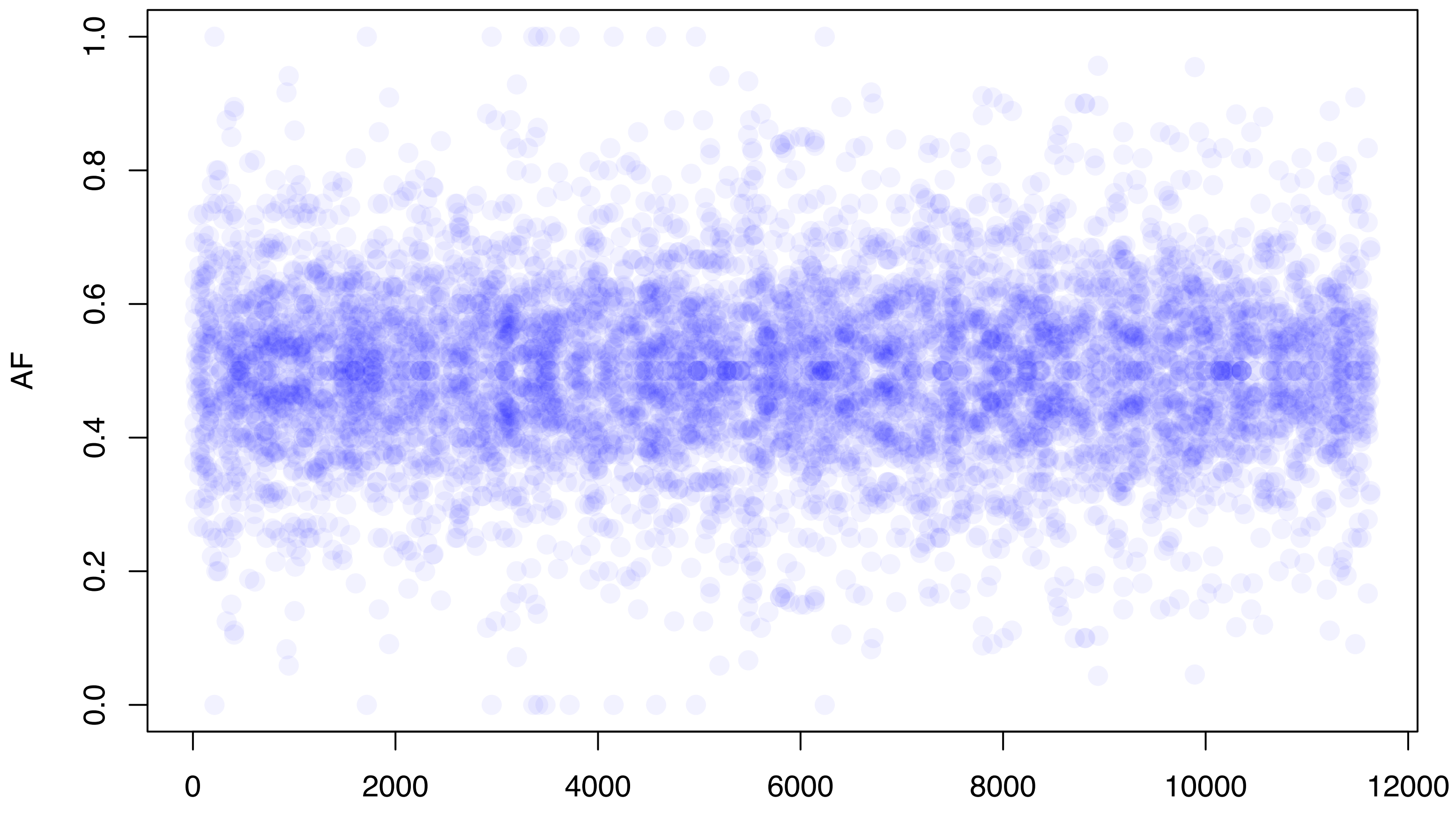

CRUK_PC_0160_Benign

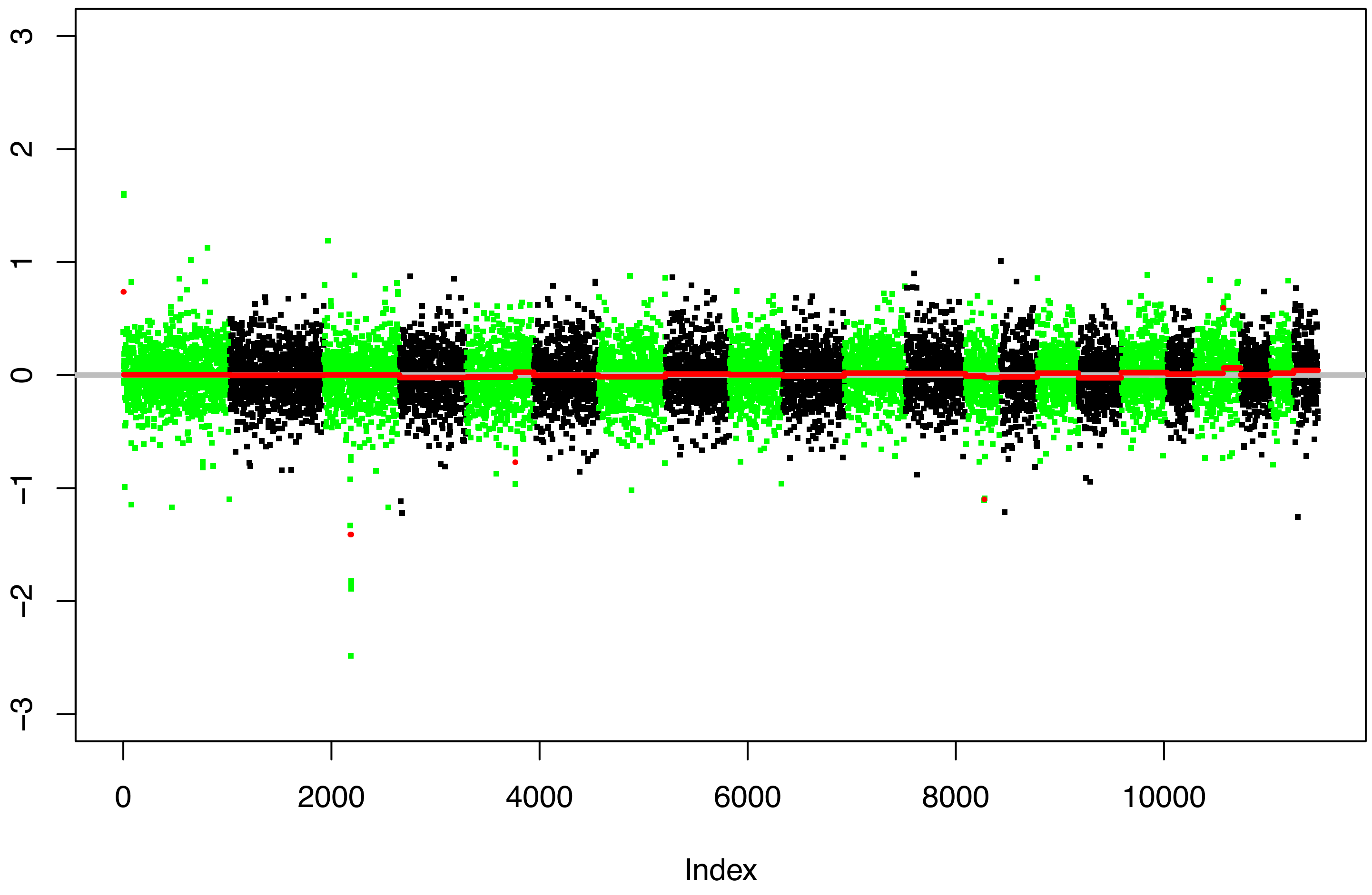


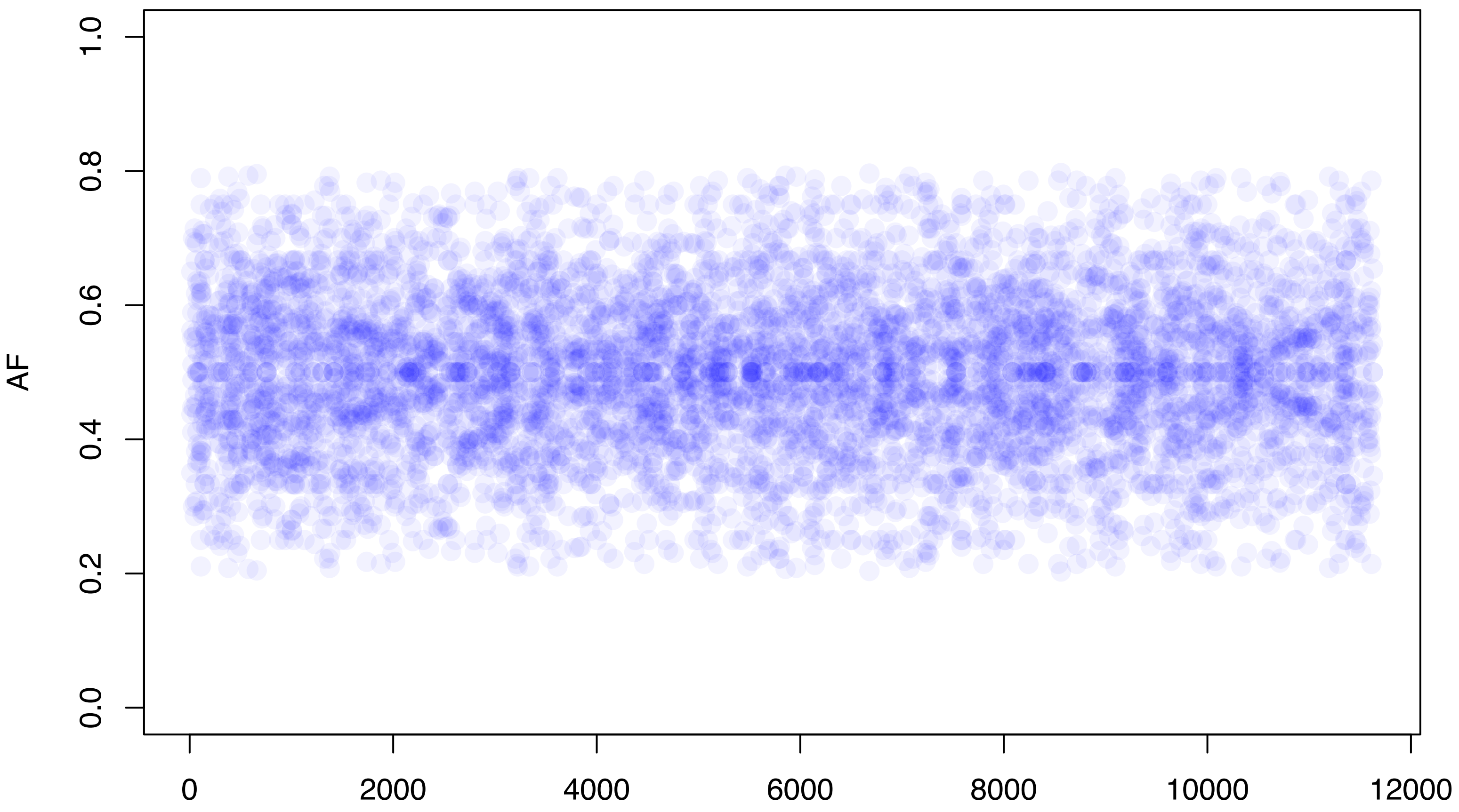

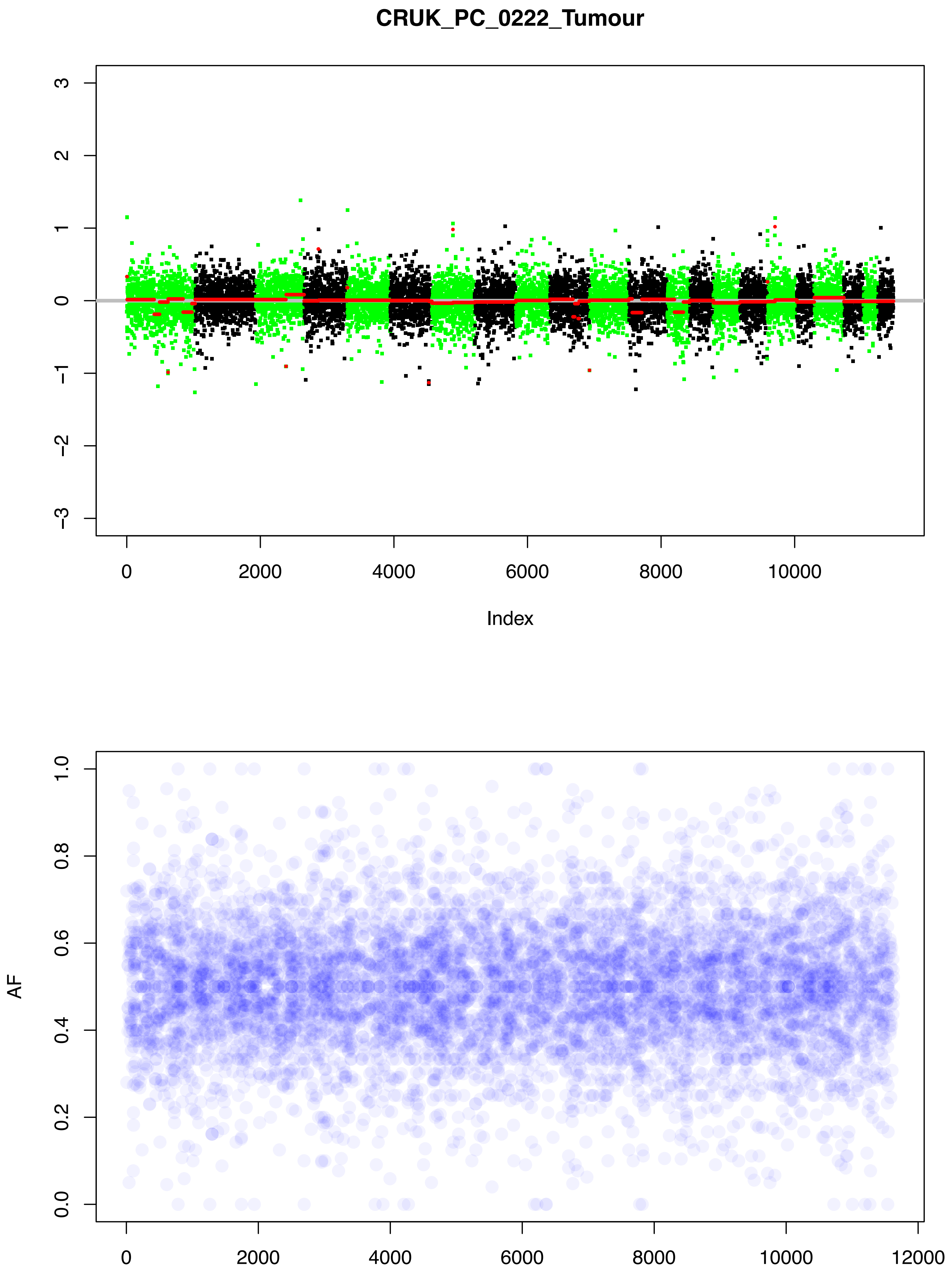
CRUK_PC_0222_Tumour
3
2
1
0
−1
−2
−3
0
2000
4000
6000
8000
10000
Index
1.0
0.8
0.6
0.4
0.2
0.0
AF
0
2000
4000
6000
8000
10000
12000

### CRUK_PC_0222_Benign

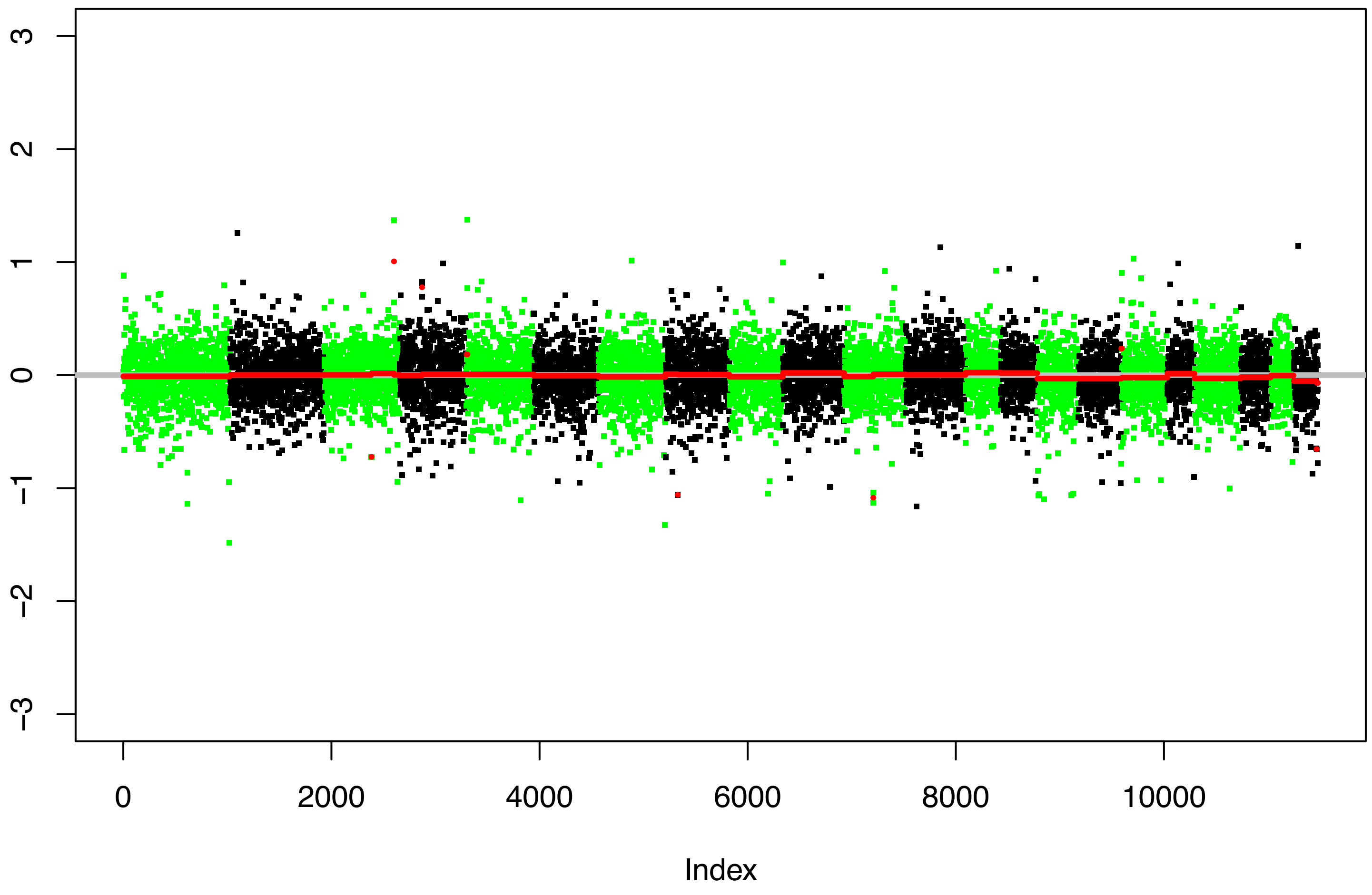


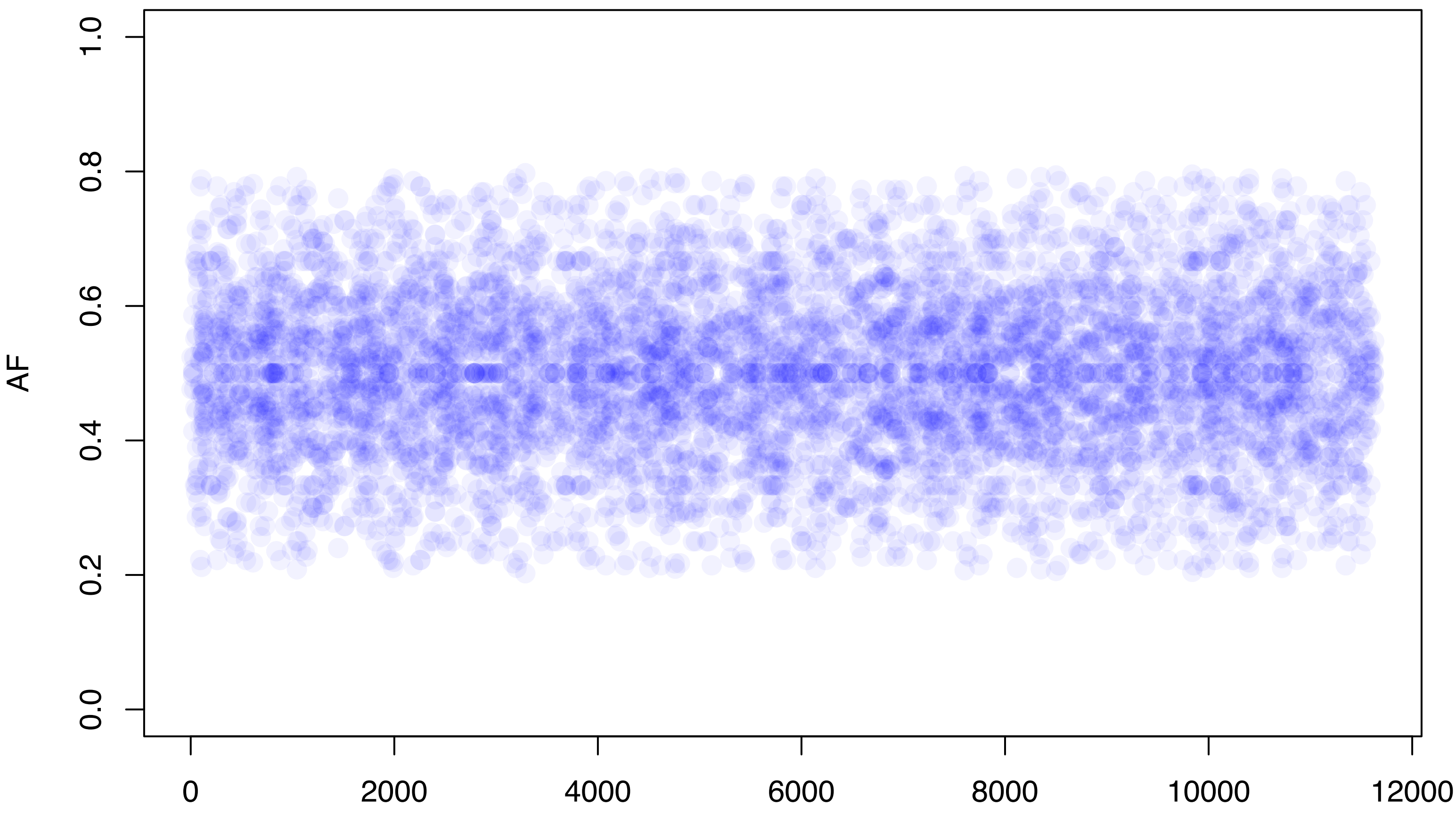

#### CRUK_PC_0202_Tumour

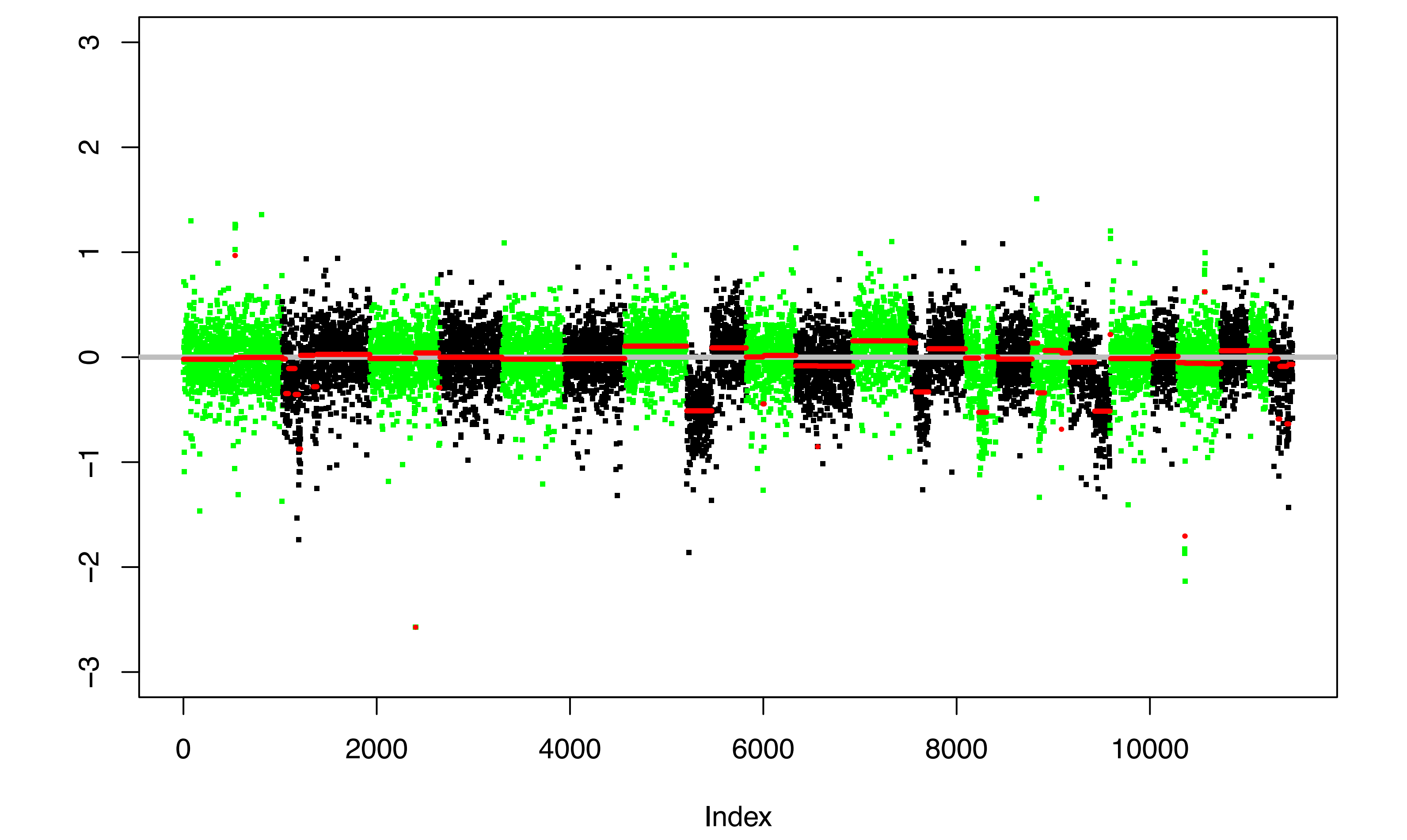


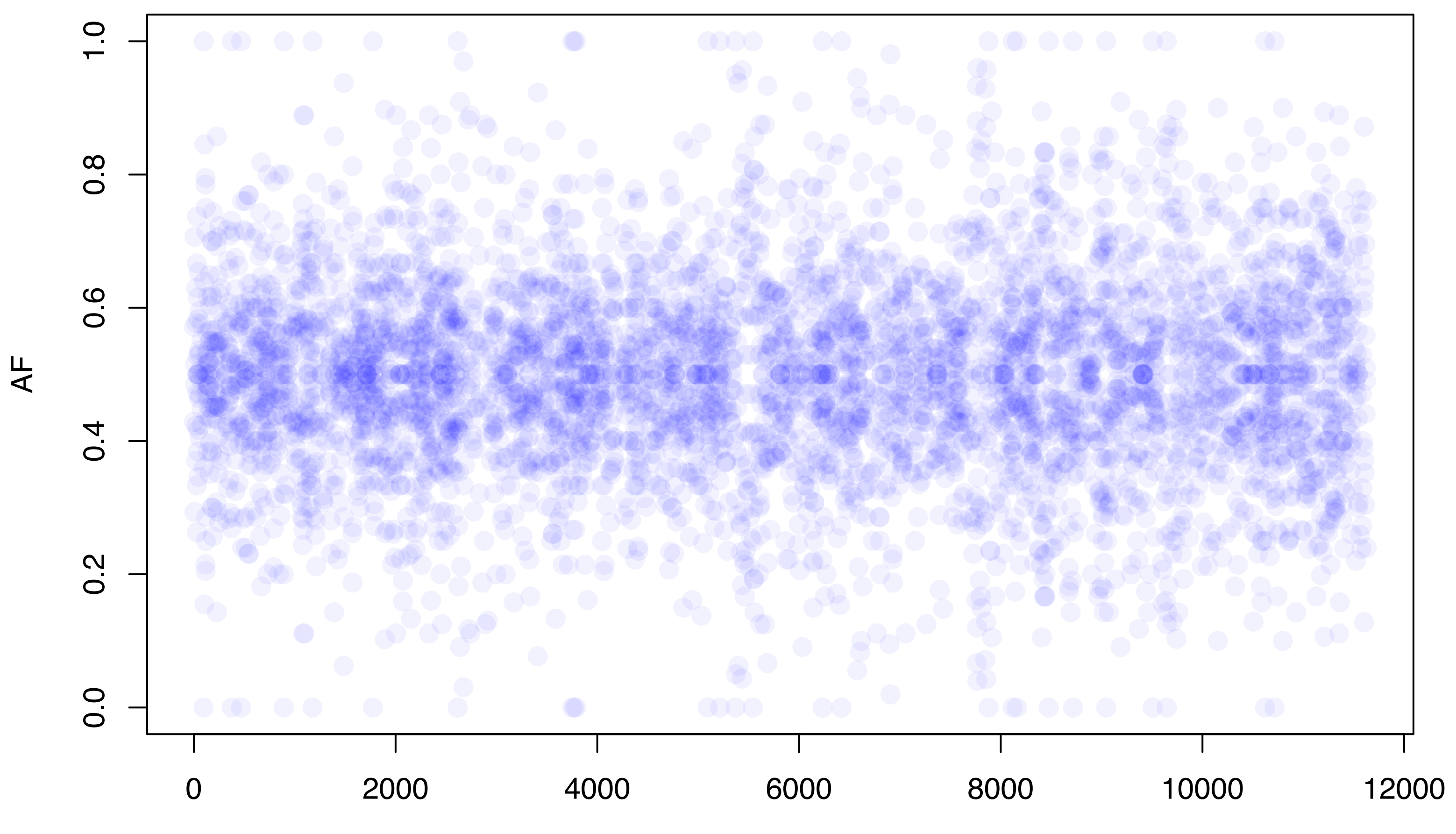

#### CRUK_PC_0202_Benign

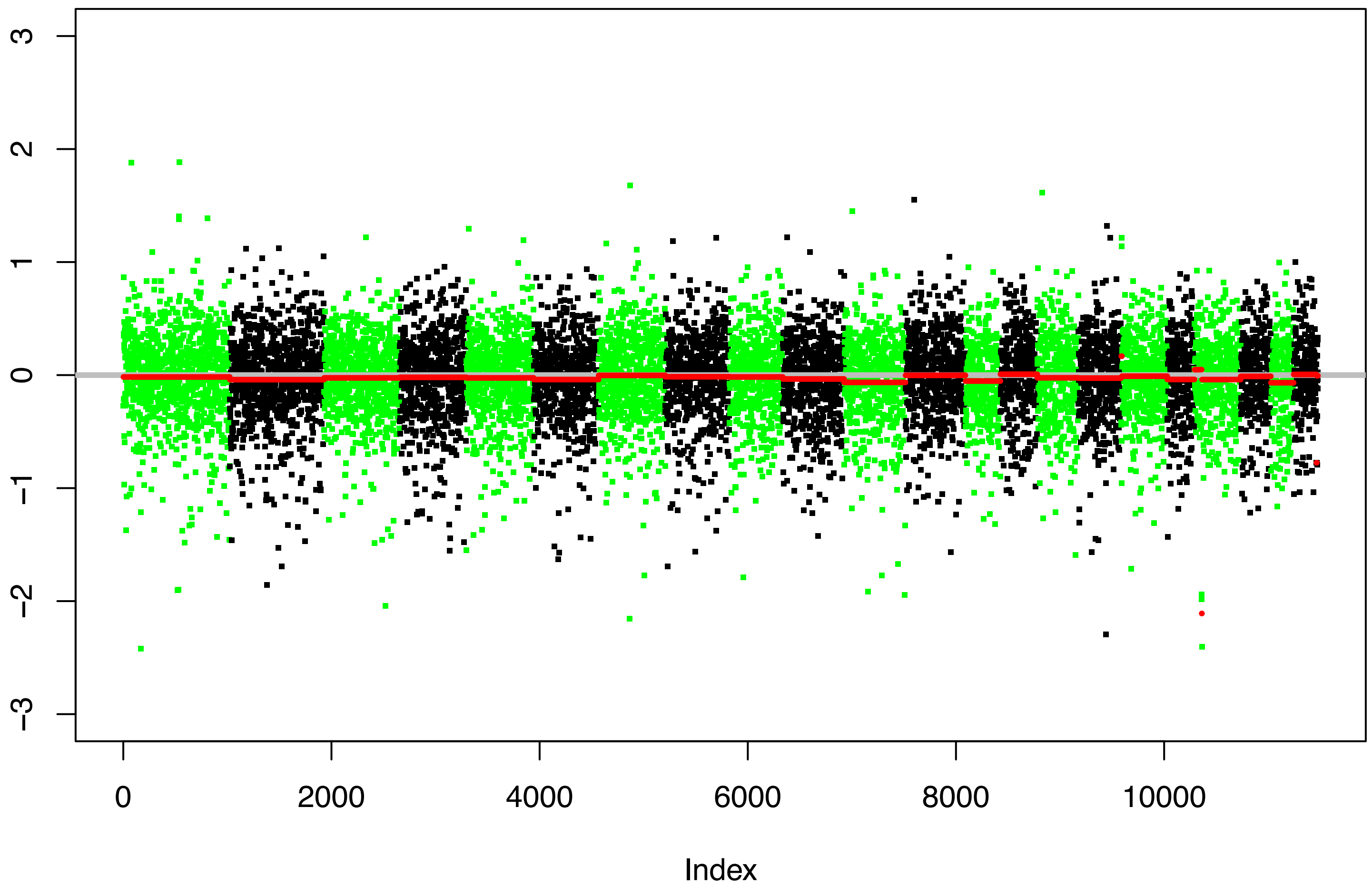


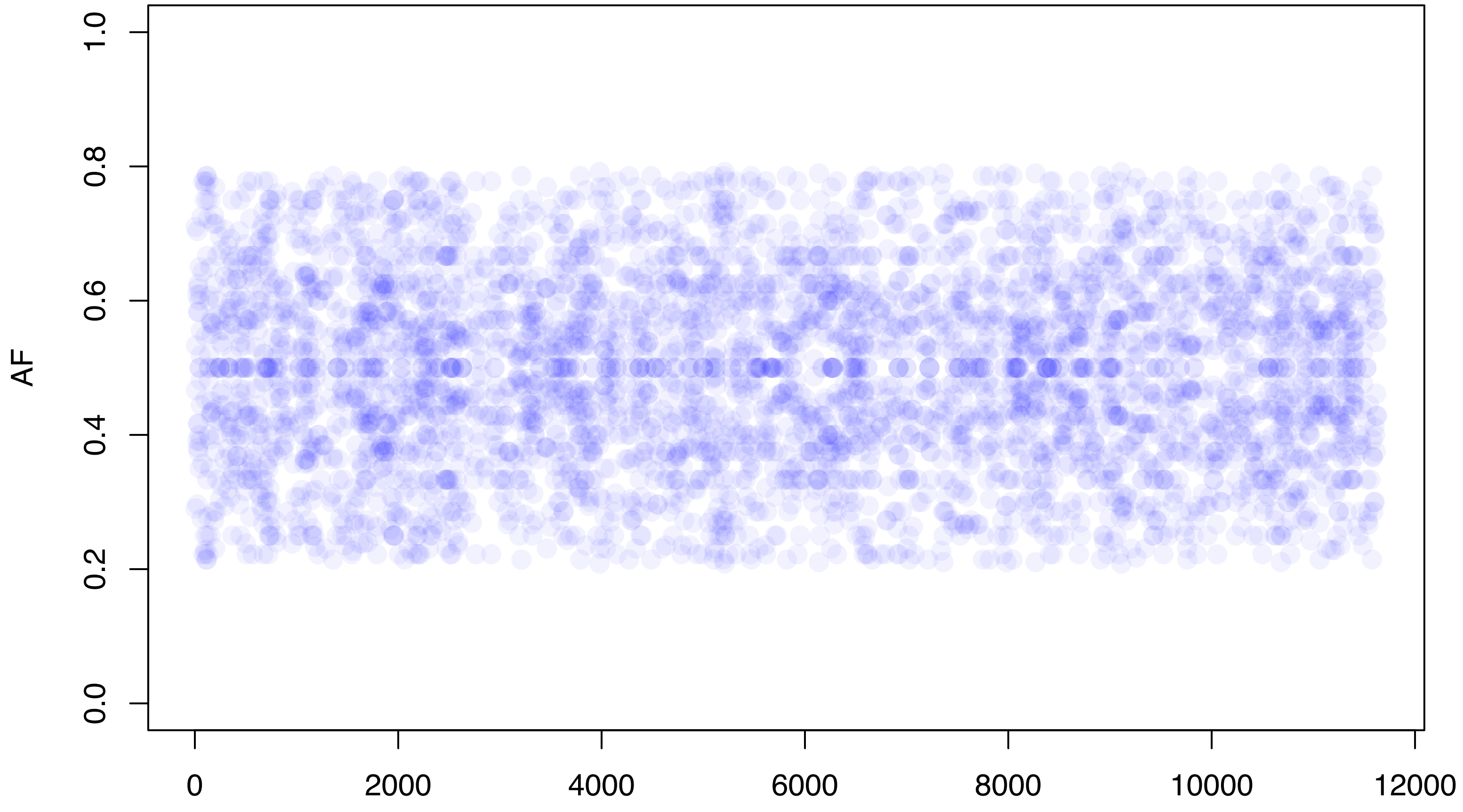
